# The PMK-3 (p38) Mitochondrial Retrograde Response Functions in Intestinal Cells to Extend Life via the ESCRT Machinery

**DOI:** 10.1101/797308

**Authors:** Oxana Radetskaya, Rebecca K. Lane, Troy Friedman, Aria Garrett, Michael Nguyen, Megan B. Borror, Joshua Russell, Shane L. Rea

**Affiliations:** Department of Pathology, University of Washington, Seattle, WA 98195, USA; Department of Microbiology, Immunology, and Molecular Genetics, University of Texas Health at San Antonio, San Antonio, TX 78229, USA; Our Lady of the Lake University, San Antonio, TX 78207, USA

**Keywords:** Aging, ageing, exosomes extracellular vesicles, *isp-1*, Mit mutants, traffic, retrograde response, TBB-6

## Abstract

The p38 mitogen-activated protein kinase (MAPK) PMK-3 controls a life-extending retrograde response in the nematode *Caenorhabditis elegans* that is activated following mitochondrial electron transport chain (ETC) disruption and is distinct from known longevity-promoting pathways. Here we show that the long isoform of PMK-3 expressed exclusively in the gut, rather than neurons, is sufficient to fully extend the life of animals exposed to mild ETC dysfunction. Surprisingly, constitutive activation of PMK-3 using a gain-of-function MAP3K/DLK-1 mutant does not extend the life of wild-type worms due to dampening of the DLK-1/PMK-3 signaling axis with age. We further show that core components of the ESCRT-III machinery, including ISTR-1, CHMP2B (CC01A4.2) and RAB-11.1, are required for life extension following ETC disruption. ESCRT proteins are needed for extracellular vesicle (EV) formation, lysosomal traffic and other functions requiring membrane encapsulation away from the cytoplasm. Together, our findings underscore PMK-3 as a pivotal factor controlling life extension in worms following mitochondrial ETC disruption and illustrate the importance of the endomembrane system to this process. Our findings raise the possibility that EVs may act as intra-organismal signaling vehicles to control aging.

## INTRODUCTION

Mitochondrial dysfunction is implicated to some degree in the etiology of most of the major age-related diseases of the Western World, including cancer, atherosclerosis, Alzheimer’s and Parkinson’s diseases, osteoporosis and obstructive lung disease (reviewed in (1)). Even in the absence of disease, mitochondrial function declines with age (2). The pronounced role of mitochondria in the pathogenesis of multiple diseases is a consequence of their function in an extensive list of essential cellular processes. For example, the mitochondrial electron transport chain (ETC) alone generates up to 90% of cellular ATP (3). Other important functions of mitochondria include calcium sequestration (4), Fe-S cluster formation (5) and nucleotide biosynthesis (6). Due to the central importance of mitochondria, their functional status is closely monitored within cells (7). In this regard, signals called “retrograde responses” originate from compromised mitochondria and act within the nucleus to coordinate adaptive gene expression changes to resolve or reduce mitochondrial stress. Higher eucaryotes have evolved different kinds of retrograde responses that are triggered by a variety of stressors, including protein aggregation (8), changes in ETC activity (9), oxidative stress (10) and loss of iron (11). One innovative approach to countering age-related decline of mitochondrial function is to exploit retrograde responses to prophylactically protect, or even rejuvenate, the mitochondrial network. Retrograde responses might therefore represent an untapped means by which to potentially ameliorate multiple age-related diseases simultaneously.

Responses that counteract mitochondrial ETC disruption have been most extensively studied in the nematode *Caenorhabditis elegans* (9, 12–16). Several years ago, we showed that the response of *C. elegans* to mitochondrial ETC dysfunction is threshold-dependent. That is, low levels of stress produce no phenotype, moderate levels increase lifespan, while severe disruption, as in humans, leads to overt pathology and shortened lifespan (13). Recently, we described a novel retrograde response in *C. elegans* that extended life and did not overlap with known longevity-promoting pathways (17). This retrograde response operates independently of the well-characterized ATFS-1-dependent mitochondrial unfolded protein response (18), the NRF2/SKN-1-dependent anti-oxidant response (19), as well as a variety of other tested pathways. The core of this signaling pathway consists of a new mitogen-activated protein kinase (MAPK) cascade composed of DLK-1 (MAP3K), SEK-3 (MAP2K), and PMK-3 (p38 MAPK). Each of these are needed for life extension following exposure to mild ETC disruption. Little or nothing is known regarding in which tissues these molecules promote longevity, or the mechanisms by which they operate other than that an unconventional β-tubulin, TBB-6, becomes 70-fold overexpressed when the pathway responds to mitochondrial ETC disruption (17). In this study, we leveraged this sensitive physiological indicator to define the tissues and molecular mechanisms by which this novel MAPK signaling cascade functions and identify a new role for the membrane remodeling ESCRT-III machinery in promoting ETC stress-induced longevity.

## METHODS

### Nematode Strains and Maintenance

A complete list of the 36 *C. elegans* strains employed in this study is provided in **Table S1**. Strains were maintained at 20ºC on standard NGM-agar plates (20).

### Transgene Construction and Transgenic Strain Generation

21 strains were constructed for this study. Detailed methods of recombinant array construction and microinjection procedures are provided in **S1 Text**.

### Feeding RNAi

Bacterial feeding RNAi constructs were purchased when available or generated in-house. Full details of clone construction and purchase IDs are provided in **S1 Text**.

### Western Analyses

Worm extracts were prepared as previously described using either 1% SDS detergent or 1% Triton-X-100 (Tx-100) detergent (21). For western blotting, samples were separated on 4-12% SDS-PAGE gels, transferred to nitrocellulose, and PMK-3 detected using an in-house generated α-PMK-3 primary antibody (Rabbit #75, 70-day bleed) in conjunction with chemiluminescence.

### Antibody Generation and Testing

Rabbit antisera directed against amino acids 438-455 of PMK-3(L) was generated under contract by Life Technologies Corporation (ThermoFisher Scientific). Antibody production details, as well as sensitivity- and specificity testing, are described in **S1 Text**.

### Homology Modeling

A model of PMK-3(L) was built via the CPHmodels-3.3(beta) homology modeling server. Human p38α (PDB structure 1R3C) served as the model-building template. Only amino acids 110-459 of PMK-3(L) were included in the final model. Full details are provided in **S1 Text**.

### *In silico* Data Mining

Strongly conserved genes in *C. elegans* that contained promoter motifs that were also strongly conserved and which matched Motif 2 from Fig. 1C in (17) were identified using the *C. elegans* v4 cisRED database (22). For screening, this sequence was approximated using 5’-RTTKYGAAAY-3’, where R = G or A, K = G or T, Y = T or C and both nucleotide choices were weighted equally. This analysis identified 206 elements spread over 188 genes. The DAVID Bioinformatics toolbox (v6.8) (23) was used to identify cellular pathways in which these genes functioned. Pathway maps were generated using KEGG (24). Full details are provided in **S1 Text**.

**Figure 1.**
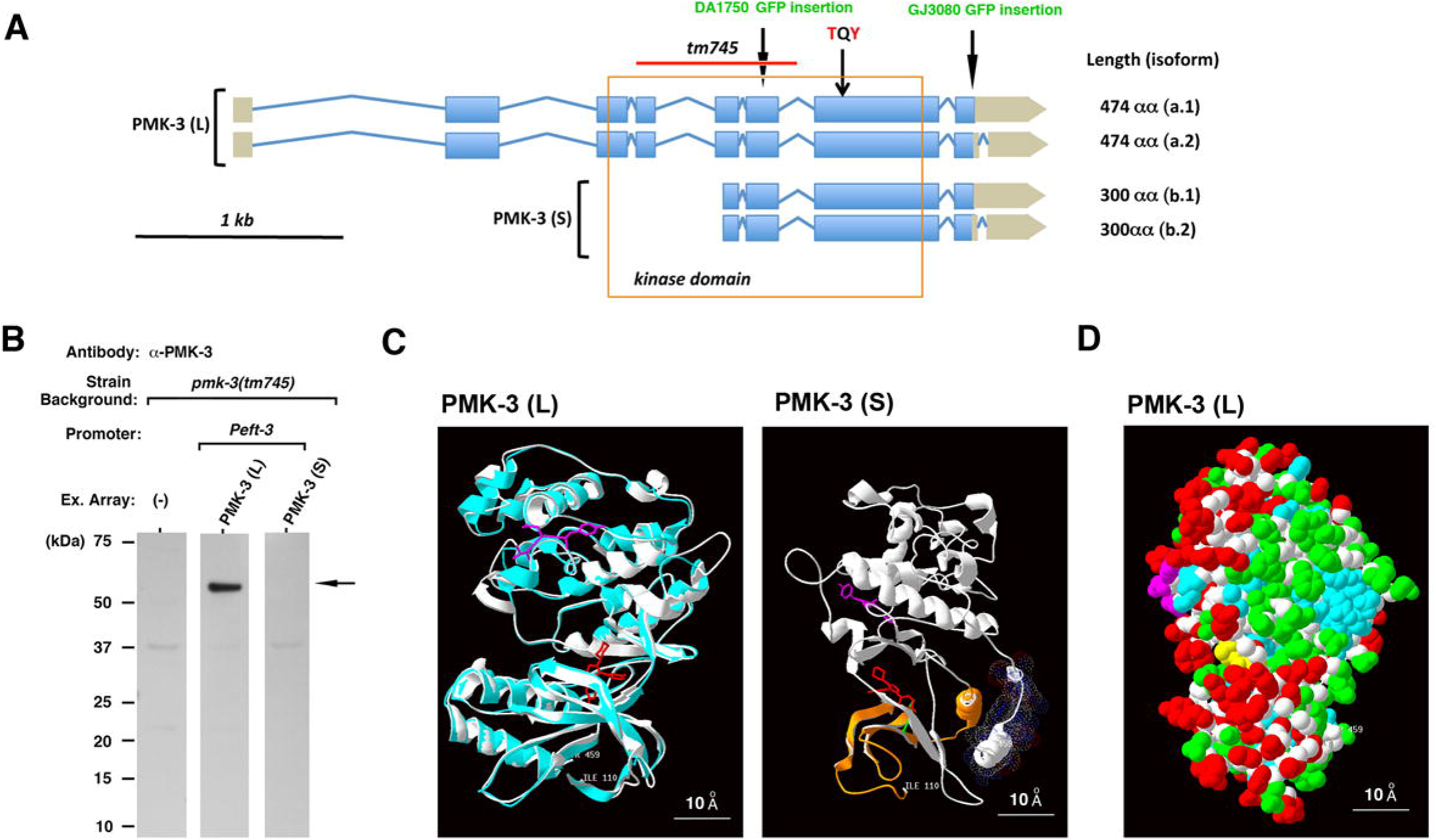
Only the long isoform of PMK-3 encodes a stable protein. (**A**) Schematic of *pmk-3* genetic locus. mRNA splice variants are shown. Size of encoded protein products listed on *right* (αα - amino acids). Genomic DNA deleted in *pmk-3(tm745)* shown with *red* bar. Location of amino acids altered by point mutation in this study are colored *red*. Exons encoding relevant protein domains are boxed. Position of the GFP fusion junction in the PMK-3 translational reporters of strains DA1750 and GJ3080 are marked (*long arrows*). (**B**) Western analysis of *pmk-3(tm745)* worms containing the long (L) or short (S) recombinant PMK-3 isoforms under control of the ubiquitously-expressed *eft-3* promoter. All lanes contain equal amounts of whole-worm lysate (1% SDS). Arrow marks PMK-3(L). Parent blot shown in **Fig. S4**. (**C**) *(Left panel)* Homology model of PMK-3 (αα 110-459) (*white)* based on crystal structure of human p38α (PDB: 1R3C) (*cyan*). TQY of the PMK-3 activation loop is shown (*magenta*), along with the predicted position of SB220025 inhibitor binding (*red*) marking the ATP-binding pocket. A structure-based alignment of both proteins is provided in **Fig. S5**. *(Right panel)* PMK-3(S) lacks the first 174 αα of PMK-3(L) (residues highlighted in *orange)*, constituting half of the C-terminal core domain and including key ATP-stabilizing residues such as LYS150 (*green*) conserved in other MAPKs. TQY of the PMK-activation loop is shown (*magenta*), along with the location of the C-terminal peptide epitope used for production of PMK-3 antisera (*dotted surface*). Disruption of the predicted ATP binding site of PMK-3(L), marked by SB220025 *(red)*, is evident. Image rotated 90° relative to *left panel*. (**D**) Space-filling model of PMK-3(L) showing asymmetry of non-conserved residues (*red*) relative to both PMK-1 and PMK-2. (Additional coloring: *white:* C_α_ carbon; *cyan:* identical residue, *green:* conservative alteration: *yellow:* SB220025, *magenta:* TQY).

### Microarray Analysis of Endocytosis Gene Set

We used the GEO profiles database (25) to analyze published microarray data (GEO#: GSE38196) for changes in mRNA expression of nine endocytosis genes of interest: *alx-1, C01A2.4, istr-1, rab-11.1, rfip-1, rme-1, usp-50, vps-2, vps-32.1*. Details regarding sample preparation and statistical testing are provided in **S1 Text**.

### qRT-PCR Analysis of Endocytosis Gene Set

cDNA was prepared from one-day old adult worms cultured on relevant RNAi plates from the L1 larval stage onward. qPCR was performed using SYBR Green and a StepOnePlus Real Time PCR System (Thermo Fisher Scientific). Changes in mRNA expression of endocytosis-related genes was calculated using the ΔΔC_t_ method and *pmp-3* as housekeeping gene control. Statistical significance was assessed using a one-tailed Student’s t-test. Full details of our qRT-PCR procedure, including all primer sequences, is provided in **S1 Text**.

### Extracellular Vesicle (EV) Isolation

EV-containing fractions from adult worms was collected using size exclusion chromatography as described (26). Samples were utilized within a 72 hour window. Detailed isolation methods are provided in **S1 Text**.

### Lifespan Analyses

Lifespan studies were performed as described previously (13). Use of FudR was avoided. The first day of adulthood was designated as day one in all experiments. Descriptive statistics and significance testing for all lifespan studies are provided in Table S2. Data was analyzed using OASIS 2 (27).

### Fluorescence Imaging and Quantification

Worm images were captured using an Olympus DP71 CCD camera connected to an Olympus SZX12 or SZX16 fluorescence dissecting microscope. Image J (NIH) was used for quantification. See **S1 Text** for full details.

## RESULTS

As PMK-3 is the most distal kinase of the new MAPK cascade controlling ETC dysfunction-induced longevity in *C. elegans*, we sought to understand its role in greater detail. The *pmk-3* gene is predicted to encode four mRNA splice variants generating two protein isoforms, 300 and 474 amino acids (αα) in length (http://www.wormbase.org, WS272, Sep. 2019, **Fig. 1A**). Hereafter, we refer to these two protein isoforms as PMK-3 short (S) and PMK-3 long (L), respectively. The evidence which supports existence of transcripts encoding PMK-3(S) is weak and is derived from a single expressed sequence tag (www.wormbase.org, *yk1033d01*) that was computationally extended upstream to the most proximal 5’ ATG.

To test which isoform is required for mediating retrograde response signaling, we began by generating recombinant clones of each isoform under the control of the ubiquitous *eft-3* promoter and microinjected them into PMK-3 deficient *pmk-3(tm745)* worms. Previous studies have reported *pmk-3* is ubiquitously expressed (28); the *eft-3* promoter thus provides a control for the maximum degree of rescue mediated via expression from a transgenic array. As a PMK-3 antibody is not commercially available, we generated antisera to PMK-3 for testing expression of each transgene (**S1 Text**, see also **Figs. S1-S3**). Western analysis of whole-worm extracts revealed that while the long isoform of PMK-3 was robustly expressed, the short isoform could not be reliably detected (**Figs. 1B**, **S4A** and **S4B**).

### Structural modeling suggests PMK-3(S) is likely a computational artifact

To further explore why PMK-3(S) remained undetectable even when overexpressed using a constitutive promoter, we turned to structural modeling. p38 MAPKs are comprised of two domains: an N-terminal domain composed largely of β-sheet (~ 135 residues) and a C-terminal domain that is mostly helical (~225 residues). The catalytic site is formed by the juncture of the two domains (29). Homology modeling of PMK-3(L) using the crystal structure of human p38α showed that the PMK-3(S) truncation removed half of the N-terminal core domain, including residues critical for ATP chelation (**Fig. 1C**). Such a severe truncation is unlikely to result in a catalytically functional protein and, given our inability to detect this protein, we believe PMK-3(S) is most likely a computationally inaccurate prediction of Wormbase. We therefore used the long isoform for all further experiments.

Structural modeling of PMK-3(L) also revealed a striking additional feature of this protein that is not immediately apparent from its primary sequence: non-conservative amino acid substitutions that distinguish PMK-3 from both PMK-1 and PMK-2 (**Fig. S5**) decorate half its surface (*red*, **Fig. 1D**). In other words, while the internal kinase core of PMK-3(L) remains almost identical to that of its two paralogs, PMK-3’s surface has evolved new textures that presumably correspond to binding interfaces for unique binding partners, such as upstream kinases and factors that control its subcellular localization. This finding underscores the uniqueness of PMK-3 and is presumably the reason why PMK-3 alone, and not any other MAPK in worms (17), controls the novel longevity-promoting retrograde response that we are currently studying.

### Tissue-specific expression of PMK-3 results in retrograde response activation in gut cells

To investigate which tissue the DLK-1/SEK-3/PMK-3 signaling cascade acts in, we expressed PMK-3(L) under the control of either a ubiquitous (*eft-3*), gut-specific (*vha-*6), or neuron-specific (*rgef-*) promoter. We have previously reported that GFP coupled to the promoter of *tbb-6* (*Ptbb-*6∷GFP) functions as a robust surrogate marker of PMK-3-dependent retrograde response activation following disruption of the mitochondrial ETC. This reporter is strongly induced in gut cells, while weaker expression is also observed in several unidentified head neurons (17). The tissue-specific PMK-3 constructs were therefore injected into *pmk-3(tm745);Ptbb-6*∷GFP worms to evaluate their capacity to activate GFP expression. All constructs contained a downstream SL2 trans-spliced mCherry reporter for validation of transgene expression. Transgenes were maintained as extrachromosomal arrays and siblings that lost the array served as negative controls. Relevant strain names used throughout this study and their full genotypic information is provided in **Table S1**.

All constructs induced robust mCherry expression in the appropriate tissue in accord with its respective promoter (**Fig. 2A-C**). After verifying transgene expression in worms by western blot (**Fig. S4A,B**), the ability of each transgene to allow retrograde signaling in response to mitochondrial stress was evaluated. ETC stress was imposed through bacterial feeding RNAi targeting complex III (*isp-1)* (13, 16, 17). Expression of PMK-3 under all three promoters permitted robust *Ptbb-6*∷GFP induction (**Fig. 2A-C**).

**Figure 2.**
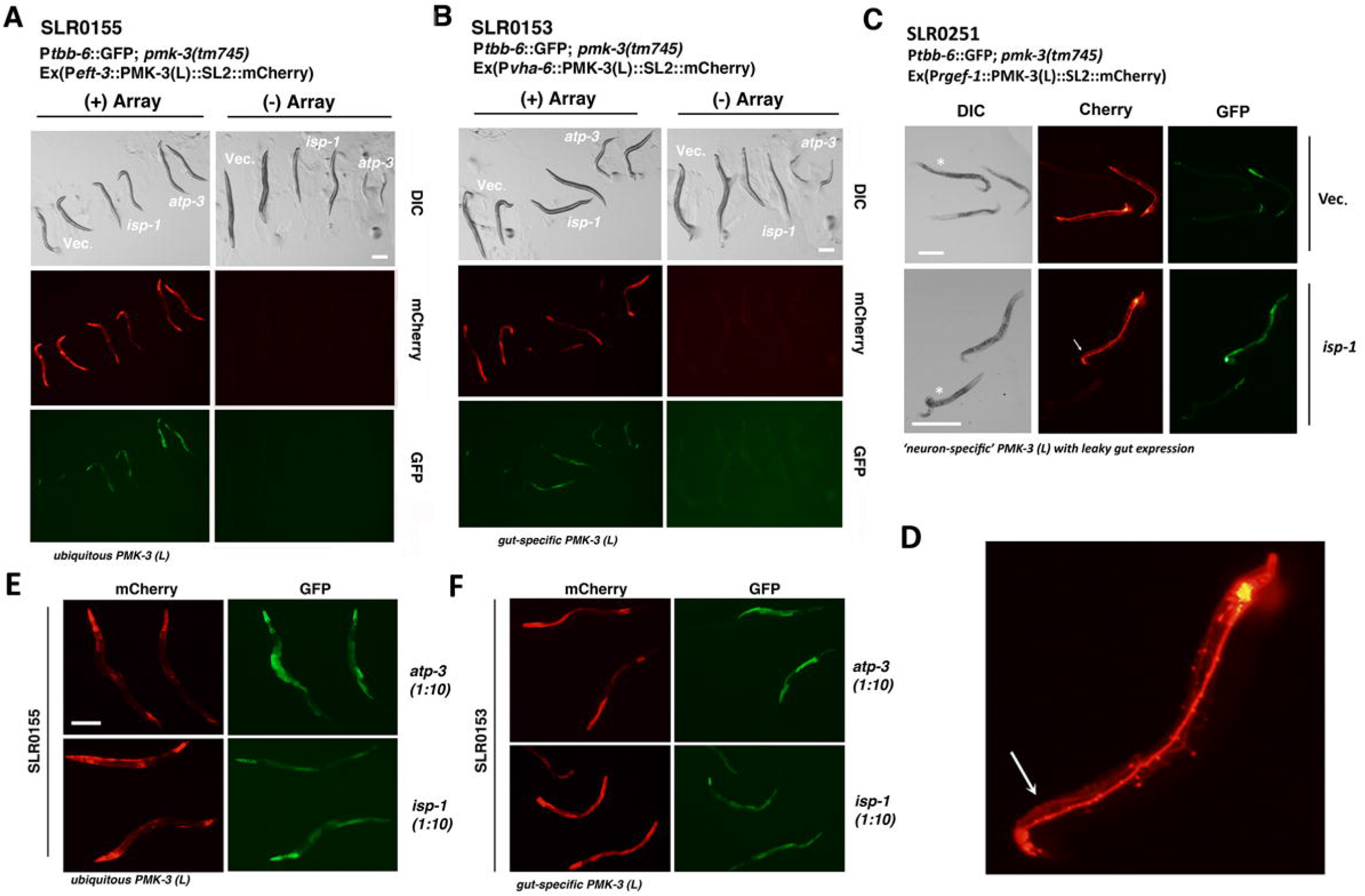
Effect of transgenic PMK-3(L) overexpression on retrograde response activation. (**A-C**) Intestinal expression of P*tbb-6*∷GFP following mitochondrial ETC disruption by *atp-3* or *isp-1* feeding RNAi is restored in *pmk-3(tm745)* mutants expressing ubiquitous (*eft-3* promoter), gut-specific (*vha-*6 promoter) or neuronal (*rgef-1* promoter) PMK-3(L), as marked by mCherry expression. No transgenic construct activates P*tbb-6*∷GFP in the absence of ETC disruption (vector-only treatment, *Vec*.). Worms that have lost their respective transgenic array [right columns in (A) and (B), and worms marked with an asterisk in (C)] are included for comparison. S*cale bars* in (A) and (B): 200 μm, and in 250 μm. Arrow in (C) marks worm that is magnified (4×) in panel (D). (**D**) Higher resolution imaging reveals activation of P*tbb-6*∷GFP in the gut of worms expressing PMK-3(L) under the *rgef-1* promoter is due to reporter mis-expression and not cell non-autonomous signaling from neurons to gut cells (*arrow*). Additional worm images are provided in **Fig. S6**. (**E, F**) Higher magnification images of transgene-containing worms in (A) and (B) cultured on1/10^th^ strength *atp-3* or *isp-1* RNAi. For each listed strain, only relevant genotype information is provided (see **Table S1** for full details). In all images, mCherry expression indicates presence of the relevant extrachromosomal array and production of transgenic PMK-3.

Surprisingly, GFP was expressed solely in the gut in each case, despite bright neuronal mCherry expression with both the ubiquitous *eft-3* promoter (**Fig. 2A**) and the neuron-specific *rgef-1* promoter (**Fig. 2C**). Initially the result with the *rgef-1* promoter seemed to imply cell non-autonomous signaling traveling from the neurons to the gut. However, closer inspection of this line revealed that the *rgef-1* promoter was leaking expression in gut tissue (**Fig. 2D**, **Fig. S6A**). When we re-engineered our construct to contain a small decoy open reading frame (ORF) 5’ of the PMK-3 start site in an effort to reduce potential bleed through transcription (**S1 Text, Fig. S6B**), we still observed mis-expression of mCherry in gut cells. Finally, use of the *rab-3* promoter, another widely used neuron-specific promoter, and also re-engineered to contain a small decoy ORF to combat potential bleed through transcription, again resulted in translation of mCherry in gut tissue. We therefore conclude that these models of neuron-specific gene expression are unreliable. Nonetheless, they clearly show that despite expression of PMK-3 in the neurons, retrograde signaling is not activated in this tissue.

Since we have previously reported a constitutively active form of DLK-1 is fully capable of functioning in neurons to cell-autonomously activate P*tbb-6*∷GFP expression (17), our findings suggest neurons are recalcitrant to feeding RNAi, consistent with other studies. In summary, we conclude that PMK-3(L) restricted to the gut is sufficient to mediate mitochondrial stress-induced retrograde response signaling. We also find no evidence that neurons are required to signal cell non-autonomously to activate the PMK-3 retrograde response in gut cells; rather, all evidence points to intestinal cells autonomously activating this pathway. Further accentuating the role of intestinal PMK-3 in responding to ETC stress, worms that express PMK-3 either ubiquitously or just in the gut, indistinguishably exhibit P*tbb-6*∷GFP reporter gene induction only in gut cells when exposed to *atp-3* feeding RNAi targeting mitochondrial complex V (compare **Fig. 2E** and **2F**).

### Life extension following isp-1 feeding RNAi is fully restored by gut-specific PMK-3(L) expression

Since we observed PMK-3 retrograde response signaling was restricted to the gut following mild mitochondrial ETC dysfunction induced by *isp-1* feeding RNAi, we next investigated whether activation of this pathway in this tissue was sufficient to extend life. We measured the survival of worms with and without various PMK-3 arrays following exposure to two different doses of *isp-1* RNAi (1/10 and 1/2 strength). Both concentrations of *isp-1* RNAi are sufficient to lengthen the life of wild-type worms in a dose-dependent manner, a response that is severely blunted in *pmk-3(tm745)* mutants, as previously reported (**Fig. 3A**).

**Figure 3.**
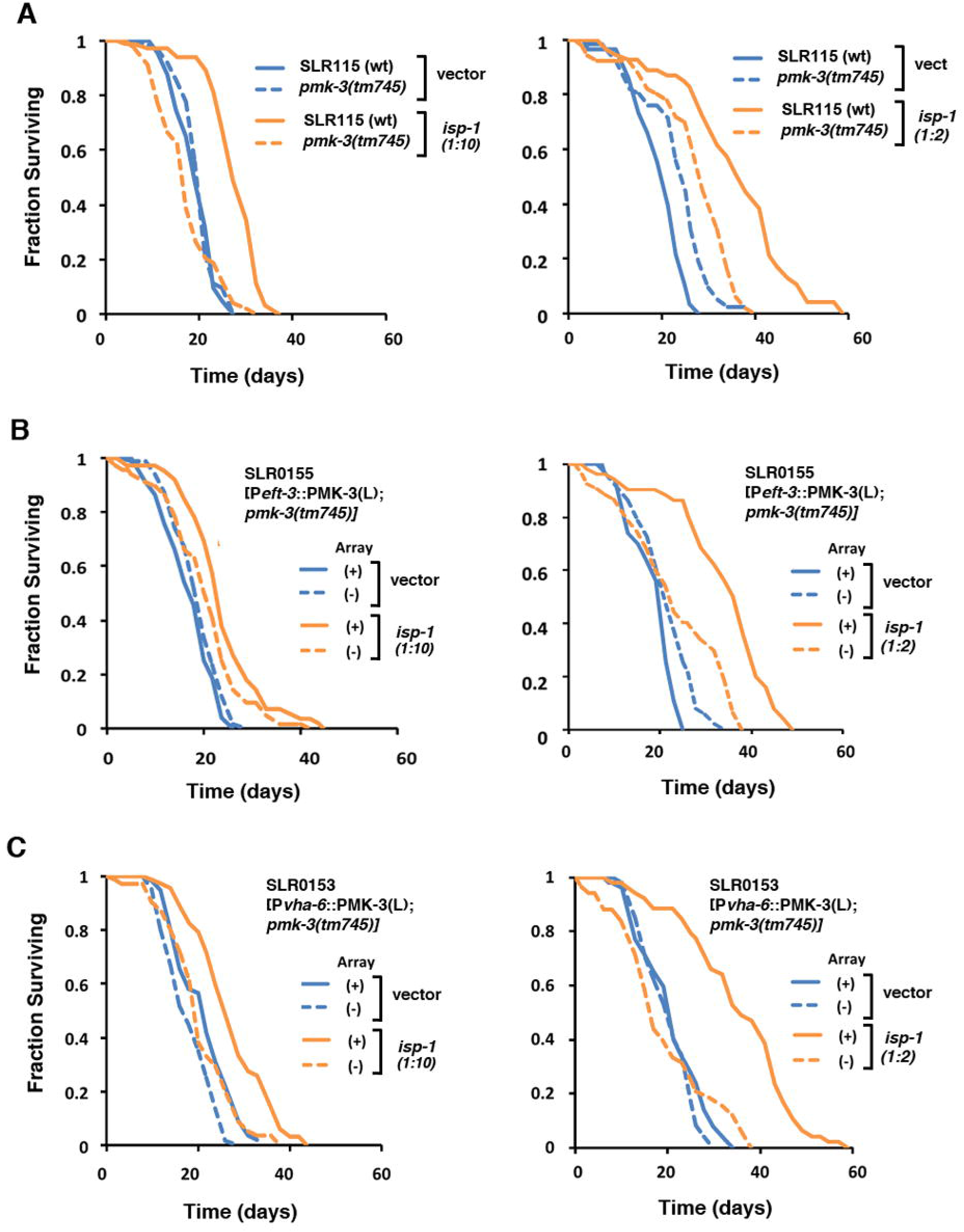
Life extension induced by*isp-1* feeding RNAi is fully restored by intestinal PMK-3 re-expression in*pmk-3(tm745)* mutants.Two different doses of *isp-1* bacterial feeding RNAi were tested. Two different doses of isp-1 bacterial feeding RNAi were tested - 1/10^th^ strength (*left panels*) and ½ strength (*right panels*). Survival statistics for all panels is provided in Table S2. (**A**) Wild type worms (strain SLR115) exhibit a dose dependent increase in life span when fed RNAi targeting *isp-1.* This response is abolished in *pmk-3(tm745)* knock-out mutants. (**B**) Ubiquitous re-expression of PMK-3(L) in *pmk-3(tm745)* null mutants restores the longevity response invoked by *isp-1* feeding RNAi (strain SLR0155). (**C**) Full life extension is also restored to *pmk-3(tm745)* mutants when PMK-3(L) re-expression is restricted to the gut (strain SLR0153). In B and C worms that have lost the array (−) serve as controls for potential strain background effects. Full genotype information for each strain is provided in **Table S1**.

Strikingly, PMK-3(L) expressed just in the gut permitted both doses of *isp-1* RNAi to extend the life of *pmk-3(tm745)* mutants to the same extent as ubiquitously-expressed PMK-3(L) (**Fig. 3B, C**). This result was replicated using an independently-generated P*vha-6∷PMK-3(L)-*containing strain (**Fig. S7A**). This limited need for only gut-specific PMK-3(L) became further apparent when the same set of strains were exposed to bacterial feeding RNAi targeting *atp-3*. In wild-type worms, severe knockdown of *atp-3* results in pathological shortening of life, in contrast to mild *atp-3* knockdown which extends life (13). In *pmk-3(tm745)* mutants, exposure to full strength *atp-3* RNAi results in L3 larval arrest and premature death (**Fig. S7B**). Re-expression of PMK-3(L) in the gut of these animals, however, leads to larval arrest by-pass in half of the animals, doubling population survival time relative to non-array containing worms. The degree of recovery mediated by gut-specific PMK-3(L) is again indistinguishable from the rescue conferred by ubiquitously re-expressed PMK-3(L) under the *eft-3* promoter (compare **Fig. S7C** and **Fig. S7D**). These results further underscore the notion that the gut appears to be the primary tissue mediating longevity signaling in worms exposed to ETC disruption.

### Constitutive activation of DLK-1 in the intestine is sufficient to drive activation of stress signaling

As mentioned previously, we have shown neuronally-expressed, constitutively active DLK-1 drives *Ptbb-*6∷GFP expression in the neurons (17). As our studies now suggest the intestine is the primary site of longevity signaling, we asked whether constitutive DLK-1 activation in the gut could drive retrograde signaling in the absence of ETC disruption. To do this, we coupled DLK-1b(EE), the constitutively active form of DLK-1, to the gut-specific *vha-6* promoter along with an mCherry transcriptional reporter and injected it into *Ptbb-6*∷GFP worms. Constitutively active, gut-specific DLK-1b(EE) successfully induced *Ptbb-6*∷GFP reporter expression in the absence of overt ETC disruption (**Fig. 4A, B**). GFP fluorescence was limited to the gut, consistent with cell autonomous activation by DLK-1b(EE). DLK-1b(EE) overexpression did not alter endogenous PMK-3 levels beyond the background signal detectable in worms that had lost the array (**Fig. S4A, B**).

**Figure 4.**
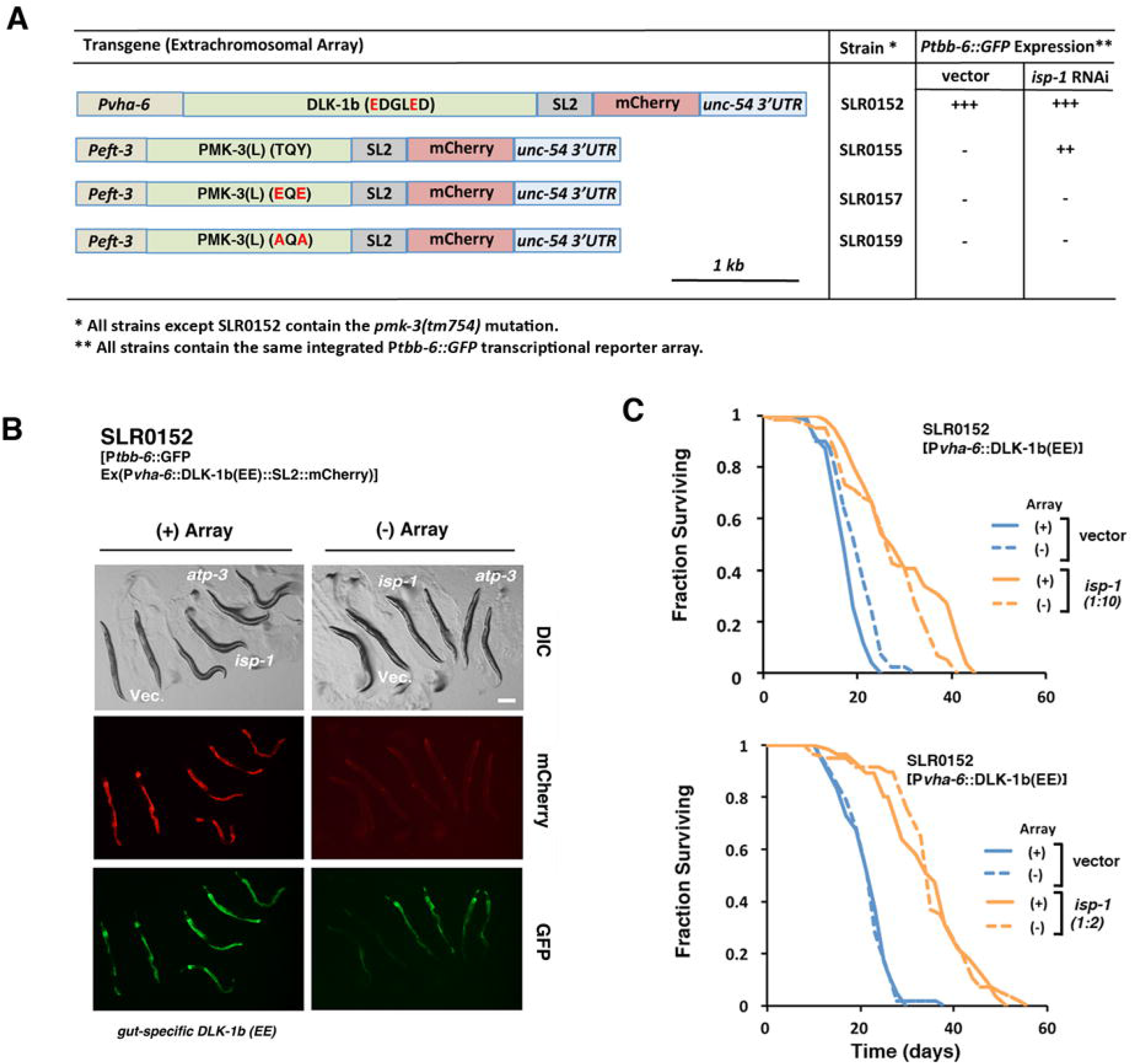
Constitutive activation of DLK-1 in the intestine is sufficient to drive activation of stress signaling. (**A**)Schematic showing *dlk-1* and mutant *pmk-3* transgene generation. Constructs are comprised of a two-gene operon assembled as follows: (*left* to *right*) promoter (P*vha-6* or P*eft-3)*, *dlk-1* or *pmk-3* cDNA, splice leader sequence (SL2), mCherry reporter, *unc-54* 3’UTR. Strain names of worms carrying the relevant transgene are indicated, along with a summary of the effect of each transgene on *Ptbb-6∷GFP* reporter expression under the listed conditions (see main text for details). (**B**) Wild type worms expressing DLK-1b(EE) exhibit constitutive activation of the PMK-3 dependent reporter gene, P*tbb-6*∷GFP, independent of mitochondrial ETC disruption (compare animals cultured on bacteria containing vector control (vec.) and *atp-3* or *isp-1* feeding RNAi. Worms that have lost their array (−) are included for comparison. Scale bar: 200 μm. (**C**) Expression of DLK-1b(EE), a constitutively active form of DLK-1, in the gut of wild type worms is by itself insufficient to extend life. Moreover, exposure of these animals to *isp-1* feeding RNAi (1/10^th^ strength (*upper panel)* and ½ strength (*lower panel),* does not extend life beyond that observed for wild type animals exposed to the same conditions (compare to Figs. 3A*left* and *right panels*, respectively).

We next sought to determine whether constitutive activation of PMK-3 is also sufficient to drive life extension in the absence of mitochondrial stress. p38 MAPKs are activated by MAP2K phosphorylation of a TGY motif (30). PMK-3 differs from most other p38 MAPKs in that its activation loop contains a predicted TQY motif instead of the canonical TGY motif. We generated two different double point mutations in PMK-3: AQA and EQE. These mutations are predicted to result in permanent inactivation and activation, respectively. These mutated proteins were expressed under the *eft-3* promoter. Interestingly, presence of the EQE phosphomimetic mutation in PMK-3(L) induced a conformational alteration in the structure of the protein that caused the recombinant protein to migrate slower by SDS-PAGE (**Fig. S4A, B**), much as phosphorylation often does for other MAPKs. Unlike constitutively active DLK-1, PMK-3(L) containing the EQE phosphomimetic mutation failed to activate *Ptbb-6*∷GFP reporter induction under basal conditions (**Fig. 4A**). Surprisingly PMK-3(L) EQE also failed to mediate P*tbb-6*∷GFP induction in worms exposed to *isp-1* RNAi (**Fig. 4A**). These findings suggest that the EQE mutation did not function as planned to induce clearance of the activation loop from the catalytic-binding site. Instead, the sequence change appears to function as a dominant negative and blocks natural activation of PMK-3 enzymatic activity.

### The lifespan of wild-type worms is not extended when DLK-1 is constitutively activated in the intestine in the absence of ETC stress

We next evaluated whether the activation of retrograde signaling in worms expressing constitutively active, gut-specific DLK-1 in the absence of ETC stress corresponded to an increase in lifespan. Unlike DLK-1b(EE) expressed in neurons where worms became sterile and died (17), we observed no detrimental effect of overexpressing DLK-1b(EE) in the gut. However, DLK-1b(EE) was not sufficient to drive life extension (**Fig. 4C**). Moreover, the presence of activated DLK-1b(EE) in the gut did not prevent or further enhance the extension of life that is induced when animals are cultured on *isp-1* RNAi (**Fig. 4C**). We conclude that while DLK-1/PMK-3 signaling in intestinal cells is necessary to extend life following mild mitochondrial ETC disruption, constitutive activation of this pathway by DLK-1b(EE) overexpression in the gut of wild type worms is alone insufficient.

### Age-dependent dampening of the DLK-1/PMK-3 signaling axis in wild type worms

One explanation for why DLK-1b(EE) overexpression is unable to extend the life of wild-type worms is because, in the absence of mitochondrial dysfunction, this signaling pathway might become dampened over time. In support of this possibility, we observed that activation of *Ptbb-*6∷GFP in DLK-1b(EE) overexpressing worms was robust in 3 day-old adults but markedly attenuated in 13 day-old adults, despite strong expression of the mCherry reporter gene at both time points (**Fig. 5A-C**). Strikingly, this age-dependent decrease in *Ptbb-*6∷GFP fluorescence did not occur in animals exposed to *isp-1* RNAi (**Fig. 5B, C**). To test the possibility that endogenous PMK-3 becomes transcriptionally repressed with age, we co-expressed DLK-1b(EE) and PMK-3(L) under the constitutive *vha-6* promoter. Again, despite strong mCherry signal indicating transcription for these genes is not dampened, *Ptbb-6∷GFP* fluorescence was greatly diminished with age (**Fig. 5D**) and lifespan was not extended (**Fig. 5E**). We conclude that loss of DLK-1/PMK-3 signaling with age is not due to decreased transcription of these genes, but a yet unknown mechanism which is overcome by mild mitochondrial ETC disruption.

**Figure 5.**
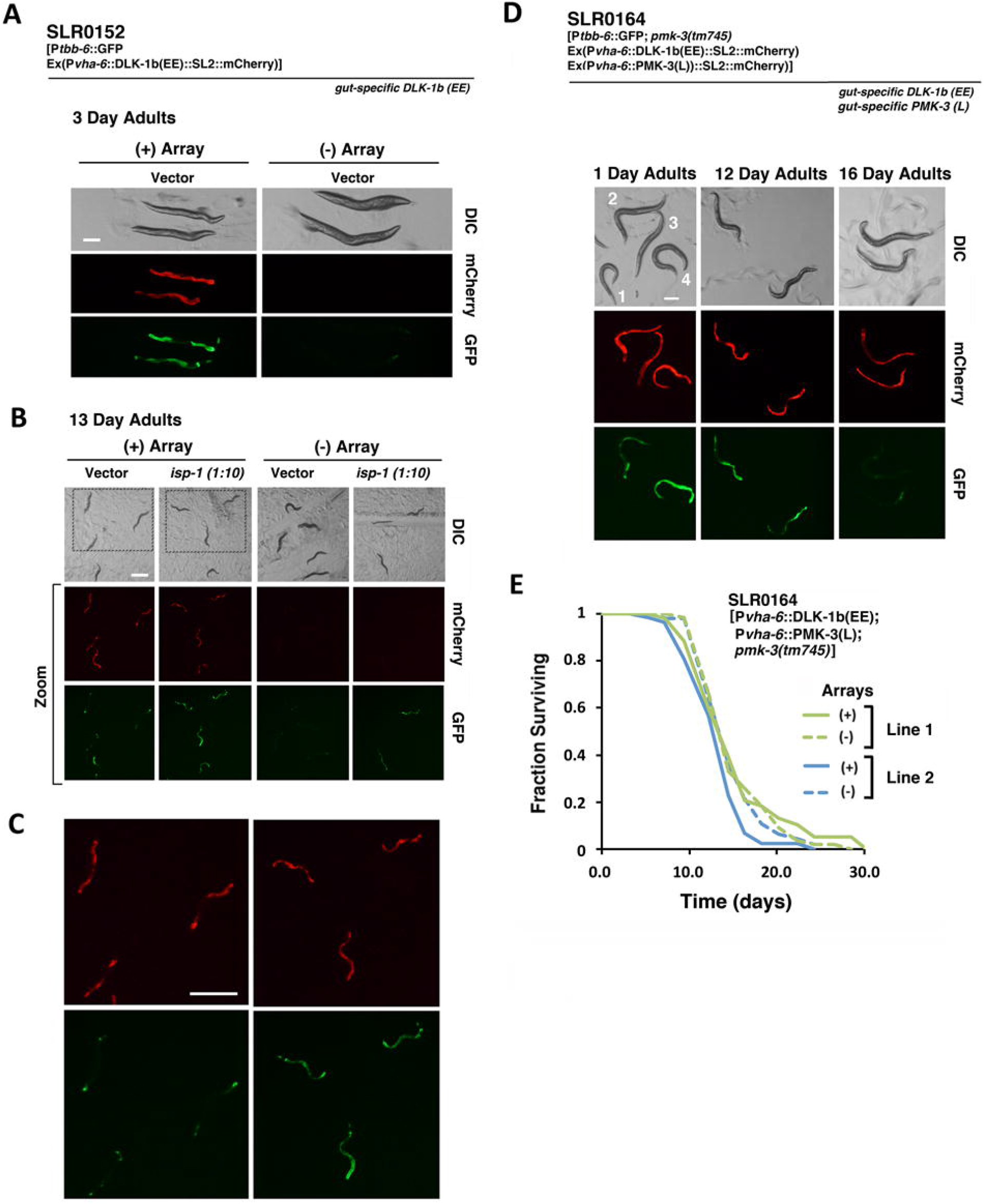
Age-dependent inactivation of the DLK-1/PMK-3 signaling axis. (**A-C**) By two weeks of age, constitutive P*tbb-6*∷GFP reporter expression is lost in wild type worms containing the DLK-1b(EE) transgenic array (strain SLR0152), even though the promoter driving DLK-1 is still transcribed [compare mCherry signal to GFP signal in (A) and (B)]. By contrast, in SLR0152 worms that have been exposed to *isp-1* feeding RNAi (1/10^th^ strength) throughout their life, P*tbb-6*∷GFP reporter expression remains robustly activated at two weeks of age. An enlargement of the fluorescent images in (B) is shown in panel C [specific areas are marked in the left two DIC images of panel B (*dotted boxes*)]. Worms that have lost their array (−) in (A) and (B) are included for comparison. All scale bars: 500 μm. (**D, E**) SLR0164 worms contain two independent arrays expressing DLK-1b(EE) and PMK-3(L), respectively. Both transgenes are controlled by the gut-specific *vha-6* promoter. In (D), *upper left panel* illustrates SLR0164 worms that have lost both arrays (#1), worms with both arrays (#2, 4), and worms with only the DLK-1b(EE) array (#3). Even though PMK-3(L) is constitutively expressed by the *vha-6* promoter, and activated by DLK-1b(EE) early in life in worms containing both arrays, as marked by P*tbb-6*∷GFP expression, the reporter gene is gradually dampened with age until it is off completely by day 16. Accordingly, SLR0164 worms exhibit no life extension (E). Survival data for two independent lines is shown. Siblings lacking both extrachromosomal arrays are included as controls. Survival statistics are provided in **Table S2.** Scale bar in (D): 200 μm.

### Post-translational regulation of PMK-3(L) following mitochondrial ETC disruption

Based on the functioning of other MAP3K-MAP2K-MAPK signaling cascades, we hypothesized that PMK-3 is activated by phosphorylation of its TQY motif by SEK-3 following ETC stress. In support of these predictions, we observe that gut-specific re-expression of a *myc*-tagged DLK-1b(EE) in *dlk-1(tm4024)* knockout mutants is sufficient to permit P*tbb-6∷GFP* induction following ETC disruption (**Fig. S8A, B**). This finding places both the start (DLK-1) and end (PMK-3) of this MAPK signaling cascade in the same tissue. The observation that a double point mutation converting the TQY motif to either AQA or EQE blocked the ability of PMK-3(L) to induce *Ptbb-6∷GFP* reporter gene expression in worms exposed to *atp-3* or *isp-1* knockdown also underscores the importance of these two residues (**Fig. 4A**).

While we were unable to generate a phospho-PMK-3-specific antibody to directly detect PMK-3 phosphorylation following ETC stress, considerable evidence points to phosphorylation as the mechanism of PMK-3 activation. MAPKs are inactivated by a family of enzymes called dual-specificity phosphatases (DSPs), each of which contains a specific MAPK-interaction motif and a C-terminal phosphatase domain that acts on both the threonine and tyrosine residues contained within the activation loop of its target MAPK (30). *vhp-1* is orthologous to the human DSP8 subfamily of DSPs which are specific for JNK and p38 MAPKs. We have previously reported that *isp-1(qm150)* mutants cultured on *vhp-1* RNAi arrest as L3 larvae and that this arrest can be overcome by removal of PMK-3 (17). This finding is consistent with PMK-3 being activated by phosphorylation on its activation loop while VHP-1 inactivates it through dephosphorylation. Furthermore, phosphorylation of the tyrosine residue of the TQY motif of PMK-3 has been detected in a prior high-throughput analysis of the *C. elegans* phosphoproteome where worms were cultured to high density in liquid media (31). This kind of growth environment is known to be limiting for oxygen tension, mimicking the effects of ETC disruption (32).

In addition to these observations, we now also report that when *pmk-3(tm745)* mutants containing the P*vha-6*∷PMK-3(L) transgene are cultured on bacterial feeding RNAi targeting *atp-3* or *isp-1,* and then whole-worm extracts assessed for changes in PMK-3(L) abundance or molecular weight, we observe no differences relative to vector-treated worms (**Fig. S4C, D**). These findings argue against ETC disruption causing acute changes in either PMK-3 translation, covalent multimerization, or protein cleavage that might otherwise result in activation. Finally, worms containing an extrachromosomal array encoding GFP fused in-frame to the N-terminal portion of PMK-3 and driven by its non-operonic promoter exhibit no change in GFP fluorescence intensity or localization following *isp-1* knockdown (**Fig. S9**). This finding argues against PMK-3(L) being activated by means of either increased transcription or regulated re-distribution of the protein. Substantial evidence therefore points toward phospho-regulation of one or both residues within the TQY motif of PMK-3 as the mode by which this kinase is activated by SEK-3 following ETC stress.

### Computational identification of PMK-3-regulated targets

In order to gain insight into the PMK-3-dependent processes that mediate life extension following ETC disruption, we used a computational approach to identify candidate downstream targets of PMK-3 signaling. We have previously reported that the promoter of *tbb-6* contains two identical copies of a motif that is most similar in sequence to the DNA binding motif of human CCAAT/enhancer binding protein β (C/EBPβ) (17). We showed that both copies of this motif are essential for transcriptional induction of *tbb-6* following PMK-3 activation by ETC stress. These and other studies led us to define a consensus C/EBPβ-like motif (**Fig. 6A**, *top*), hereafter referred to as “Motif 2” in accord with (17). For the present study, we make the working assumption that PMK-3 controls binding of an unidentified transcription factor that recognizes Motif 2. We hypothesize, then, that Motif 2 promoter elements which remain strictly conserved across evolutionary time and which are localized to genes encoding components of the same biochemical pathway, define mechanisms that are required for PMK-3-mediated survival in response to ETC stress.

**Figure 6.**
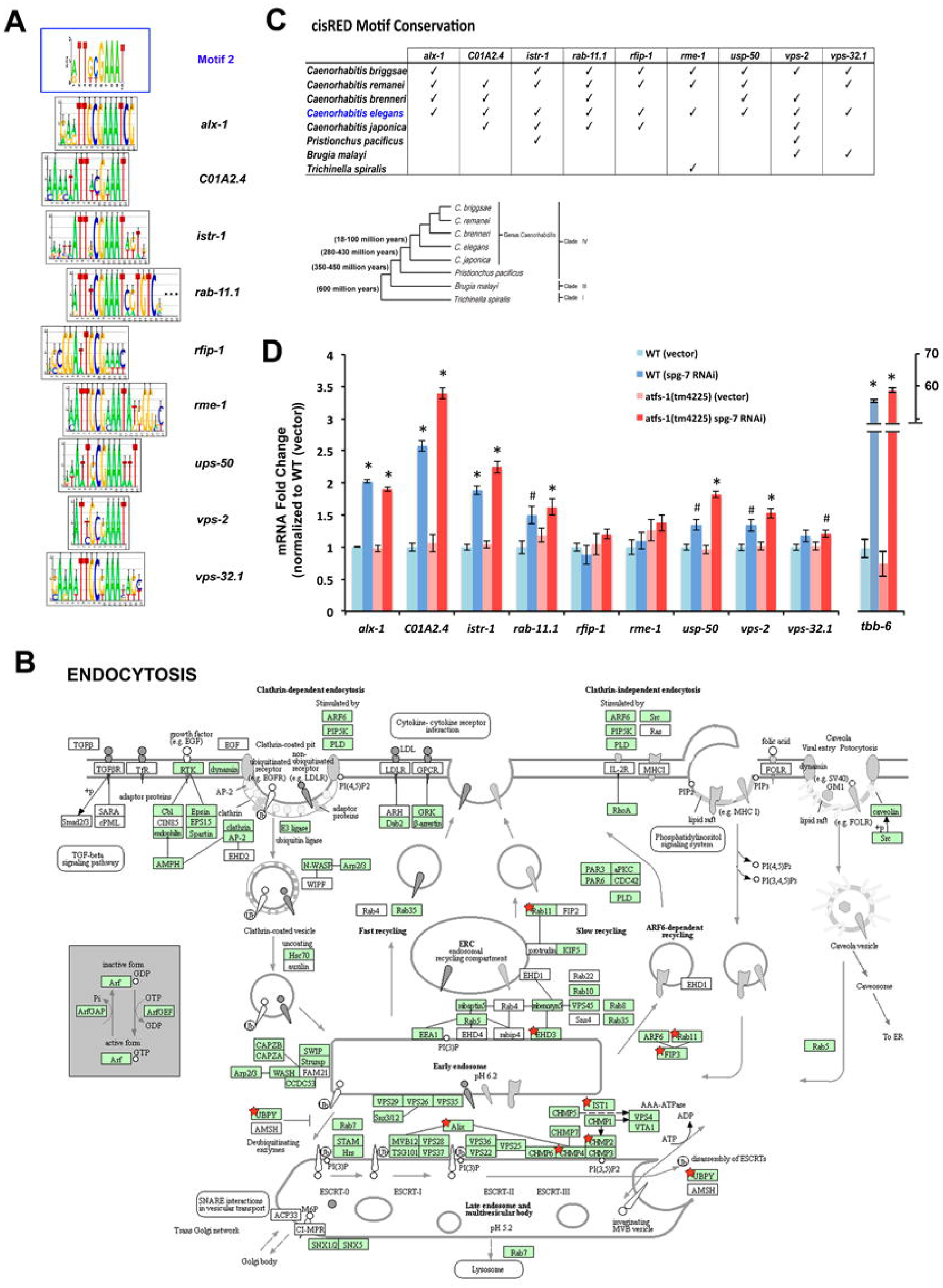
A Computational Approach for Identifying PMK-3 target proteins. (**A**) Consensus sequences in the promoters of nematode endocytic genes matching “Motif 2”. Sequence LOGOs were generated using cisRED (22). (**B**) Generic KEGG pathway map of endocytosis (24). Human proteins are shown. Proteins with unambiguous *C. elegans* orthologs are colored *green.* Worm genes containing Motif 2 are highlighted with a *red star*. (Box and star for CHMP2 includes both CHMP2A and CHM2B). (**C**) *Upper panel –* Summary of nematode orthologs used for construction of each LOGO in (A). *Lower panel* - Phylogenetic tree of nematode species used by cisRED. Genetic distances are not drawn to scale. Divergence times based on (57). (**D**) Affymetrix mRNA expression analysis of wild type (WT) worms and *atfs-1(tm4525)* mutants exposed to *spg-7* feeding RNAi (58). Bars illustrate mean (+/− SD) of three independent biological replicates for each tested condition. All data was normalized to wild type worms on vector control for each listed gene. (Student’s t-test, * *q-value* <0.05, # *q-value* <0.07).

The cisRED database contains a collection of 158,017 novel conserved promoter motifs that reside upstream of 3,847 conserved *C. elegans* transcripts, spanning some 600 million years of nematode evolution (22). We screened this library for matches to Motif 2 and identified 206 conserved promoter sites spread over 188 genes. Among the protein products encoded by these genes, we discovered that components of the endocytic pathway were significantly over-represented (Benjamini-Hochberg *q-value* 0.0025). Nine genes were associated with this biochemical pathway (**Fig. 6B**), all of which have human orthologs (shown in brackets): *alx-1* (ALIX), C01A2.4 (CHMP2B), *istr-1* (IST1), *rab-11.1* (Rab11), *rfip-1* (FIP3), *rme-1* (EHD3), *ups-50* (UBPY), *vps-2* (CHMP2A) and *vps-32.1* (CHMP4C). Intriguingly, five of the nine genes encode subunits of the endosomal sorting complexes required for transport-III (ESCRT-III) complex, which plays a pivotal role in the biogenesis of multivesicular bodies (MVBs) and, in association with RAB-11, exosome and ectosome formation (33). Both of these latter structures are used for communication between cells (34). The consensus promoter sequence for each of the nine genes that match Motif 2 is shown in **Fig. 6A** *(upper panel)*. Orthologous genes present in the cisRED database that contributed to each consensus sequence are shown in **Fig. 6C** *(upper panel)*, along with the evolutionary relationship between the nematode species from which these orthologs are derived *(lower panel)*. We conclude that a signaling pathway utilizing Motif 2 has been conserved for at least 100 million years of nematode evolution to control endosomal trafficking (**Fig. 6B**).

### Experimental validation of PMK-3-regulated targets

To determine the biological relevance of our findings, we tested if the expression of each of the nine endocytic pathway genes identified above was enhanced by mitochondrial ETC disruption. We also tested if, like *tbb-6*, these newly identified genes exhibited transcriptional independence from ATFS-1. To do this we utilized a published microarray dataset (**S1 Text**), with many of the genes we tested being represented by multiple primer sequences. We confirmed that seven of the nine genes exhibited significant upregulation following mitochondrial disruption induced by *spg-7* feeding RNAi, in a manner that was independent of *atfs-1* (**Fig. 6D**).

We next focused on four of the most strongly responsive genes: *alx-1*, C01A2.4 (CHMP2B), *istr-1* and *rab-11.1*. We tested whether their removal in wild-type worms blocked the life extension that is normally observed when these animals are placed on *isp-1* feeding RNAi. Remarkably, we observed that knockdown of *istr-1* completely abolished life extension under these circumstances (**Fig. 7A**), an effect that was not due to reduced RNAi efficacy (**Fig. S10** and **S1 Text**). Likewise, knockdown of *rab-11.1* also completely abolished life extension following ETC disruption by *isp-1* feeding RNAi. However, *rab-11.1* knockdown caused worms to die much earlier in the absence of ETC stress (**Fig. 7A**). Knockdown of C01A2.4 induced a small but significant reduction in life extension following exposure to ETC disruption, and had no effect in the absence of ETC stress (**Fig. 7A**). Finally, we observed no effect of *alx-1* knockdown on *isp-1* RNAi-induced life extension, though lifespan was shortened under basal conditions (**Fig. 7A**).

**Figure 7.**
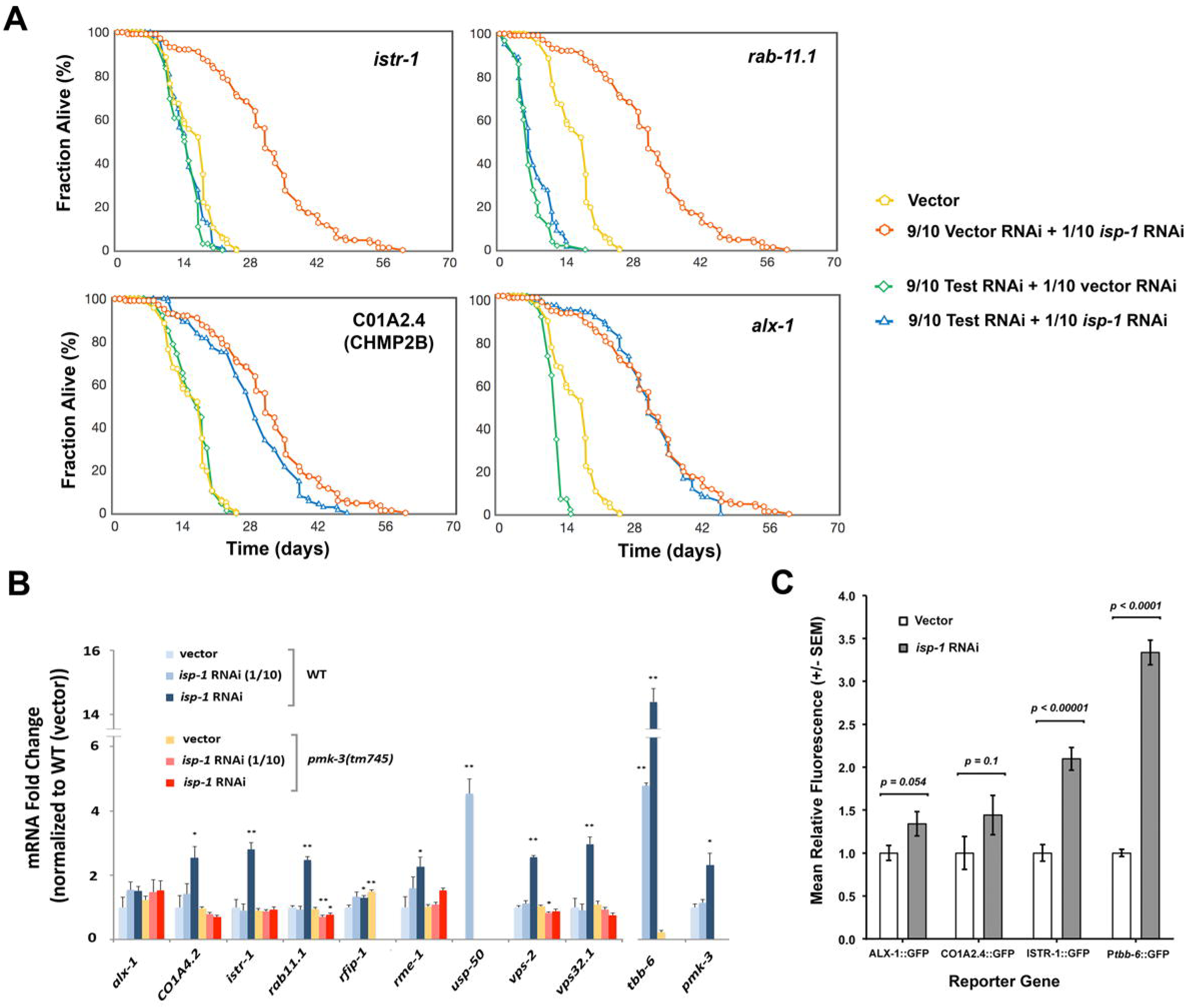
ESCRT Proteins Play an Essential Role in the PMK-3 Retrograde Response. (**A**) Survival analyses of wild type worms (SLR115) exposed to dual feeding RNAi targeting *isp-1* (1/10^th^ strength) and the listed ESCRT gene (9/10^th^ strength). Data was pooled from two independent experiments (N = 95 - 180 worms per tested condition). (*istr-1*+*isp-1 vs.* vector) and (*rab-11.1*+*isp-1 vs.* vector), Bonferroni corrected *p-*values 0.0; (C01A2.4+*isp-1 vs. isp-1*), Bonferroni corrected *p*-value <0.04. (**B**) qRT-PCR analysis of ESCRT genes in wild type worms (SLR115) and *pmk-3(tm745)* knockout mutants (SLR0150) following growth on 1/10^th^ strength- or undiluted *isp-1* feeding RNAi from the time of hatching until the first day of adulthood. *pmk-3* and *tbb-6* genes were included as controls. Bars illustrate mean (+/− SEM) from seven independent biological replicates. Data was normalized to the wild type vector control for each listed gene. (Student’s one-tailed t-test, * *p-value* <0.05, ** *p-value* <0.01). (**C**) ‘Third allele’ versions of ALX-1, CO1A2.4 and ISTR-1 encoding C-terminal GFP fusion proteins were generated in HT1593 [*unc-119(ed3)*] worms by microinjection. Extrachromosomal arrays also contained a rescuing UNC-119 marker gene. Each strain was cultured on *isp-1* feeding RNAi and GFP fluorescence recorded at the first day of adulthood. Data is presented relative to control (vector-only) treated animals. (N = 18-24 worms per condition; mean +/− SEM).

To formally test whether PMK-3 mediates the downstream activation of the nine endocytic pathway genes identified from the cisRED database in response to mitochondrial ETC disruption, we measured mRNA expression of these genes in worms either containing or lacking a functional *pmk-3* locus and following growth on *isp-1* feeding RNAi. Strong *isp-1* knockdown resulted in a significant increase in the abundance of six endocytic genes (*C01A2.4* (CHMP2B), *istr-1*, *rab-11.1*, *ups-50*, *vps-2* and *vps-32.1*) in a *pmk-3* dependent manner (**Fig. 7B**). To assess whether these changes in mRNA populations were functionally relevant at the protein level, we generated translational reporter genes corresponding to *alx-1*, C01A2.4(CHMP2B), and *istr-1* fused to GFP and then expressed them as “third alleles” (**S1 Text**). We observed only a significant increase in ISTR-1∷GFP fluorescence when worms were exposed worms to *isp-1* feeding RNAi (**Fig. 7C**). Combined with the lifespan data, this points to ISTR-1 as a key downstream mediator of PMK-3-dependent retrograde response signaling which functions to promote longevity in worms.

## DISCUSSION

In recent years it has become increasingly clear that multiple retrograde responses function to extend life in *C. elegans* following mild mitochondrial ETC disruption (12). The specific mode of ETC disruption (genetic mutation, RNAi knockdown or chemical inhibition), as well as the mitochondrial complex targeted by that disruption, together determine the nature of the cell response and the subsequent life-extending mechanisms that are activated. Several recent studies have shed light on some of these life-extending processes such as activation of HIF-1 following the aberrant accumulation of mitochondrial-derived α-ketoacids (16), changes in global gene expression following epigenetic reprogramming of histones (35, 36), autophagic clearance of damaged mitochondria (37), activation of the intrinsic apoptotic pathway (38), stimulation of DNA damage response proteins (39), and activation of the UBL-5 sub-branch of the mitochondrial unfolded protein response (36). Several proteins with less clear functions, including *ceh-23* and *taf-4* have also been implicated in longevity signaling (9, 40). The PMK-3-dependent pathway examined in this study acts independently of all well-known life-extending pathways in *C. elegans* (17) and, to our knowledge, represents a novel survival mechanism.

In the current work, we expanded our understanding of the molecular and physiological details behind the PMK-3 retrograde response. We showed that the full-length isoform of PMK-3 is specifically required for life extension following ETC disruption, and that restriction of this isoform to the worm intestine is sufficient to confer complete life extension under these conditions. Our data also show that neurons do not activate the PMK-3 retrograde response cell autonomously following feeding RNAi targeting *atp-3* or *isp-1*, even though these cells are fully capable of activating this pathway (17). Our data also reveal a technical flaw in use of the *rgef-1* and *rab-3* neuronal promoters when expressed from an extrachromosomal array. While both reporters clearly induce strong neuronal expression, they nonetheless also result in mis-expression in the gut. The strong quantum yield of mCherry, as opposed to that of GFP, made it possible for us to capture this mis-expression, and also to avoid making the wrongful conclusion that intestinal retrograde response signaling is activated cell non-autonomously via neuronal PMK-3 signaling. Our findings underscore the worm intestine as the most important site for PMK-3 to activate life-promoting signals in response to ETC disruption.

Previous studies using *C. elegans* have implicated both intestinal and neuronal tissue in longevity signaling following mitochondrial ETC disruption (41, 42). Notwithstanding the potential confound of neuronal promoters leaking on in gut tissue, a model was developed whereby mitochondrial stress in neurons causes the release of a “mitokine” that relays a signal to the intestine to upregulate the mitochondrial unfolded protein response (UPR^mt^) and promote longevity. Although a role for the UPR^mt^ in longevity specification is contentious (41, 43), support for the dual tissue component of this model was provided by two studies from the same group. Mild mitochondrial ETC disruption was shown to induce distinct epigenetic alteration of histones in neuronal and intestinal cells, both of which were required for life extension. The neuronal pathway utilized the histone demethylases JMJD-1.2 and JMJD-3.1 (36), while the intestinal pathway required DVE-1, LIN-65, MET-2 and UBL-5 (35). It is possible that the PMK-3 retrograde response examined in the current study intersects with the DVE-1/LIN-65/MET-2/UBL-5 retrograde response to control longevity, since both pathways localize to the intestine. One possibility is they act linearly. However, we find that knockdown of DVE-1 does not prevent P*tbb-6*∷GFP expression, which is under the control of PMK-3. Moreover, we have previously reported that ATFS-1, which functions downstream of DVE-1/LIN-65/MET-2/UBL-5 to turn on the UPR^mt^, does not require PMK-3 (17). Another possibility is that both retrograde responses converge to control longevity. This might explain why overexpression of a constitutively active form of DLK-1 in the gut was unable to extend life in the absence of ETC disruption – either other factors controlled by the DVE-1/LIN-65/MET-2/UBL-5 retrograde response needed to be present or the necessary epigenetic reprograming of gut cells by this system was not in place. Finally, it is also possible that the PMK-3 and DVE-1/LIN-65/MET-2/UBL-5 retrograde responses function independently of each other.

One surprising finding that arose from our current study is the apparent decline in DLK-1/PMK-3 signaling capacity with age. Loss of DLK-1 signaling capacity seems to be the simplest reason why life was not extended when constitutively active DLK-1 was overexpressed in wild type worms in the absence of ETC dysfunction. We have excluded transcriptional downregulation as the mechanism by which signaling is inactivated, but many potential factors remain to be tested such as increased DLK-1 turnover. Likewise, what factors or signals from dysfunctional mitochondria establish conditions that are permissive to the continued expression of the DLK-1/PMK-3 pathway as animals age is also unclear. As mentioned previously, epigenetic reprograming by the DVE-1/LIN-65/MET-2/UBL-5 retrograde response is one possibility. Other possibilities relate to mitokine signaling, or downregulation of *vhp-1*. Alternatively, Jeong and colleagues (44) recently identified a novel mechanism by which the p38 MAPK PMK-1 is activated in *C. elegans*. Studying the innate immune response of worms exposed to pathogenic *Pseudomonas aeruginosa* (PA14), they found that a cytosolically-localized form of the mitochondrial chaperone HSP-60 physically bound to and stabilized the MAP2K SEK-1. This, in turn, up-regulated PMK-1 activity and increased PA14 resistance. Whether a related mechanism acts to stabilize DLK-1 or SEK-3 following mild mitochondrial ETC disruption remains to be determined.

We have additionally shown there may be a role for the endosomal trafficking pathway in mediating ETC stress-induced life extension downstream of PMK-3. In particular, we identified the ESCRT-III-associated proteins ISTR-1 and C01A2.4 (CHMP2B), as well as RAB-11.1 as necessary for this longevity (45–47). The ESCRT-III machinery plays important roles in extracellular vesicle (EV) formation. EVs can be categorized into two broad types depending on their mode of formation. Exosomes originate late in the endocytic pathway and are formed when multivesicular bodies (MVBs) are re-routed for fusion with the plasma membrane instead of the lysosome. Ectosomes bud directly from the plasma membrane. In both instances, ESCRT-III proteins play a pivotal role in vesicle formation and release, directing the final membrane scission event (48).

Recent studies have implicated extracellular vesicles in an array of functions, including the control of mouse aging by hypothalamic stem cells (49), cardiac rejuvenation (50), and cancer metastasis (51). Such findings raise the provocative possibility that *C. elegans* might use EVs to broadcast the metabolic status of its intestinal cells to the rest of the body to control longevity. On the other hand, these proteins might simply play a role in increased lysosomal turnover. Consistent with this possibility, it has been reported that mitophagy is increased in worms following disruption of various ETC targets (52, 53), and there is a well-established requirement for autophagy in promoting the longevity of mitochondrial ETC mutants (37, 54). Furthermore, genetic screens have linked DLK-1 and PMK-3 signaling with elevated endocytosis in neurons (28, 55), as well as increased RAB-11.1 levels and lysosomal trafficking in muscle cells (56). Nonetheless, it remains unclear whether increased endolysosomal trafficking or reduced extracellular vesicle formation was responsible for the phenotypes observed in each of these prior studies.

We predict that EVs emitted from the baso-lateral surface of intestinal cells into the coelomic cavity in response to PMK-3 activation will mediate signaling to the rest of the body to control longevity. However, formal testing of the role of EVs in PMK-3 retrograde response signaling is currently hampered by technical limitations which prevent isolation of EVs from specific tissues (26). Current, state-of-the-art isolation of EVs from *C. elegans* is restricted to extracorporeal populations from starved worms (26). When non-stressed worms were treated with these EV populations from worms with activated PMK-3 signaling to see if they might mediate a broader retrograde signal, we find not unexpectedly that they do not (**Fig. S11**). Biological and technical advances which permit EV signaling to be dissected in a tissue-specific manner are consequently an important area of need for evaluation of their role in the PMK-3 retrograde response and other pathways.

## Abbreviations

C/EBPβ: CCAAT/enhancer binding protein β
DLK-1b(EE): constitutively active form of DLK-1 isoform b
ESCRT-III: endosomal sorting complexes required for transport type III complex
ETC: electron transport chain
EVs: extracellular vesicles (exosomes and exosomes)
MAPK: mitogen-activated protein kinase
*Ptbb-*GFP∷6: GFP coupled to the promoter of *tbb-6*

## ACKNOWLEDGEMENTS

Several strains were provided by the CGC, which is funded by NIH Office of Research Infrastructure Programs (P40 OD010440). We thank Drs. Yishi Jin (UC San Diego) and Alex Soukas (Harvard University) for sharing plasmid reagents. Drs. Gert Jansen (Erasmus MC, Netherlands), Dennis Kim (MIT, USA) and Shohei Mitani (National Bioresource Project, Japan) for providing worm strains.

## REFERENCES

1. Lane RK, Hilsabeck T, Rea SL. The role of mitochondrial dysfunction in age-related diseases. Biochim Biophys Acta. 2015.

2. Melov S, Shoffner J, Kaufman A, Wallace D. Marked increase in the number and variety of mitochondrial DNA rearrangements in aging human skeletal muscle. Nucl Acids Res. 1995;23:4122–4126.

3. Nicholls DG, Ferguson SJ. Bioenergetics3. London: Academic Press; 2002.

4. Parekh AB, Putney JW, Jr. Store-operated calcium channels. Physiol Rev. 2005;85:757–810.

5. Rouault TA, Tong WH. Iron-sulfur cluster biogenesis and human disease. Trends Genet. 2008;24:398–407.

6. Khutornenko AA, Roudko VV, Chernyak BV, Vartapetian AB, Chumakov PM, Evstafieva AG. Pyrimidine biosynthesis links mitochondrial respiration to the p53 pathway. Proc Natl Acad Sci U S A. 2010;107:12828–12833.

7. Arnould T, Michel S, Renard P. Mitochondria Retrograde Signaling and the UPR(mt): Where Are We in Mammals? Int J Mol Sci. 2015;16:18224–18251.

8. Benedetti C, Haynes CM, Yang Y, Harding HP, Ron D. Ubiquitin-Like Protein 5 Positively Regulates Chaperone Gene Expression in the Mitochondrial Unfolded Protein Response. Genetics. 2006;174:229–239.

9. Khan MH, Ligon M, Hussey LR, Hufnal B, Farber R, 2nd, Munkacsy E, et al. TAF-4 is required for the life extension of isp-1, clk-1 and tpk-1 Mit mutants. Aging (Albany NY). 2013;5:741–758.

10. Owusu-Ansah E, Yavari A, Mandal S, Banerjee U. Distinct mitochondrial retrograde signals control the G1-S cell cycle checkpoint. Nat Genet. 2008;40:356–361.

11. Kirienko NV, Ausubel FM, Ruvkun G. Mitophagy confers resistance to siderophore-mediated killing by Pseudomonas aeruginosa. Proc Natl Acad Sci U S A. 2015;112:1821–1826.

12. Munkácsy E, Rea SL. The paradox of mitochondrial dysfunction and extended longevity. Exp Gerontol. 2014;56:221–233.

13. Rea SL, Ventura N, Johnson TE. Relationship Between Mitochondrial Electron Transport Chain Dysfunction, Development, and Life Extension in Caenorhabditis elegans. PLoS Biol. 2007;5:e259.

14. Yang W, Hekimi S. Two modes of mitochondrial dysfunction lead independently to lifespan extension in Caenorhabditis elegans. Aging Cell. 2010;9:433–447.

15. Butler JA, Mishur RJ, Bhaskaran S, Rea SL. A metabolic signature for long life in the Caenorhabditis elegans Mit mutants. Aging Cell. 2013;12:130–138.

16. Mishur RJ, Khan M, Munkacsy E, Sharma L, Bokov A, Beam H, et al. Mitochondrial metabolites extend lifespan. Aging Cell. 2016;15:336–348.

17. Munkacsy E, Khan MH, Lane RK, Borror MB, Park JH, Bokov AF, et al. DLK-1, SEK-3 and PMK-3 Are Required for the Life Extension Induced by Mitochondrial Bioenergetic Disruption in C. elegans. PLoS Genet. 2016;12:e1006133.

18. Qureshi MA, Haynes CM, Pellegrino MW. The mitochondrial unfolded protein response: Signaling from the powerhouse. J Biol Chem. 2017;292:13500–13506.

19. Kwak MK, Wakabayashi N, Kensler TW. Chemoprevention through the Keap1-Nrf2 signaling pathway by phase 2 enzyme inducers. Mutat Res. 2004;555:133–148.

20. Wood WB, ed. The Nematode Caenorhabditis elegans. New York: Cold Spring Harbor Laboratory; 1988.

21. Bhaskaran S, Butler JA, Becerra S, Fassio V, Girotti M, Rea SL. Breaking Caenorhabditis elegans the easy way using the Balch homogenizer: an old tool for a new application. Analytical Biochemistry. 2011;413:123–132.

22. Sleumer MC, Bilenky M, He A, Robertson G, Thiessen N, Jones SJ. Caenorhabditis elegans cisRED: a catalogue of conserved genomic elements. Nucleic Acids Res. 2009;37:1323–1334.

23. Huang da W, Sherman BT, Lempicki RA. Systematic and integrative analysis of large gene lists using DAVID bioinformatics resources. Nat Protoc. 2009;4:44–57.

24. Fabris F, Freitas AA. New KEGG pathway-based interpretable features for classifying ageing-related mouse proteins. Bioinformatics. 2016.

25. Barrett T, Wilhite SE, Ledoux P, Evangelista C, Kim IF, Tomashevsky M, et al. NCBI GEO: archive for functional genomics data sets--update. Nucleic Acids Res. 2013;41:D991–995.

26. Russell JC, Merrihew GE, Robbins JE, Postupna N, Kim T-K, Golubeva A, et al. Isolation and characterization of extracellular vesicles from Caenorhabditis elegans for multi-omic analysis. 2018;bioRxiv

27. Han SK, Lee D, Lee H, Kim D, Son HG, Yang JS, et al. OASIS 2: online application for survival analysis 2 with features for the analysis of maximal lifespan and healthspan in aging research. Oncotarget. 2016;7:56147–56152.

28. van der Vaart A, Rademakers S, Jansen G. DLK-1/p38 MAP Kinase Signaling Controls Cilium Length by Regulating RAB-5 Mediated Endocytosis in Caenorhabditis elegans. PLoS Genet. 2015;11:e1005733.

29. Wang Z, Harkins PC, Ulevitch RJ, Han J, Cobb MH, Goldsmith EJ. The structure of mitogen-activated protein kinase p38 at 2.1-A resolution. Proc Natl Acad Sci U S A. 1997;94:2327–2332.

30. Camps M, Nichols A, Arkinstall S. Dual specificity phosphatases: a gene family for control of MAP kinase function. FASEB J. 2000;14:6–16.

31. Zielinska DF, Gnad F, Jedrusik-Bode M, WisÃÅniewski JR, Mann M. Caenorhabditis elegans Has a Phosphoproteome Atypical for Metazoans That Is Enriched in Developmental and Sex Determination Proteins. Journal of Proteome Research. 2009.

32. Butler JA, Mishur RJ, Bokov AF, Hakala KW, Weintraub ST, Rea SL. Profiling the Anaerobic Response of C. elegans Using GC-MS. PLoS ONE. 2012;7:e46140.

33. Olmos Y, Carlton JG. The ESCRT machinery: new roles at new holes. Curr Opin Cell Biol. 2016;38:1–11.

34. Cocucci E, Meldolesi J. Ectosomes and exosomes: shedding the confusion between extracellular vesicles. Trends Cell Biol. 2015;25:364–372.

35. Tian Y, Garcia G, Bian Q, Steffen Kristan K, Joe L, Wolff S, et al. Mitochondrial Stress Induces Chromatin Reorganization to Promote Longevity and UPRmt. Cell. 2016.

36. Merkwirth C, Jovaisaite V, Durieux J, Matilainen O, Jordan Sabine D, Quiros Pedro M, et al. Two Conserved Histone Demethylases Regulate Mitochondrial Stress-Induced Longevity. Cell. 2016.

37. Schiavi A, Torgovnick A, Kell A, Megalou E, Castelein N, Guccini I, et al. Autophagy induction extends lifespan and reduces lipid content in response to frataxin silencing in C. elegans. Experimental Gerontology. 2013;48:191–201.

38. Yee C, Yang W, Hekimi S. The Intrinsic Apoptosis Pathway Mediates the Pro-Longevity Response to Mitochondrial ROS in C. elegans. Cell. 2014;157:897–909.

39. Ventura N, Rea SL, Schiavi A, Torgovnick A, Testi R, Johnson TE. p53/CEP-1 increases or decreases lifespan, depending on level of mitochondrial bioenergetic stress. Aging Cell. 2009;8:380–393.

40. Walter L, Baruah A, Chang HW, Pace HM, Lee SS. The homeobox protein CEH-23 mediates prolonged longevity in response to impaired mitochondrial electron transport chain in C. elegans. PLoS Biology. 2011;9:e1001084.

41. Durieux J, Wolff S, Dillin A. The Cell-Non-Autonomous Nature of Electron Transport Chain-Mediated Longevity. Cell. 2011;144:79–91.

42. Shao LW, Niu R, Liu Y. Neuropeptide signals cell non-autonomous mitochondrial unfolded protein response. Cell Res. 2016;26:1182–1196.

43. Bennett CF, Vander Wende H, Simko M, Klum S, Barfield S, Choi H, et al. Activation of the mitochondrial unfolded protein response does not predict longevity in Caenorhabditis elegans. Nat Commun. 2014;5:3483.

44. Jeong DE, Lee D, Hwang SY, Lee Y, Lee JE, Seo M, et al. Mitochondrial chaperone HSP-60 regulates anti-bacterial immunity via p38 MAP kinase signaling. EMBO J. 2017.

45. Wehman AM, Poggioli C, Schweinsberg P, Grant BD, Nance J. The P4-ATPase TAT-5 inhibits the budding of extracellular vesicles in C. elegans embryos. Curr Biol. 2011;21:1951–1959.

46. Savina A, Vidal M, Colombo MI. The exosome pathway in K562 cells is regulated by Rab11. J Cell Sci. 2002;115:2505–2515.

47. Christ L, Raiborg C, Wenzel EM, Campsteijn C, Stenmark H. Cellular Functions and Molecular Mechanisms of the ESCRT Membrane-Scission Machinery. Trends Biochem Sci. 2017;42:42–56.

48. Scourfield EJ, Martin-Serrano J. Growing functions of the ESCRT machinery in cell biology and viral replication. Biochem Soc Trans. 2017;45:613–634.

49. Zhang Y, Kim MS, Jia B, Yan J, Zuniga-Hertz JP, Han C, et al. Hypothalamic stem cells control ageing speed partly through exosomal miRNAs. Nature. 2017;advance online publication.

50. Grigorian-Shamagian L, Liu W, Fereydooni S, Middleton RC, Valle J, Hyung Cho J, et al. Cardiac and systemic rejuvenation after cardiosphere-derived cell therapy in senescent rats. European Heart Journal. 2017; ehx454.

51. Takasugi M, Okada R, Takahashi A, Virya Chen D, Watanabe S, Hara E. Small extracellular vesicles secreted from senescent cells promote cancer cell proliferation through EphA2. Nat Commun. 2017;8:15729.

52. Schiavi A, Maglioni S, Palikaras K, Shaik A, Strappazzon F, Brinkmann V, et al. Iron-Starvation-Induced Mitophagy Mediates Lifespan Extension upon Mitochondrial Stress in C. elegans. Curr Biol. 2015;25:1810–1822.

53. Chikka MR, Anbalagan C, Dvorak K, Dombeck K, Prahlad V. The Mitochondria-Regulated Immune Pathway Activated in the C. elegans Intestine Is Neuroprotective. Cell Rep. 2016;16:2399–2414.

54. Egan DF, Shackelford DB, Mihaylova MM, Gelino S, Kohnz RA, Mair W, et al. Phosphorylation of ULK1 (hATG1) by AMP-activated protein kinase connects energy sensing to mitophagy. Science. 2011;331:456–461.

55. Park EC, Glodowski DR, Rongo C. The ubiquitin ligase RPM-1 and the p38 MAPK PMK-3 regulate AMPA receptor trafficking. PLoS ONE. 2009;4:e4284.

56. D’Souza SA, Rajendran L, Bagg R, Barbier L, van Pel DM, Moshiri H, et al. The MADD-3 LAMMER Kinase Interacts with a p38 MAP Kinase Pathway to Regulate the Display of the EVA-1 Guidance Receptor in Caenorhabditis elegans. PLoS Genet. 2016;12:e1006010.

57. Cutter AD. Divergence times in Caenorhabditis and Drosophila inferred from direct estimates of the neutral mutation rate. Mol Biol Evol. 2008;25:778–786.

58. Nargund AM, Pellegrino MW, Fiorese CJ, Baker BM, Haynes CM. Mitochondrial import efficiency of ATFS-1 regulates mitochondrial UPR activation. Science. 2012;337:587–590.

