## Supplementary figures and images for "The PMK-3 (p38) Mitochondrial Retrograde Response Functions in Intestinal Cells to Extend Life via the ESCRT Machinery"

### Supplemental Figure S1

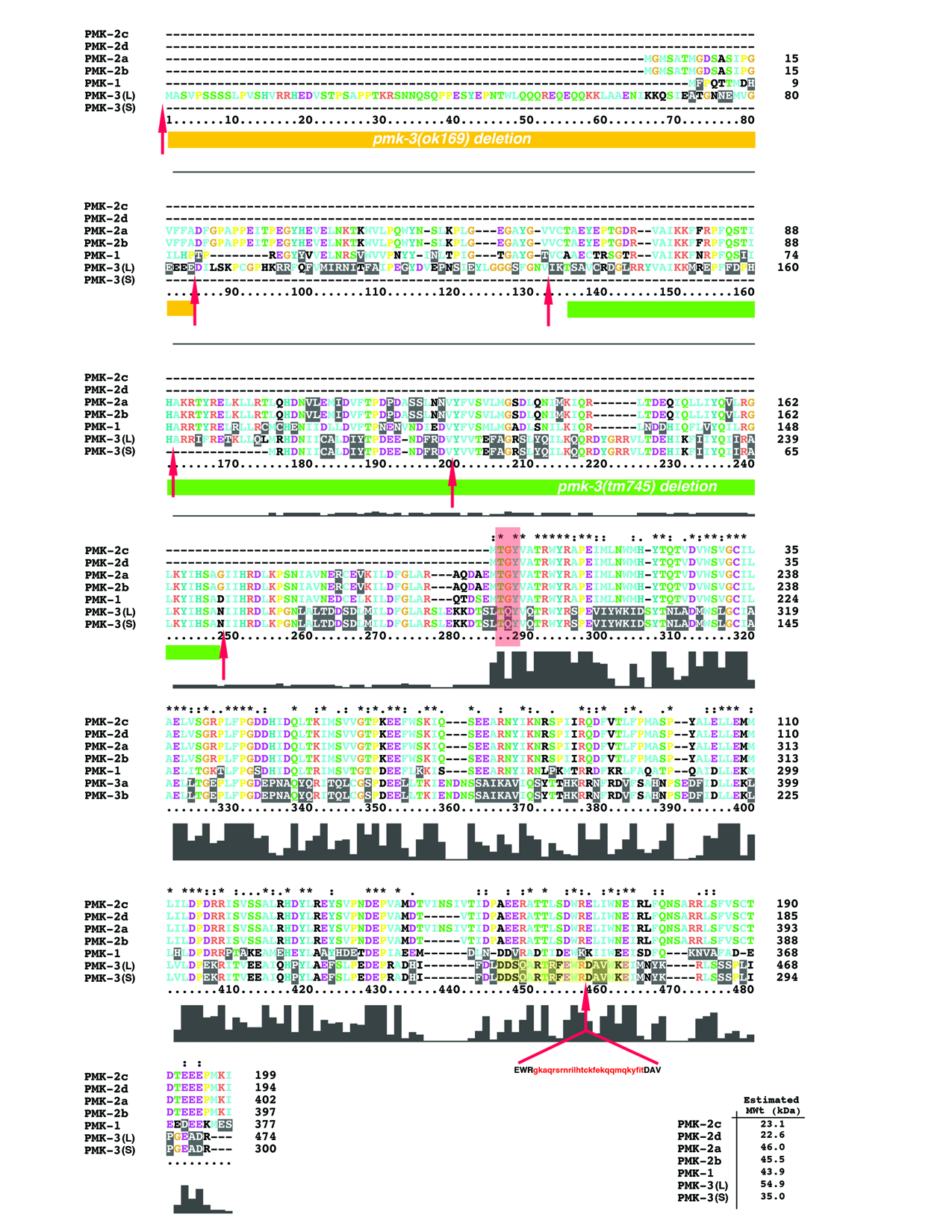

### Supplemental Figure S2

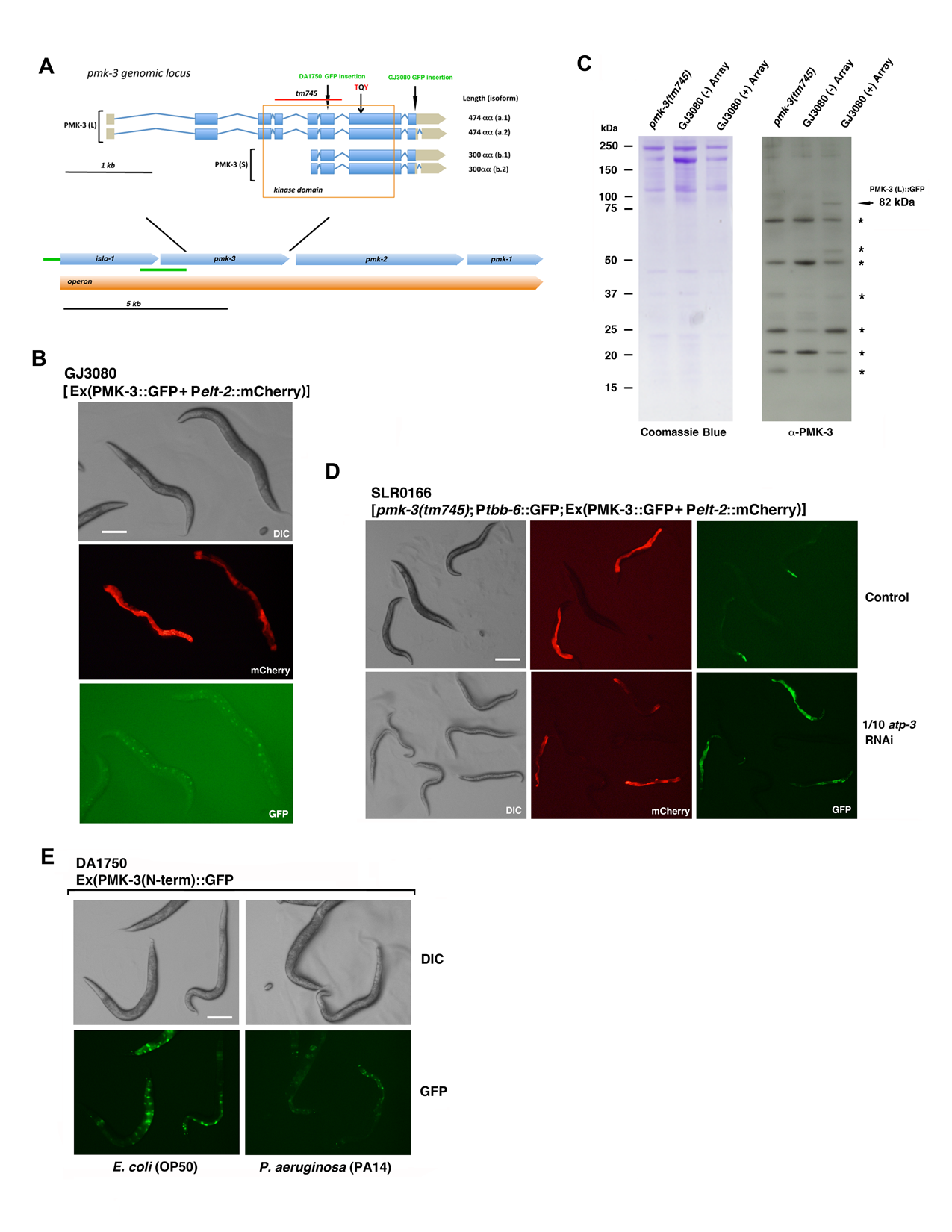

### Supplemental Figure S3

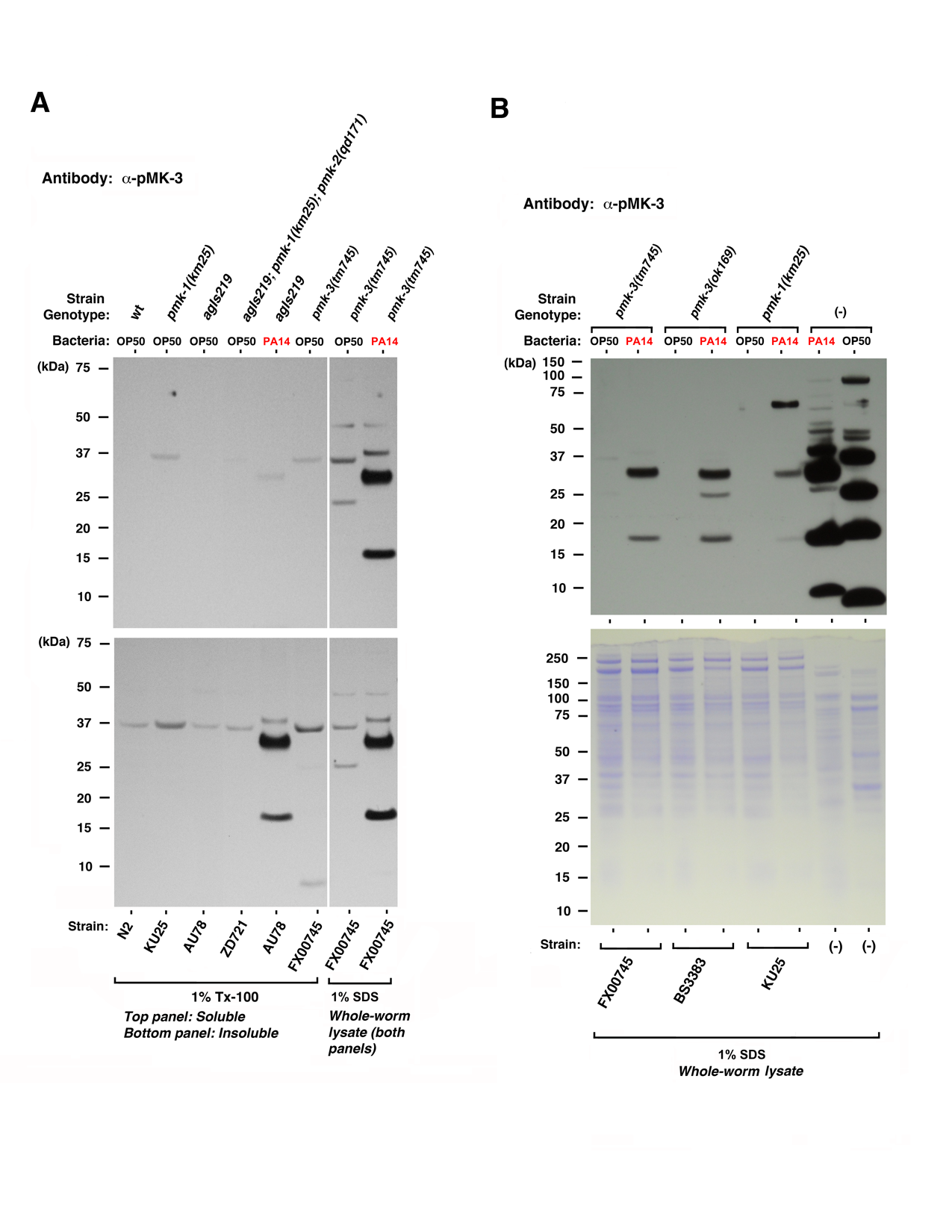

### Supplemental Figure S4

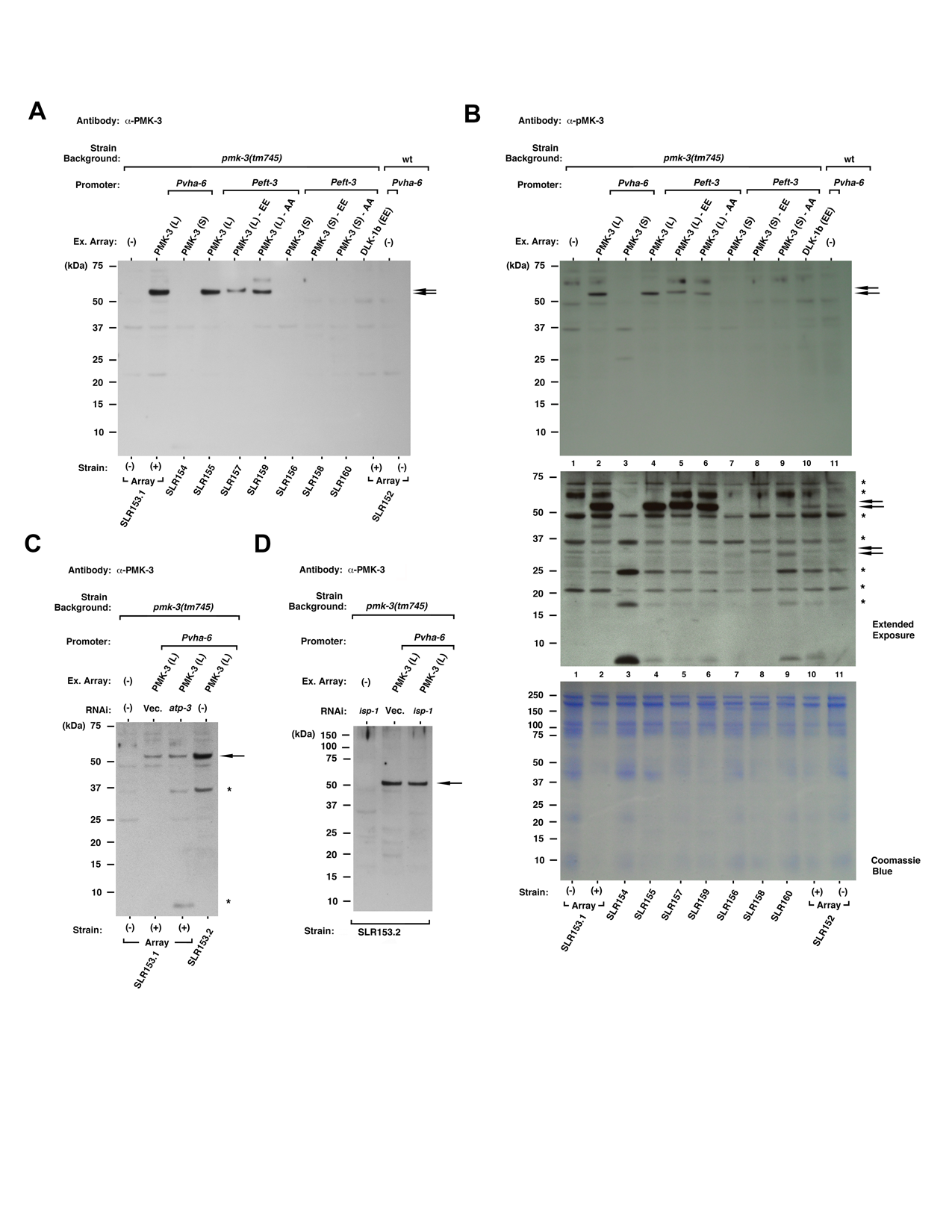

### Supplemental Figure S5

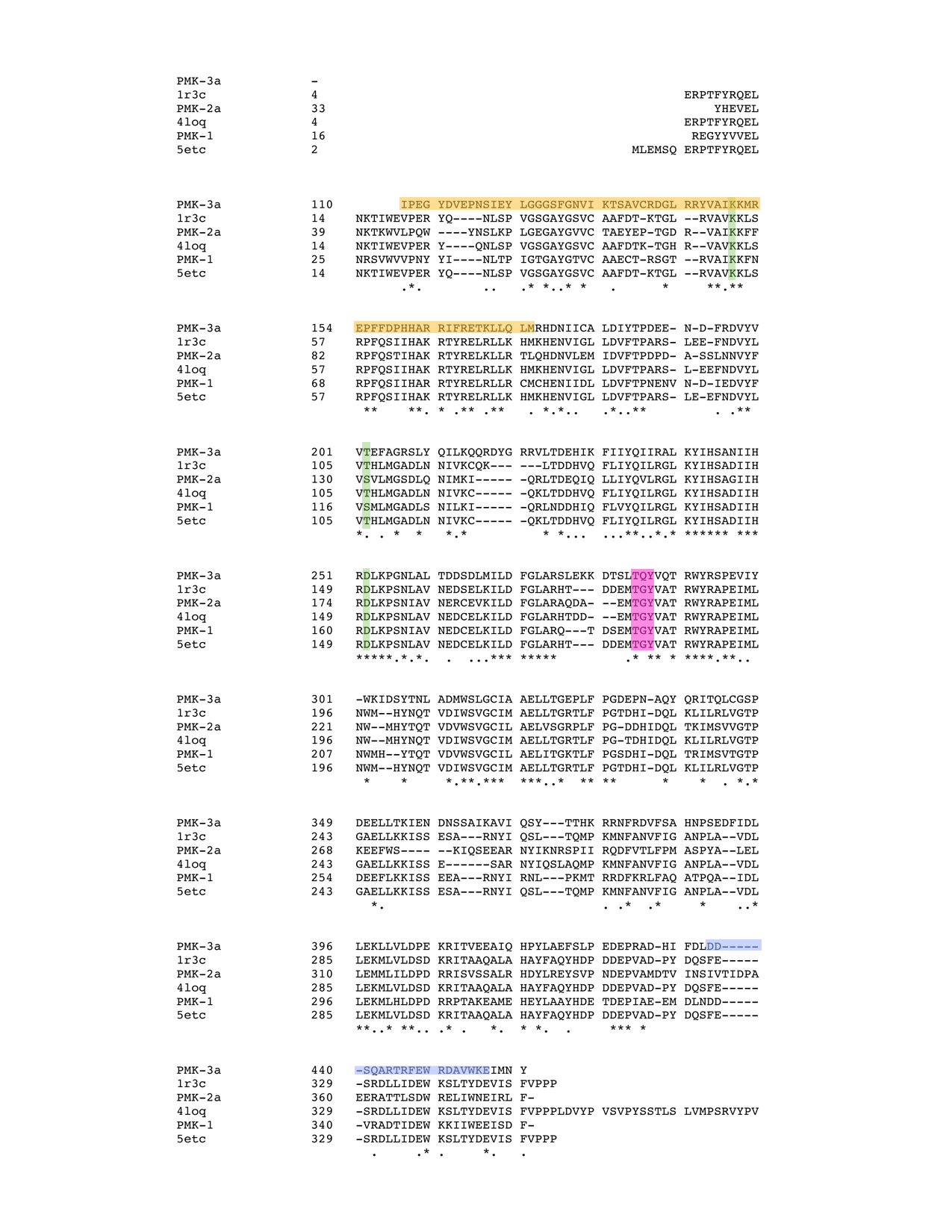

### Supplemental Figure S6

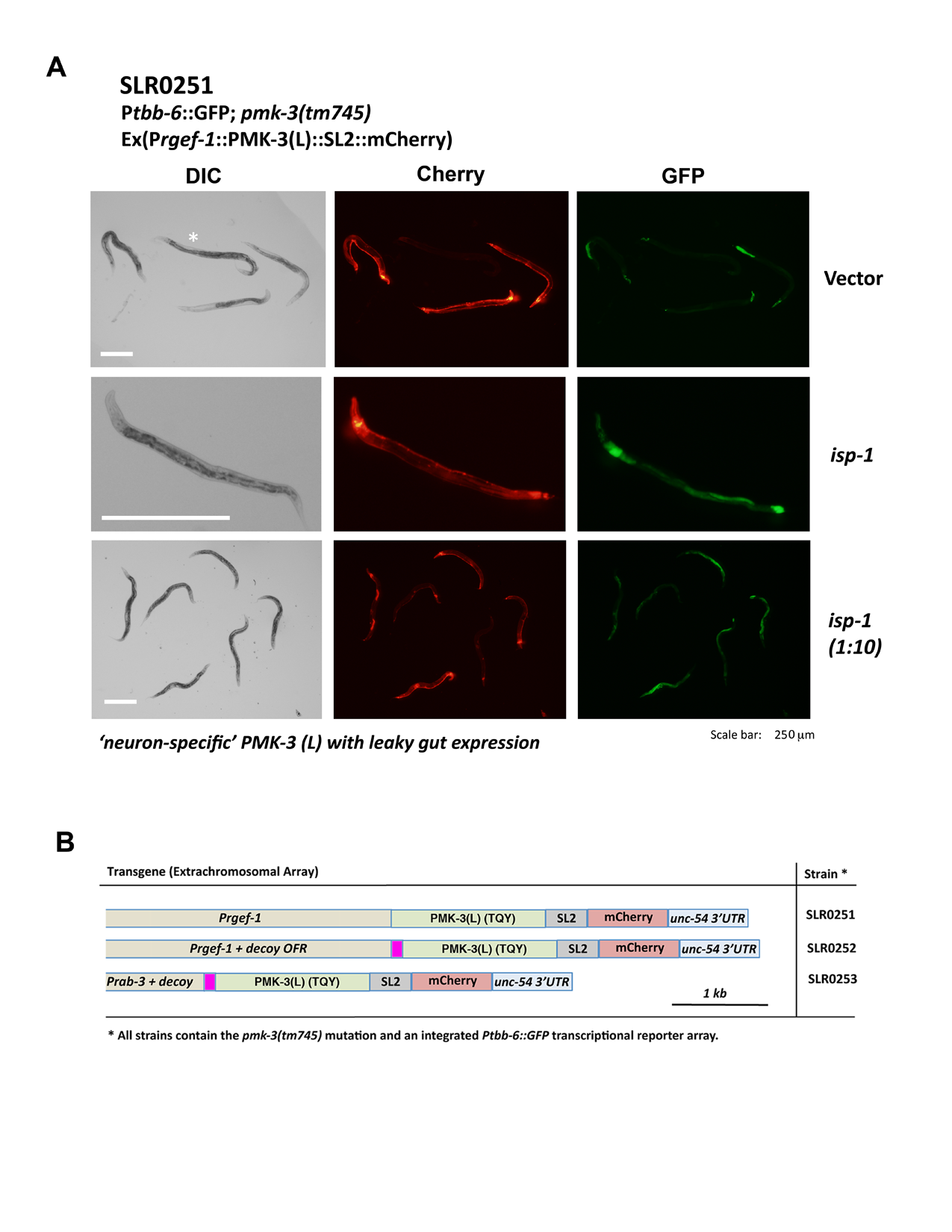

### Supplemental Figure S7

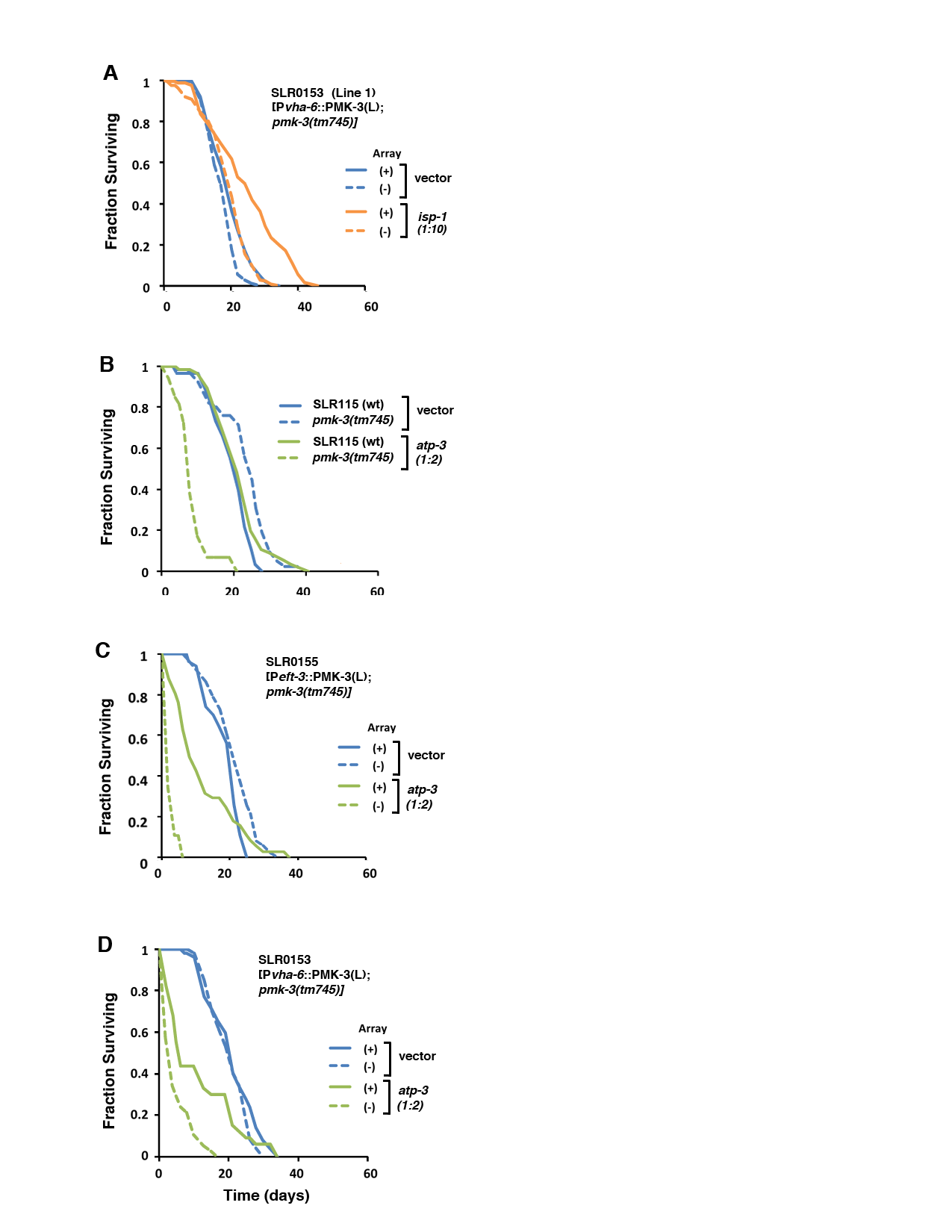

### Supplemental Figure S8

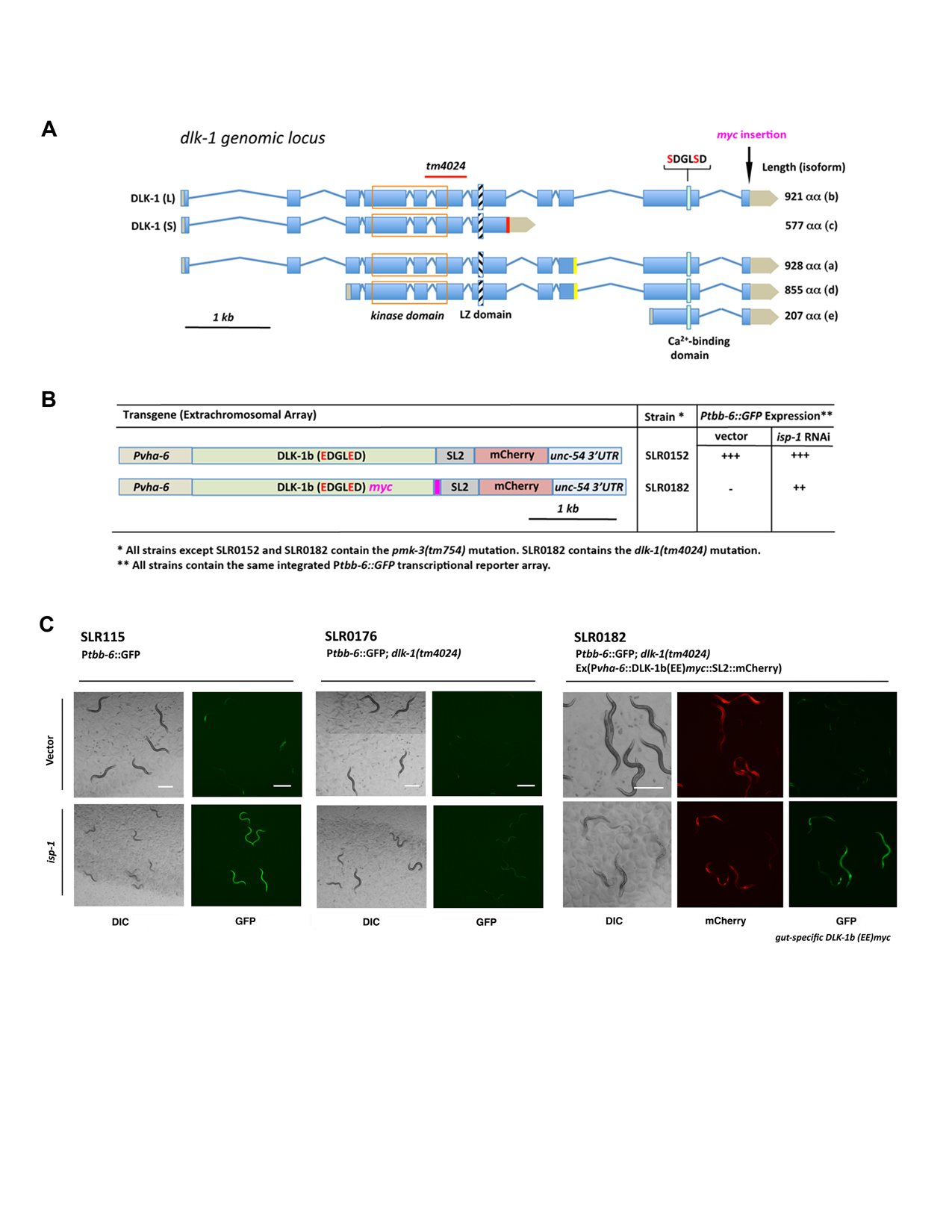

### Supplemental Figure S9

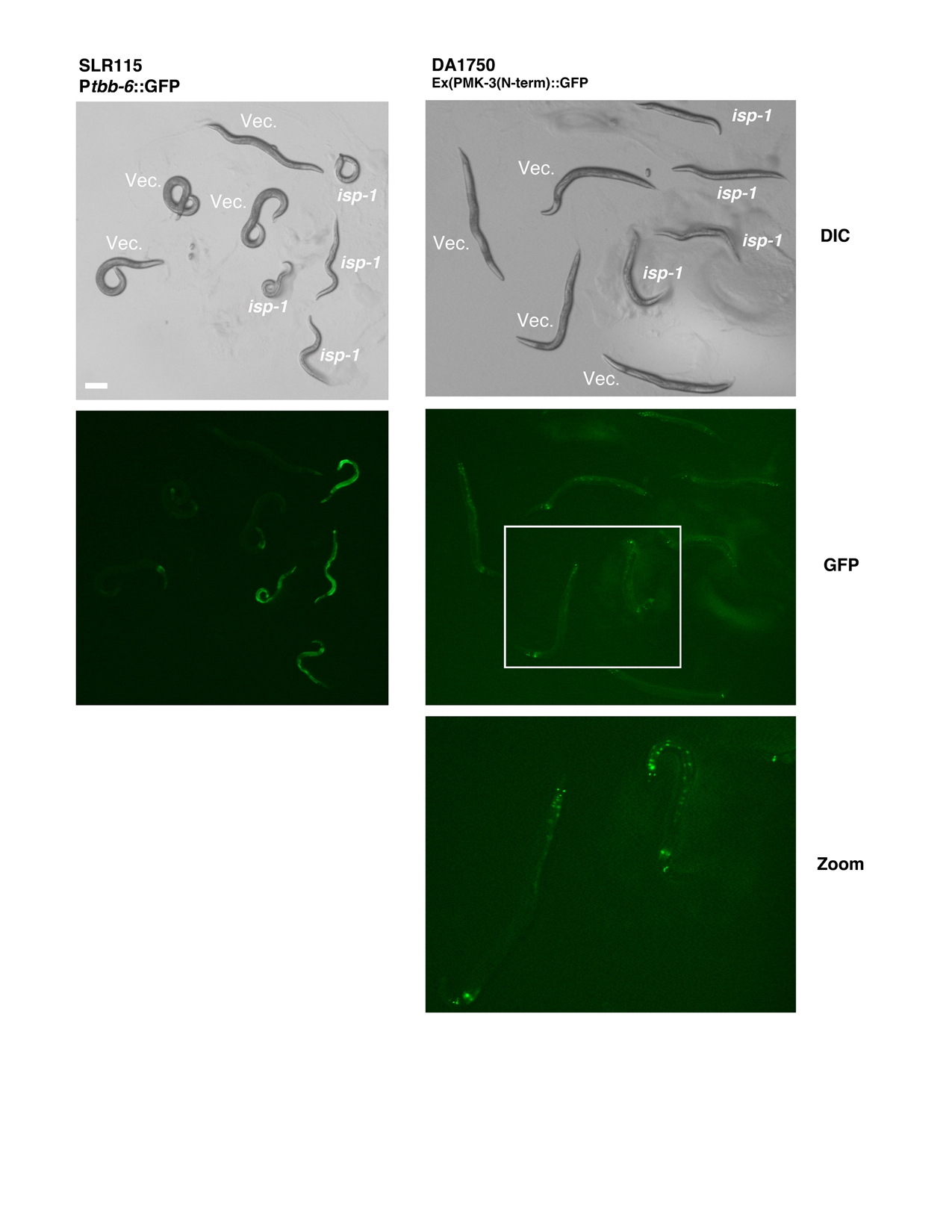

### Supplemental Figure S10

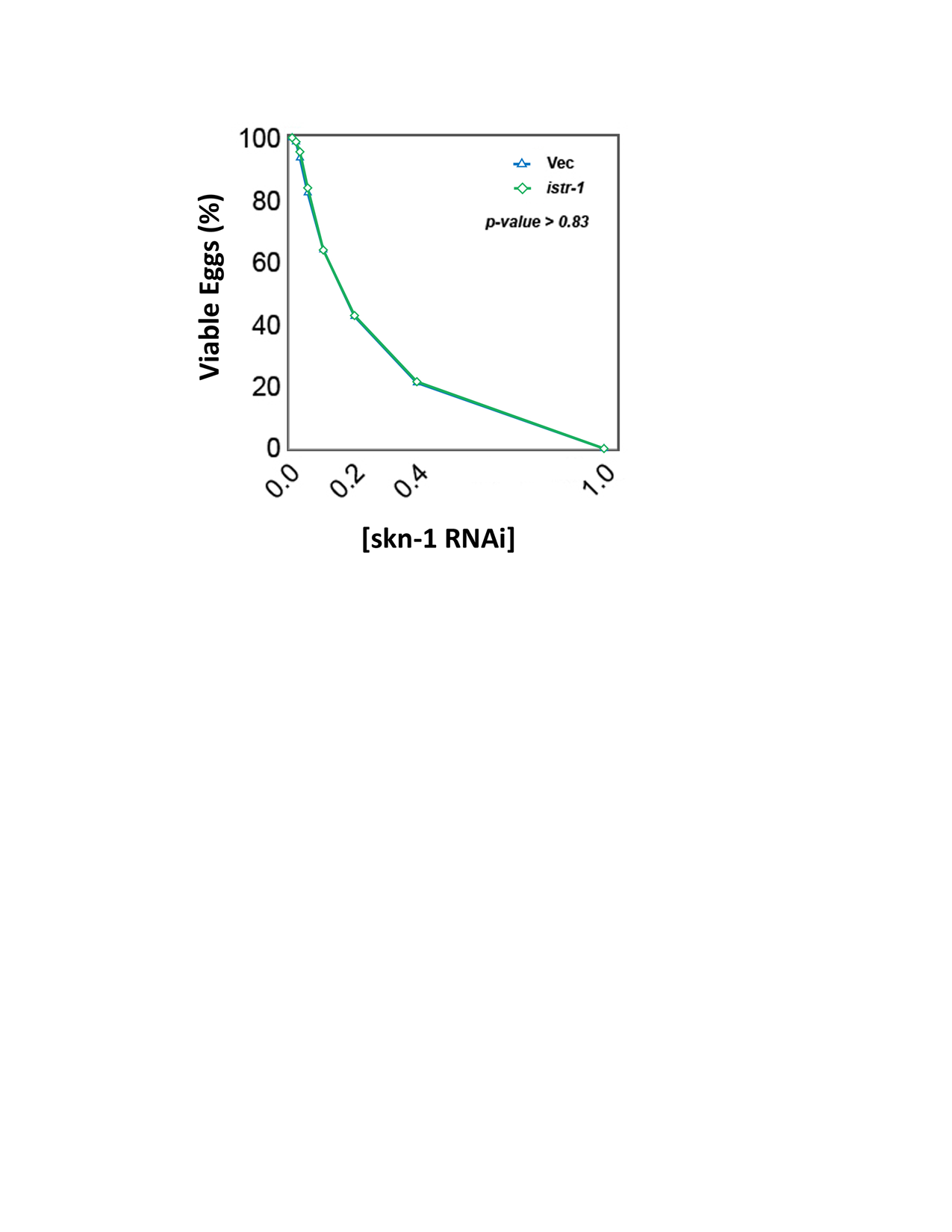

### Supplemental Figure S11

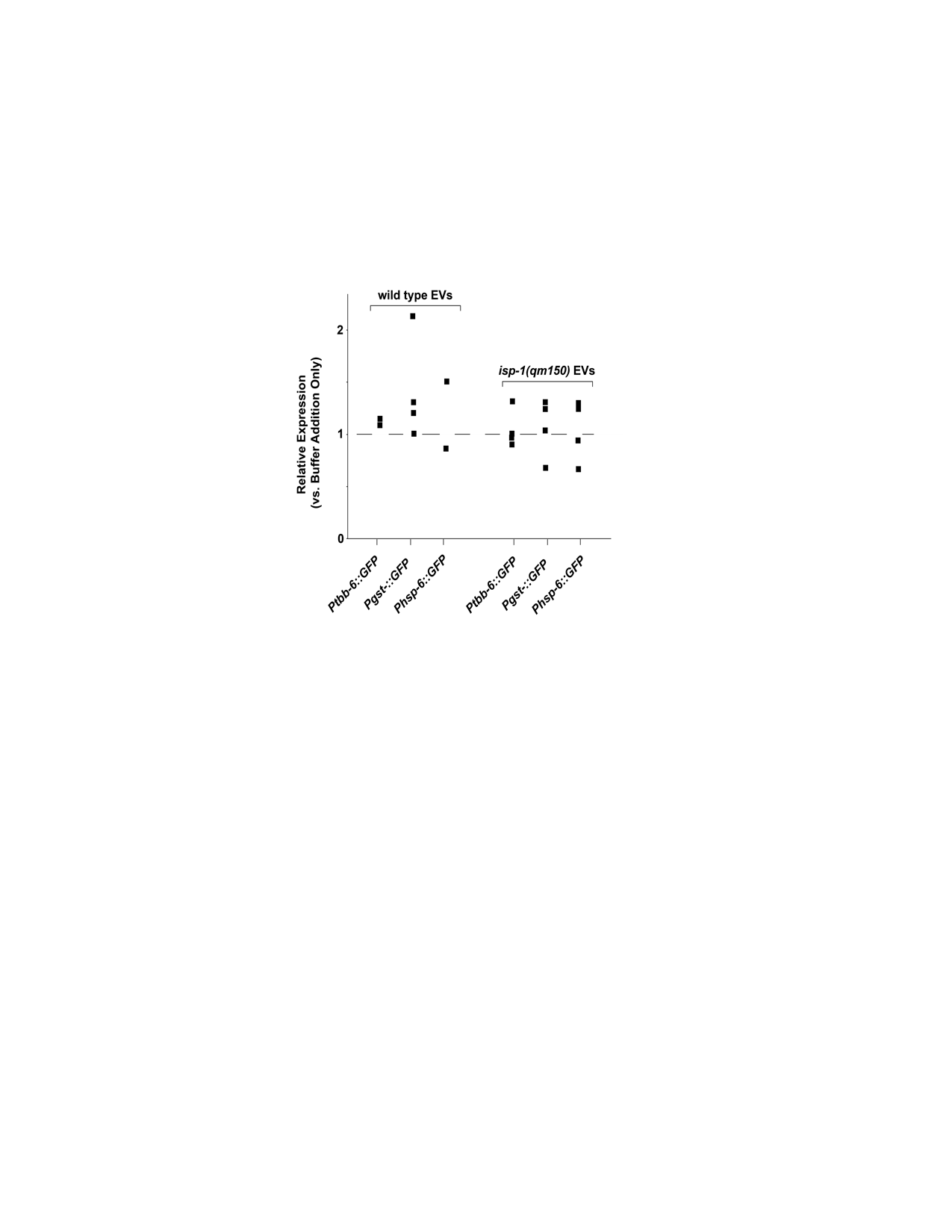
