## Supplemental Table S1 for "The PMK-3 (p38) Mitochondrial Retrograde Response Functions in Intestinal Cells to Extend Life via the ESCRT Machinery"

**Table S1: Strain List**

| Strain | Genotype* | Source | Strain Construction** |
|---|---|---|---|
| AU78*** | *agIs219 [T24B8.5p::GFP::unc-54-3' UTR + ttx-3p::GFP::unc-54-3' UTR] III* | CGC | *Shivers, RP, et. al. PLoS Genet. 2010 Apr 1;6(4):e1000892* |
| BS3383 | *pmk-3(ok169) IV* | CGC | |
| CL2166 | *dvIs19[pAF15(Pgst-4::GFP::NLS)] III* | CGC | |
| CL3462 | *smg-1(cc546) I; dvIs67[pCL179(Ptbb-6::GFP) + pCL148(Pmyo-3::dsRed monomer)] III* | Dr. Christopher Link, UC Boulder | *Munkacsy E, et. al. PLoS Genet. 2016;12(7):e1006133* |
| DA1750 | *adEx1750 [pmk-3::GFP + rol-6(su1006)]* | CGC | *Berman K, et.al. Mol Cell Biol Res Commun. 2001;4(6):337-44* |
| FX00745 | *pmk-3(tm745) IV* | Dr. Shohei Mitani, National Bioresource Project (Japan) | |
| FX0424 | *dlk-1(tm4024) I* | GCG | |
| GJ3080 | *gjEx1863[pmk-3::gfp + Pelt-2::cherry]* | Dr. Gert Jansen, Erasmus MC (Netherlands) | *van der Vaart A, et. al. PLoS Genet. 2015;11(12):e1005733* |
| HT1593 | *unc-119(ed3) III* | CGC | |
| KU25 | *pmk-1(km25) IV* | CGC | |
| N2 (Bristol) | wild-type | CGC | |
| SJ4100 | *zcIs13[Phsp-6::GFP] V* | CGC | Outcrossed 5 x with N2 |
| SLR0115 | *dvIs67[pCL179(Ptbb-6::GFP) + pCL148(Pmyo-3::dsRed monomer)] III* | Rea Lab. | *Munkacsy E, et. al. PLoS Genet. 2016;12(7):e1006133* |
| SLR0150 | *pmk-3(tm745) IV; dvIs67[pCL179(Ptbb-6::GFP) + pCL148(Pmyo-3::dsRed monomer)] III* | Rea Lab. | *Munkacsy E, et. al. PLoS Genet. 2016;12(7):e1006133* |
| SLR0152 | *smg-1(cc546) I; dvIs67[pCL179(Ptbb-6::GFP) + pCL148(Pmyo-3::dsRed monomer)] III; stxEx6 [pSR1(Pvha-6::dlk-1 b (EE)::SL2::mCherry)]* | This study | Plasmid pSR1 microinjected into CL3462 (20ng/uL) |
| SLR0153 | *pmk-3(tm745) IV; dvIs67[pCL179(Ptbb-6::GFP) + pCL148(Pmyo-3::dsRed monomer)] III; stxEx7 [pSR2(Pvha-6::pmk-3 L::SL2::mCherry)]* | This study | Plasmid pSR2 microinjected into SLR150 (20ng/uL) |
| SLR0154 | *pmk-3(tm745) IV; dvIs67[pCL179(Ptbb-6::GFP) + pCL148(Pmyo-3::dsRed monomer)] III; stxEx8 [pSR3(Pvha-6::pmk-3 S::SL2::mCherry)]* | This study | Plasmid pSR3 microinjected into SLR150 (20ng/uL) |
| SLR0155 | *pmk-3(tm745) IV; dvIs67[pCL179(Ptbb-6::GFP) + pCL148(Pmyo-3::dsRed monomer)] III; stxEx9 [pSR4(Peft-3::pmk-3 L::SL2::mCherry)]* | This study | Plasmid pSR4 microinjected into SLR150 (20ng/uL) |
| SLR0156 | *pmk-3(tm745) IV; dvIs67[pCL179(Ptbb-6::GFP) + pCL148(Pmyo-3::dsRed monomer)] III; stxEx10 [pSR5(Peft-3::pmk-3 S::SL2::mCherry)]* | This study | Plasmid pSR5 microinjected into SLR150 (20ng/uL) |
| SLR0157 | *pmk-3(tm745) IV; dvIs67[pCL179(Ptbb-6::GFP) + pCL148(Pmyo-3::dsRed monomer)] III; stxEx11 [pSR6(Peft-3::pmk-3 L (EE)::SL2::mCherry)]* | This study | Plasmid pSR6 microinjected into SLR150 (20ng/uL) |
| SLR0158 | *pmk-3(tm745) IV; dvIs67[pCL179(Ptbb-6::GFP) + pCL148(Pmyo-3::dsRed monomer)] III; stxEx12 [pSR7(Peft-3::pmk-3 S (EE)::SL2::mCherry)]* | This study | Plasmid pSR7 microinjected into SLR150 (20ng/uL) |
| SLR0159 | *pmk-3(tm745) IV; dvIs67[pCL179(Ptbb-6::GFP) + pCL148(Pmyo-3::dsRed monomer)] III; stxEx13 [pSR8(Peft-3::pmk-3 L (AA)::SL2::mCherry)]* | This study | Plasmid pSR8 microinjected into SLR150 (20ng/uL) |
| SLR0160 | *pmk-3(tm745) IV; dvIs67[pCL179(Ptbb-6::GFP) + pCL148(Pmyo-3::dsRed monomer)] III; stxEx14 [pSR9(Peft-3::pmk-3 S (AA)::SL2::mCherry)]* | This study | Plasmid pSR9 microinjected into SLR150 (20ng/uL) |
| SLR0161 | *stxEx6 [pSR1(Pvha-6::dlk-1 L (EE)::SL2::mCherry)]* | This study | SLR0152 x N2 (note: *smg-1* allele uncharacterized) |
| SLR0162 | *pmk-3(tm745) IV; dvIs67[pCL179(Ptbb-6::GFP) + pCL148(Pmyo-3::dsRed monomer)] III; stxEx6 [pSR1(Pvha-6::dlk-1 b (EE)::SL2::mCherry)]* | This study | SLR0161 x SLR0150; SLR0150 x F1 male progeny |
| SLR0164 | *pmk-3(tm745) IV; dvIs67[pCL179(Ptbb-6::GFP) + pCL148(Pmyo-3::dsRed monomer)] III; stxEx6 [pSR1(Pvha-6::dlk-1 b(EE)::SL2::mCherry)]; stxEx7 [pSR2(Pvha-6::pmk-3 L::SL* | This study | SLR0162 x SLR0153 |
| SLR0166 | *pmk-3(tm745) IV; dvIs67[pCL179(Ptbb-6::GFP) + pCL148(Pmyo-3::dsRed monomer)] III; gjEx1863[pmk-3::gfp + Pelt-2::cherry]* | This study | GJ3080 x N2; SLR0153 x F1 male progeny |
| SLR0176 | *dlk-1(tm4024) I; dvIs67[pCL179(Ptbb-6::GFP) + pCL148(Pmyo-3::dsRed monomer)] III* | This study | FX0424 x SLR115 |
| SLR0182 | *stxEx54 [pSR16(Pvha-6::DLK-1b(EE)::myc::SL2::mCherry)]; dlk-1(tm4024) I; dvIs67[pCL179(Ptbb-6::GFP) + pCL148(Pmyo-3::dsRed monomer)] III* | This study | Plasmid pSR16 microinjected into SLR0176 (50ng/uL) |
| SLR0246 | *unc-119(ed3) III; stxEx39 [pSR10(alx-1::GFP]* | Source BioScience (CBGtg9050F0275D) & this study | Rescued Fosmid (pSR10) microinjected into HT1593 (70ng/uL) |
| SLR0247 | *unc-119(ed3) III; stxEx48 [pSR11(C01A4.2)* | Source BioScience (CBGtg9050D12203D) & this study | Rescued Fosmid (pSR11) microinjected into HT1593 (95ng/uL) |
| SLR0248 | *unc-119(ed3) III; stxEx35 [pSR12(istr-1::GFP]* | Source BioScience (CBGtg9050D1061D) & this study | Rescued Fosmid (pSR12) microinjected into HT1593 (75ng/uL) |
| SLR0251 | *pmk-3(tm745) IV; dvIs67[pCL179(Ptbb-6::GFP) + pCL148(Pmyo-3::dsRed monomer)] III; stxEx27 [pSR13(Prgef-1::PMK-3(L)::SL2::mCherry)]* | This study | Plasmid pSR13 microinjected into SLR0150 (5 ng/uL + 50ng/uL pL4440 (filler DNA)) |
| SLR0252 | *pmk-3(tm745) IV; dvIs67[pCL179(Ptbb-6::GFP) + pCL148(Pmyo-3::dsRed monomer)] III; stxEx49 [pSR14(Prgef-1::(decoyATG)::PMK-3(L)::SL2::mCherry)]* | This study | Plasmid pSR14 microinjected into SLR0150 (5 ng/uL + 50 ng/uL Invitrogen Cat. no 15615-016) |
| SLR0253 | *pmk-3(tm745) IV; dvIs67[pCL179(Ptbb-6::GFP) + pCL148(Pmyo-3::dsRed monomer)] III; stxEx50 [pSR15(Prab-3::(decoyATG)::PMK-3(L)::SL2::mCherry)]* | This study | Plasmid pSR15 microinjected into SLR0150 (5 ng/uL + 50 ng/uL Invitrogen Cat. no 15615-016) |
| ZD721 | *agIs219 [T24B8.5p::GFP::unc-54-3' UTR + ttx-3p::GFP::unc-54-3' UTR] III; pmk-1(km25) IV; pmk-2(qd171) IV* | Dr. Dennis Kim, MIT (USA) | *Pagano DJ et.al. PLoS Genet. 2015;11(2):e1004997* |

* For all transcriptional reporters, genes contributing the promoter are prefixed with P. Many strains were co-injected with "stuffer" DNA to minimize array silencing. Full details are provided in S1 Text.

** For crosses, the hermaphrodite strain is written first, followed by the male stain. Sequential crosses are separated by a semicolon.

*** T24B8.5 is regulated by PMK-1
