## Supplemental Table S2 for "The PMK-3 (p38) Mitochondrial Retrograde Response Functions in Intestinal Cells to Extend Life via the ESCRT Machinery"

### Study #1 (Corresponds to Figs. 3 and 4C)

**Control Strains: (n.b. only relevant genotype information is provided. For Full genotypic information refer to Table S1.)**

**1.SLR115 [Ptbb-6::GFP]**
**2. *pmk-3(tm745)***

**Test Strains**
**3. SLR152 [*Pvha-6::dlk-1 b (EE)::SL2::mCherry*; Ptbb-6::GFP]**
**4. SLR152 Sans Array [*Ptbb-6::GFP*]**

**5. SLR155 [*Peft-3::pmk3-L::SL2::mCherry; Ptbb-6::GFP; pmk-3(tm745)*]**
**6. SLR155 Sans Array [*Ptbb-6::GFP; pmk-3(tm745)*]**

**7. SLR153.2* [*Pvha-6::pmk-3-L::SL2::mCherry; Ptbb-6::GFP; pmk-3(tm745)*]**
**8. SLR153.2 Sans Array [*Ptbb-6::GFP; pmk-3(tm745)*]**
*** SLR0153 Line 2**

**Descriptive Statistics**

| Strain | Condition | Sample Size (censored events) | Expected Life Span (days) |
|---|---|---|---|
| **Control Strains** | | | |
| **1. SLR115** | pL4440 *(empty vector)* | 68 (6) | 19.5 |
| | *isp-1 (1:10)* | 70 (6) | 27.7 |
| **2. *pmk-3(tm745)*** | pL4440 (empty vector) | 69 (6) | 20.2 |
| | *isp-1 (1:10)* | 70 (7) | 17.5 |
| **Test Strains** | | | |
| **3. SLR152** | pL4440 *(empty vector)* | 59 (7) | 17.8 |
| | *isp-1 (1:10)* | 70 (11) | 29.8 |
| **4. SLR152 Sans Array** | pL4440 *(empty vector)* | 69 (25) | 20.1 |
| | *isp-1 (1:10)* | 69 (10) | 26.7 |
| **5. SLR155** | pL4440 *(empty vector)* | 67 (16) | 17.2 |
| | *isp-1 (1:10)* | 70 (13) | 24.0 |
| **6. SLR155 Sans Array** | pL4440 *(empty vector)* | 71 (11) | 18.7 |
| | *isp-1 (1:10)* | 70 (6) | 20.5 |
| **7. SLR153.2** | pL4440 *(empty vector)* | 70 (15) | 21.3 |
| | *isp-1 (1:10)* | 73 (7) | 27.2 |
| **8. SLR153 Sans Array** | pL4440 *(empty vector)* | 70 (10) | 18.2 |
| | *isp-1 (1:10)* | 70 (12) | 20.6 |

**Log Rank Analyses**

| Within Strain Comparisons | *Long Rank Test (p-value*)* |
|---|---|
| **Control Strains** | |
| **1. SLR115** | |
| pL4440 vs *isp-1 (1:10)* | 2.77E-14 |
| **2. *pmk-3(tm745)*** | |
| pL4440 vs *isp-1 (1:10)* | 1.03E-01 |
| **Test Strains** | |
| **3. SLR152** | |
| pL4440 vs *isp-1 (1:10)* | 6.82E-12 |
| **4. SLR152 Sans Array** | |
| pL4440 vs *isp-1 (1:10)* | 1.26E-05 |
| **5. SLR155** | |
| pL4440 vs *isp-1 (1:10)* | 1.92E-06 |
| **6. SLR155 Sans Array** | |
| pL4440 vs *isp-1 (1:10)* | 4.30E-02 |
| **7. SLR153.2** | |
| pL4440 vs *isp-1 (1:10)* | 6.16E-05 |
| **8. SLR153.2 Sans Array** | |
| pL4440 vs *isp-1 (1:10)* | 2.88E-02 |

* Red = significant after correction for multiple testing (Bonferroni Correction) p < 0.05/20 or 0.0025)

**Log Rank Analyses Contd.**

| ***Strains & Conditions*** | ***Long Rank Test (p-value*)*** |
|---|---|
| ***1. SLR115 vs. pmk-3(tm745)*** | |
| pL4440 | 6.80E-01 |
| *isp-1 (1:10)* | 6.64E-12 |
| ***2. SLR152 w/w.o. Array*** | |
| pL4440 | 1.12E-02 |
| *isp-1 (1:10)* | 2.21E-02 |
| ***3. SLR155 w/w.o. Array*** | |
| pL4440 | 2.23E-01 |
| *isp-1 (1:10)* | 4.90E-02 |
| ***4. SLR153.2 w/w.o. Array*** | |
| pL4440 | 5.32E-03 |
| *isp-1 (1:10)* | 7.49E-05 |
| ***5. SLR152 (+) Array vs. SLR115*** | |
| pL4440 | 3.13E-02 |
| *isp-1 (1:10)* | 1.93E-02 |
| ***6. SLR155 (+) Array vs. SLR153.2 (+) Array*** | |
| pL4440 | 9.69E-04 |
| *isp-1 (1:10)* | 6.05E-02 |

* Red = significant after correction for multiple testing (Bonferroni Correction) p < 0.05/20 or 0.0025)

### Study #2 ( (Corresponds to Figs. 3, 4C and Fig. S7C-D)

**Control Strains**

**1.SLR115 [Ptbb-6::GFP]**

**2. *pmk-3(tm745)***

**Test Strains**

**3. SLR152 [*Pvha-6::dlk-1 b (EE)::SL2::mCherry;* Ptbb-6::GFP]**

**4. SLR152 Sans Array [*Ptbb-6::GFP*]**

**5. SLR155 [*Peft-3::pmk3-L::SL2::mCherry; Ptbb-6::GFP; pmk-3(tm745)*]**

**6. SLR155 Sans Array [*Ptbb-6::GFP; pmk-3(tm745)*]**

**7. SLR153.2* [*Pvha-6::pmk-3-L::SL2::mCherry; Ptbb-6::GFP; pmk-3(tm745)*]**

**8. SLR153.2 Sans Array [*Ptbb-6::GFP; pmk-3(tm745)*]**

*** SLR0153 Line 2**

**Descriptive Statistics**

| Strain | Condition | Sample Size (censored events) | Expected Life Span (days) |
|---|---|---|---|
| **Control Strains** | | | |
| **1. SLR115** | 4440 (empty vecto | 61 (5) | 19.8 |
| | *atp-3 (1:2)* | 60 (4) | 21.5 |
| | *isp-1 (1:2)* | 60 (12) | 36.0 |
| ***2. pmk-3(tm745)*** | 4440 (empty vecto | 60 (15) | 23.3 |
| | *atp-3 (1:2)* | 60 (28) | 8.7 |
| | *isp-1 (1:2)* | 60 (11) | 27.1 |
| **Test Strains** | | | |
| **3. SLR152** | 4440 (empty vecto | 60 (1) | 21.6 |
| | *isp-1 (1:2)* | 60 (5) | 34.7 |
| **4. SLR152 Sans A** | 4440 (empty vecto | 60 (3) | 21.7 |
| | *isp-1 (1:2)* | 60 (3) | 35.5 |
| **5. SLR155** | 4440 (empty vecto | 60 (11) | 18.8 |
| | *atp-3 (1:2)* | 62 (16) | 12.4 |
| | *isp-1 (1:2)* | 61 (11) | 34.1 |
| **6. SLR155 Sans A** | 4440 (empty vecto | 60 (9) | 21.4 |
| | *atp-3 (1:2)* | 60 (10) | 2.8 |
| | *isp-1 (1:2)* | 60 (11) | 23.3 |
| **7. SLR153.2** | 4440 (empty vecto | 60 (9) | 21.0 |
| | *atp-3 (1:2)* | 60 (17) | 11.6 |
| | *isp-1 (1:2)* | 56 (6) | 35.7 |
| **8. SLR153.2 Sans** | 4440 (empty vecto | 60 (13) | 20.3 |
| | *atp-3 (1:2)* | 63 (12) | 5.0 |
| | *isp-1 (1:2)* | 60 (15) | 19.5 |

**Log Rank Analyses**

| Within Strain Comparisons | | | |
|---|---|---|---|
| **Control Strains** | | | ***Long Rank Test (p-value*)*** |
| **1. SLR115** | | | |
| pL4440 | vs | *atp-3 (1:2)* | 1.09E-01 |
| pL4440 | vs | *isp-1 (1:2)* | 3.12E-14 |
| ***2. pmk-3(tm745)*** | | | |
| pL4440 | vs | *atp-3 (1:2)* | 2.44E-12 |
| pL4440 | vs | *isp-1 (1:2)* | 1.18E-03 |
| **Test Strains** | | | |
| **3. SLR152** | | | |
| pL4440 | vs | *isp-1 (1:2)* | 4.00E-15 |
| **4. SLR152 Sans Array** | | | |
| pL4440 | vs | *isp-1 (1:2)* | 2.48E-14 |
| **5. SLR155** | | | |
| pL4440 | vs | *atp-3 (1:2)* | 2.26E-02 |
| pL4440 | vs | *isp-1 (1:2)* | 8.83E-14 |
| **6. SLR155 Sans Array** | | | |
| pL4440 | vs | *atp-3 (1:2)* | 8.81E-18 |
| pL4440 | vs | *isp-1 (1:2)* | 2.26E-02 |
| **7. SLR153.2** | | | |
| pL4440 | vs | *atp-3 (1:2)* | 6.38E-04 |
| pL4440 | vs | *isp-1 (1:2)* | 6.42E-11 |
| **8. SLR153.2 Sans Array** | | | |
| pL4440 | vs | *atp-3 (1:2)* | 4.35E-16 |
| pL4440 | vs | *isp-1 (1:2)* | 6.47E-01 |

* Red = significant after correction for multiple testing (Bonferroni Correction) p < 0.05/30 or 0.0017)

**Log Rank Analyses Contd.**

| Between Strains Comparisons | |
|---|---|
| ***Strains & Conditions*** | ***Long Rank Test (p-value*)*** |
| ***1. SLR115 vs. pmk-3(tm745)*** | |
| pL4440 | 3.34E-04 |
| *atp-3 (1:2)* | 3.65E-13 |
| *isp-1 (1:2)* | 7.86E-06 |
| ***2. SLR152 w/w.o. Array*** | |
| pL4440 | 8.30E-01 |
| *isp-1 (1:2)* | 4.90E-01 |
| ***3. SLR155 w/w.o. Array*** | |
| pL4440 | 8.08E-03 |
| *atp-3 (1:2)* | 3.38E-09 |
| *isp-1 (1:2)* | 9.10E-07 |
| ***4. SLR153.2 w/w.o. Array*** | |
| pL4440 | 2.37E-01 |
| *atp-3 (1:2)* | 3.33E-18 |
| *isp-1 (1:2)* | 4.97E-08 |
| ***5. SLR152 (+) Array vs. SLR115*** | |
| pL4440 | 8.08E-02 |
| *isp-1 (1:2)* | 2.48E-01 |
| ***6. SLR155 (+) Array vs. SLR153.2 (+) Array*** | |
| pL4440 | 2.35E-02 |
| *atp-3 (1:2)* | 5.35E-01 |
| *isp-1 (1:2)* | 3.76E-02 |

* Red = significant after correction for multiple testing (Bonferroni Correction) p < 0.05/30 or 0.0017)

### Study #3 (Corresponds to Fig. 5E)

**Control Strains**

**1.SLR115 [Ptbb-6::GFP]**
**2. *pmk-3(tm745)***
**3. SLR0161 [*Pvha-6::pmk-3-L::SL2::mCherry*]**
**4. SLR0161 Sans Array**
**5. SLR153.2* [*Pvha-6::pmk-3-L::SL2::mCherry; Ptbb-6::GFP; pmk-3(tm745)*]**
**6. SLR153.2 Sans Array [*Ptbb-6::GFP; pmk-3(tm745)*]**
*** SLR0153 Line 2**

**Test Strains**

**7. SLR164 Line 1 [*Pvha-6::dlk-1 b(EE)::SL2::mCherry); Pvha-6::pmk-3-L::SL2::mCherry; Ptbb-6::GFP; pmk-3(tm745)]*]**
**8. SLR164 Line 1 Sans Arrays [*Ptbb-6::GFP; pmk-3(tm745)]]***

**9. SLR164 Line 2 [*Pvha-6::dlk-1 b(EE)::SL2::mCherry); Pvha-6::pmk-3-L::SL2::mCherry; Ptbb-6::GFP; pmk-3(tm745)]*]**
**10. SLR164 Line 2 Sans Arrays [*Ptbb-6::GFP; pmk-3(tm745)]]***

**Descriptive Statistics**

| Strain | Condition | Sample Size (censored events) | Expected Life Span (days) |
|---|---|---|---|
| **Control Strains** | | | |
| **1. SLR115** | OP50 | 65 (18) | 17.5 |
| **2. *pmk-3(tm745)*** | OP50 | 70 (7) | 16.3 |
| **3. SLR161** | OP50 | 70 (12) | 13.0 |
| **4. SLR161 Sans Arra** | OP50 | 69 (12) | 13.2 |
| **5. SLR153.2** | OP50 | 70 (11) | 17.1 |
| **6. SLR153.2 Sans Ar** | OP50 | 69 (10) | 16.9 |
| **Test Strains** | | | |
| **7. SLR164 Line 1** | OP50 | 55 (16) | 15.4 |
| **8. SLR164 Line 1 San** | OP50 | 64 (9) | 15.5 |
| **9. SLR164 Line 2** | OP50 | 61 (14) | 13.5 |
| **10. SLR164 Line 2 Sa** | OP50 | 63 (16) | 15.1 |

**Log Rank Analyses**

**Between Strains Comparisons**

| *Strains* | *Long Rank Test (p-value*)* |
|---|---|
| ***1. SLR115 vs. pmk-3(tm745)*** | 2.06E-01 |
| ***2. SLR115 vs. SLR153.2 Sans Array*** | 5.21E-01 |
| ***3. SLR115 vs SLR0161 Sans Array*** | 3.85E-04 |
| ***4. SLR115 vs SLR164 Line 1 Sans Arrays*** | 7.61E-02 |
| ***5. SLR115 vs SLR164 Line 1 Sans Arrays*** | 5.57E-02 |
| ***6. SLR153.2 w/w.o. Array*** | 7.06E-01 |
| ***7. SLR0161 w/w.o. Array*** | 5.61E-01 |
| ***8. SLR164 Line 1 w/w.o. Array*** | 8.32E-01 |
| ***9. SLR164 Line 2 w/w.o. Array*** | 1.14E-01 |

* Red = significant after correction for multiple testing (Bonferroni Correction) p < 0.05/9 or 0.0056)

### Study #4 (Corresponds to Fig. S7A)

**Strains**

**1. SLR153.1*** ***[Pvha-6::pmk-3-L::SL2::mCherry; Ptbb-6::GFP; pmk-3(tm745)]***

**2. SLR153.1 Sans Array** ***[Ptbb-6::GFP; pmk-3(tm745)]***

*** SLR0153 Line 1**

**Descriptive Statistics**

| Strain | Condition | Sample Size (censored events) | Expected Life Span (days) |
|---|---|---|---|
| **Test Strains** | | | |
| **1. SLR153.1** | pL4440 *(empty vector)* | 174 (18) | 19.6 |
| | *isp-1 (1:10)* | 158 (15) | 24.8 |
| **2. SLR153.1 Sans Array** | pL4440 *(empty vector)* | 170 (22) | 17.6 |
| | *isp-1 (1:10)* | 168 (30) | 19.3 |

**Log Rank Analyses**

| Within Strain Comparisons | | | Long Rank Test (p-value*) |
|---|---|---|---|
| **1. SLR153.1** | | | |
| pL4440 | vs | *isp-1 (1:10)* | 8.69E-09 |
| **2. SLR153.1 Sans Array** | | | |
| pL4440 | vs | *isp-1 (1:10)* | 1.92E-04 |

* Red = significant after correction for multiple testing (Bonferroni Correction) p < 0.05/6 or 0.0083)

**Log Rank Analyses Contd.**

| Between Strains Comparisons | |
|---|---|
| ***Strains & Conditions*** | ***Long Rank Test (p-value*)*** |
| ***1. SLR153.1 w/w.o. Array*** | |
| pL4440 | 7.20E-04 |
| *isp-1 (1:10)* | 2.15E-08 |
| ***2. SLR155** (+) Array vs. SLR153.1 (+) Array*** | |
| pL4440 | 4.20E-02 |
| *isp-1 (1:10)* | 3.06E-01 |

* Red = significant after correction for multiple testing (p < 0.0083)

** SLR155 data from Study #1 above

### Study #5 (Corresponds to Fig. 7A)

**Strain**

1.SLR115 [Ptbb-6::GFP]

**Descriptive Statistics**

| Strain | Condition (RNAi #1 | + RNAi #2) | Sample Size (censored events) | Expected Life Span (days) |
|---|---|---|---|---|
| SLR115 | *pL4440 (9:10)* | *pL4440 (1:10)* | 180 (16) | 15.9 |
| | *pL4440 (9:10)* | *isp-1 (1:10)* | 120 (30) | 31.4 |
| | *alx-1 (9:10)* | *pL4440 (1:10)* | 180 (17) | 11.7 |
| | *alx-1 (9:10)* | *isp-1 (1:10)* | 120 (35) | 31.2 |
| | *C01A2.4 (9:10)* | *pL4440 (1:10)* | 123 (14) | 16.4 |
| | *C01A2.4 (9:10)* | *isp-1 (1:10)* | 120 (31) | 28.5 |
| | *istr-1 (9:10)* | *pL4440 (1:10)* | 120 (16) | 14.1 |
| | *istr-1 (9:10)* | *isp-1 (1:10)* | 123 (17) | 14.6 |
| | *rab-11.1 (9:10)* | *pL4440 (1:10)* | 95 (21) | 6.4 |
| | *rab-11.1 (9:10)* | *isp-1 (1:10)* | 121 (30) | 7.3 |

**Log Rank Analyses**

**Within RNAi GROUP Comparisons**

| Condition 1 (RNAi #1 | + RNAi #2) | versus | Condition 2 (RNAi #1 | + RNAi #2) | Long Rank Test (p-value*) |
|---|---|---|---|---|---|
| *pL4440 (9:10)* | *pL4440 (1:10)* | *vs* | *pL4440 (9:10)* | *isp-1 (1:10)* | 0.00E+00 |
| | | *vs* | *alx-1 (9:10)* | *pL4440 (1:10)* | 0.00E+00 |
| | | *vs* | *alx-1 (9:10)* | *isp-1 (1:10)* | 0.00E+00 |
| | | *vs* | *C01A2.4 (9:10)* | *pL4440 (1:10)* | 5.01E-01 |
| | | *vs* | *C01A2.4 (9:10)* | *isp-1 (1:10)* | 0.00E+00 |
| | | *vs* | *istr-1 (9:10)* | *pL4440 (1:10)* | 3.30E-09 |
| | | *vs* | *istr-1 (9:10)* | *isp-1 (1:10)* | 1.00E-04 |
| | | *vs* | *rab-11.1 (9:10)* | *pL4440 (1:10)* | 0.00E+00 |
| | | *vs* | *rab-11.1 (9:10)* | *isp-1 (1:10)* | 0.00E+00 |
| *pL4440 (9:10)* | *isp-1 (1:10)* | *vs* | *alx-1 (9:10)* | *isp-1 (1:10)* | 3.01E-01 |
| | | *vs* | *C01A2.4 (9:10)* | *isp-1 (1:10)* | 1.28E-02 |
| | | *vs* | *istr-1 (9:10)* | *isp-1 (1:10)* | 0.00E+00 |
| | | *vs* | *rab-11.1 (9:10)* | *isp-1 (1:10)* | 0.00E+00 |

* Red = significant after correction for multiple testing (Bonferroni Correction) p < 0.05/13 or 0.0038)
