## Supplemental Text S1 for "The PMK-3 (p38) Mitochondrial Retrograde Response Functions in Intestinal Cells to Extend Life via the ESCRT Machinery"

### SUPPLEMENTAL MATERIAL

**File Summary:**

**S1 Text.** Supplemental experimental procedures.

**Figure S1.** Selection of a PMK-3 epitope for antibody generation.

**Figure S2.** Sensitivity of PMK-3 antisera.

**Figure S3.** Specificity of PMK-3 antisera.

**Figure S4.** Raw data for western blot images shown in **Fig. 1B**.

**Figure S5.** Structure-based sequence alignment of PMK-3(L) with other p38 MAPK isoforms.

**Figure S6.** Additional fluorescent images of *pmk-3(tm745)* knockout mutants expressing *pmk-3* transgenes under the control of the *rgef-1* neuronal promoter.

**Figure S7.** Additional survival analyses.

**Figure S8**. *myc*-tagged DLK-1b(EE) expressed in *dlk-1(tm4024)* knockout mutants becomes activated only in the presence of ETC stress.

**Figure S9.** No alteration in PMK-3::GFP fluorescence following mitochondrial ETC disruption in DA1750 worms.

**Figure S10.** Knockdown of *istr-1* by RNAi does not inhibit the efficacy of the RNAi machinery.

**Figure S11.** Extracorporeal EVs isolated from starved *isp-1(qm150)* mutants do not activate ETC stress reporter genes.

**Table S1.** Worm strains.

**Table S2.** Lifespan Analyses

### SUPPLEMENTAL EXPERIMENTAL PROCEDURES

**Additional Strain Information.** Strain GJ3080 [gjEx1863(*pmk-3*::GFP; P*elt-2*::mCherry] was generously provided by Dr. Gert Jansen (Erasmus MC, Netherlands). Construction of this line has been described before [1], but briefly it was generated by injecting N2 Bristol worms with a fosmid rescued from TransgeneOme clone 2666739528585155 H05 (at a concentration of 10 ng/µl). This fosmid contains a genomic fragment of *C. elegans* containing full length versions of the following genes (in order): F42G8.10, *moc-3*, *sph-1, isp-1, islo-1, pmk-3, pmk-2, pmk-1, faah-2,* and *B0218.5.* Of note, the entire *islo-1, pmk-3, pmk-2* and *pmk-1* operon is present. The *pmk-3* gene in this fosmid is C-terminally tagged with the following multi-antigen tag (in order, 2xTY1 epitope, EGFP, FRT site, 3xFlag epitope (where x means number of copies of the epitope in tandem). The fosmid also contains the *C. briggsae unc-119* gene and a *NAT* selection cassette elsewhere on the construct. Expression from this transgenic array is designed to mimic endogenous gene levels (a so-called "third allele") [2].

Strains SLR0246, SLR0247 and SLR0248 were constructed by injecting HT1593 [*unc-119(ed3)*] mutant worms with rescued fosmid DNA from TransgeneOme bacterial clones

095794875301489 F02 (pSR10), 7225558184899882 D12 (pSR11) and 5438954842362824 D10 (pSR12), respectively, at a concentration of 70-95 ng/µl. TransgeneOme clones were purchased from Source BioScience (see **Table S1** for additional details).

**Transgene Construction and Transgenic Strain Generation.** In regard to the following cloning experiments, all restriction enzymes were obtained from New England Biolabs (MA), all PCRs were undertaken using Phusion (Thermo Fisher), and for all ligations we utilized T4 DNA ligase (New England Biolabs, MA).

***Pvha-6::DLK-1b(EE)::SL2::mCherry*:** An NheI – KpnI fragment containing Dlk-1b(EE) was removed from plasmid pCZGY1969 (a generous gift from Dr. Yishi Jin, UC San Diego), and cloned into the same sites of plasmid P*vha-6*::*sid-1*(+)::SL2::GFP (a generous gift from Alex Soukas, Harvard University) to generate P*vha-6*::DLK-1b(EE)::SL2::GFP. The GFP reporter was then exchanged with a PCR generated mCherry fragment using the AgeI and EcoRI sites to generate plasmid pSR1. The plasmid *Ptbb-6::mCherry::attB2::unc-54 3'UTR/*pBluescript-SK(-) [3] was used as template for the PCR reaction. The entire transgene was sequence confirmed and then microinjected into strain CL3462 (*smg-1(cc546) I; dvIs67[pCL179(Ptbb-6::GFP) + Pmyo-3::dsRed]*) at a concentration of 20 ng/µl.

***Pvha-6::DLK-1b(EE)::SL2::mCherry*:** Site-directed mutagenesis was used to introduce a Myc-tag at the C-terminus of DLK-1(EE) in vector pSR1. Briefly, the following primer pair was employed with the Q5® Site-Directed Mutagenesis Kit from NEB (cat # E0554S).

*FW primer*: 5'-AGCGAAGAAGATCTGTAAGGTACCGCTGTCTCATC-3'

*RV primer*: 5'-AATCAGTTTCTGTTCAATTCGGACTGCTCCGGC-3'

The resulting alteration to plasmid pSR1 was confirmed by sequencing, generating the new plasmid P*vha-6*::DLK-1b(EE)myc::SL2::mCherry (hereafter called plasmid pSR16).

***Pvha-6::PMK-3(L)::SL2::mCherry:*** A cDNA clone encoding the long isoform of *pmk-3* [PMK-3(L)] was retrieved from N2 Bristol (wild type worms) using RT-PCR and the following primer pair:

*pmk-3* fw 1: 5'-TTTGCTAGCATGGCGTCAGTCCCATC -3'

*pmk-3* rev 2: 5'-TTGGTACCTCAGCGATCTGCTTCTCCAG-3'

These primers introduced a NheI and KpnI restriction site 5' and 3' to the *pmk-3* sequence, respectively. The resulting PCR fragment was blunt-end cloned into the SmaI site of pBluescript SK(-) and sequenced. *pmk-3* cDNA contains an endogenous KpnI site and it was removed using the Phusion Site-Directed Mutagenesis Kit (Thermo Scientific) in conjunction with the following primer pair:

*pmk-3 KpnI SDM fw*: 5'-CAAACAAGATGGTATCGTAGTCCAGAAGTC-3'

*pmk-3 KpnI SDM rev*: 5'-TACATATTGTGTTAGAGAAGTATCTTTCTTTTCC-3'

This primer pair preserved the underlying amino acid sequence encoded by the *pmk-3* cDNA. The resulting alteration to *pmk-3* was confirmed by DNA sequencing. The final construct was designated*)* pBluescript SK(-)/*pmk-3(L)(sans_Kpn1).* Finally, the NheI - KpnI fragment containing *pmk-3(L)(sans Kpn1)* was swapped with the NheI - KpnI fragment of P*vha-6*::DLK-1b(EE)::SL2::mCherry to generate P*vha-6*::PMK-3(L)::SL2::mCherry (hereafter called plasmid pSR2).

***Pvha-6::PMK-3(S)::SL2::mCherry****:* The short isoform of *pmk-3* [PMK-3(S)] was derived from pBluescript SK(-)/*pmk-3(L)(sans_Kpn1)*, using PCR and the following primer pair:

*pmk-3 short form fw*: 5'-CATGCTAGCATGAGACATGACAATATCATC-3'

*pmk-3 short form rev*: 5'-TTGGTACCTCAGCGATCTGCTTC-3'

These primers introduced NheI and KpnI sites, 5' and 3', respectively to the PMK-3(S) sequence. Following restriction with NheI and KpnI, the PCR fragment encoding *pmk-3(S)* was swapped with the NheI - KpnI fragment of P*vha-6*::DLK-1b(EE)::SL2::mCherry to generate P*vha-6*::PMK-3(S)::SL2::mCherry (hereafter called plasmid pSR3).

***Peft-3::PMK-3(L)::SL2::mCherry*** and ***Peft-3::PMK-3(S)::SL2::mCherry***: The *eft-3* promoter was PCR-amplified from plasmid P*eft-3*::2mt8::ArchT::EGFP::*tbb-2* 3' UTR using the following primer pair:

*Peft-3 NotI fw*: 5'-CATGCGGCCGCGCACCTTTGGTCTTTTATTGTCAACTTCC-3'

*Peft-3 AscI rev*: 5'-CATGGCGCGCCTGAGCAAAGTGTTTCCCAACTGAAA-3'

These primers introduced NotI and AscI sites, 5' and 3', respectively to the *eft-3* promoter sequence. These sites were then used to exchange the *vha-6* promoter from both pSR2 (P*vha-6*::PMK-3(L)::SL2::mCherry) and pSR3 (P*vha-6*::PMK-3(S)::SL2::mCherry) with the *eft-3* promoter (hereafter called plasmids pSR4 and pSR5, respectively).

***Peft-3::PMK-3(L)_EQE::SL2::mCherry, Peft-3::PMK-3(S)_EQE::SL2::mCherry, Peft-3::PMK-3(L)_AQA::SL2::mCherry*** *and* ***Peft-3::PMK-3(S)_AQA::SL2::mCherry:*** To introduce the EQE and AQA double point mutations into the PMK-3 sequence of both pSR4 (P*eft-3*::PMK-3(L)::SL2::mCherry) and pSR5 (P*eft-3*::PMK-3(S)::SL2::mCherry) plasmid sequences, site-directed mutagenesis using the Phusion Site-Directed Mutagenesis Kit (Thermo Scientific) in conjunction with following two primer pairs was employed:

*For the EQE mutation:*

CA pmk-3 SDM fw: 5'-AAAGATACTTCTCTAGAACAAGAAGTACAAACAAGATGG-3'

CA pmk-3 SDM rev: 5'-CTTTTCCAAGGATCTTGCAAGTCCAAAATCGAG-3'

*For the AQA mutation:*

CI pmk-3 SDM fw: 5'-AAAGATACTTCTCTAGCACAAGCTGTACAAACAAGATGG-3'

CI pmk-3 SDM rev: 5'-CTTTTCCAAGGATCTTGCAAGTCCAAAATCGAG-3'

The resulting plasmids, *Peft-3::PMK-3(L)_EQE::SL2::mCherry, Peft-3::PMK-3(S)_EQE::SL2::mCherry, Peft-3::PMK-3(L)_AQA::SL2::mCherry and Peft-3::PMK-3(S)_AQA::SL2::mCherry,* were called pSR6, pSR7, pSR8 and pSR9, respectively.

***Prgef-1::PMK-3(L)::SL2::mCherry:*** PMK-3(L) was placed under control of the *rgef-1* promoter as follows. A NotI/NheI vector fragment of pSR2 (*Pvha-6::PMK-3(L)::SL2::mCherry)* was isolated. Next the primer pair:

Prgef-1 fw (NotI site): 5'-GCATGCGCGGCCGCGCATGCAAGACTAATTTTCGATTAACCCG-3'

Prgef-1 rev (NheI site): 5'-GCCATGCTAGCTCAACTTTACTGCTGATCGTCGTCGTCGTCGATGCCG-3'

was used to PCR amplify the *rgef-1* promoter sequence from vector pDEST- F25B3.3p (a generous gift from Dr. Hidehito Kuroyanagi, Tokyo Medical and Dental University, Japan). This product was then cut with NotI and NheI, and sub-cloned into the corresponding sites of the pSR2 vector fragment forming *Prgef-1::PMK-3(L)::SL2::mCherry* (hereafter called pSR13). pSR13 was sequence verified across the full *Prgef-1* promoter region.

***Prgef-1::(decoyATG)::PMK-3(L)::SL2::mCherry:*** We inserted a small 'decoy' open reading frame (ORF) in front of the normal ATG of *rgef-1* in an effort to reduce bleed-through expression in non-neuronal tissues from this promoter. We used the decoy ORF contained in the *vha-6* promoter of P*vha-6*::*sid-1*(+)::SL2::GFP (which incidentally was included in all our P*vha-6* containing constructs). Briefly, the following primer pair was used to amplify the *rgef-1* promoter from pDEST-F25B3.3p :

Prgef-1 FW (NotI site): 5'-GCATGCGCGGCCGCGCATGCAAGACTAATTTCGATTAACCCG-3'

Prgef-1 RV (NheI/TspMI site): 5'-ATATCCCGGGATGCTAGCTCAACTTTACTGCTGATCGTCGTC GTCGTCGATGCCG-3'

This amplicon was then cut with NotI and TspMI and exchanged into the corresponding sites of pSR2 to form vector *Prgef-1::PMK-3(L)::SL2::mCherry* (hereafter called pSR14). pSR14 was sequence verified across the entire *Prgef-1* +decoy ORF promoter region.

***Prab-3::(decoyATG)::PMK-3(L)::SL2::mCherry:*** The *rab*-3 promoter was PCR amplified from N2 genomic DNA using the following primer pair:

P*rab3* FW NotI : 5'-GCATGCGCGGCCGCATCTTCAGATGGGAGCAGTGGAC-3'

P*rab3* RV NheI/TspMI : 5'-
ATATCCCGGGATGCTAGCCTGAAAATAGGGCTACTGTAGATTTATTTTAAAAGAGCATTTAG-3'

The resulting amplicon was cut with NotI and TspMI, then exchanged into the corresponding sites of pSR2, to form *Prab-3::(decoyATG)::PMK-3(L)::SL2::mCherry* (hereafter called pSR15). As for pSR14, this construct contained a small decoy ORF immediately after the *rab-3* promoter and before the PMK-3(L) ORF in order to blunt leak-through expression in non-neuronal tissues.

Following plasmid construction and sequence verification, the eleven newly generated PMK-3 transgenic constructs were individually microinjected into strain SLR150 (*pmk-3(tm745) IV; dvIs67[pCL179(Ptbb-6::GFP) + Pmyo-3::dsRed]*) at a concentration of 20 ng/µL. Names of strains containing relevant extrachromosomal arrays are provided in **Table S1**.

**Feeding RNAi.** All feeding RNAi experiments were undertaken using HT115 *Escherichia coli* bacteria. Bacterial feeding RNAi constructs targeting *atp-3* and *isp-1* have been described previously [4]. C01A2.4 (CHMP2B) and *skn-1* RNAi feeding constructs were obtained from the Ahringer Feeding RNAi Library [5]. *istr-1* (K10C8.3) and *rab-11.1* (F53G12.1) RNAi feeding constructs were purchased from Dharmacon (catalogue #: RCE1182-202297727 and RCE1182-202296485, respectively). An *alx-1* (R10E12.1) feeding RNAi construct was generated by PCR using the following primer pair:

*alx-1* fw: 5'- GCCGCGCTCGAGCAGAATGGCTACATTCGGCTTTCTCAGCGC -3'

*alx-1* rev: 5'- CCGCCCAGATCTTATTTTCGGGCCACTTTGAGCCAACG -3'

pSR10 was used as genomic DNA template. The resulting amplicon was digested with BglII and XhoI then sub-cloned into the BamHI and SalI sites of pL4440. We have previously developed and published standardized procedures for *C. elegans* feeding RNAi and for the construction of bacterial feeding RNAi dilution series [4]. These procedures were utilized unchanged in this study.

To test whether reduction of *istr-1* altered the efficacy of the RNAi machinery, we used a dual feeding RNAi approach and asked whether the potency of *skn-1* RNAi to sterilize adult worms was differentially reduced in worms co-fed RNAi targeting *istr-1* versus empty vector (**Fig. S10**). Briefly, SLR0115 worms were fed bacteria containing *skn-1* and increasing amounts of each *test* RNAi, from the arrested L1 stage onward. At the L4/YA boundary, exactly 20 animals were transferred to fresh RNAi plates and then allowed to lay eggs for another 24 or 36 hours (depending on the experimental replicate, n=2 independent replicates). The number of dead eggs and hatchlings on each plate was then counted (~1000 progeny per plate). These values were converted into percent survival across the full dilution series for each *test* RNAi, then significance testing for differences in the shape of the resulting curve was undertaken using a log-rank test. No significant effect of *istr-1* was observed.

**Western Analyses.** Worm extracts were prepared as previously described using either 1% SDS (whole-worm extract) or 1% Triton-X-100 (Tx-100) detergent [6]. The latter samples were collected using a Balch homogenizer and fractionated into soluble and insoluble components, as previously described [6]. For western blotting we used the following reagents and suppliers: 4-12% Bis-Tris NuPAGE Gels (Invitrogen); nitrocellulose (Protran, BA83); 5% milk powder in Tris-buffered saline + 0.05% Tween-20 (TBS-$T_{0.05\%}$) for blocking; TBS-$T_{0.05\%}$ for washing steps, α-PMK-3 primary antibody (Rabbit #75, 70-day bleed) incubated at 4$^{o}$C for 24-48 hours, HRP-coupled goat α-rabbit secondary antibody (ab6721, Abcam) incubated at 23$^{o}$C for one hour. α-PMK-3 reactive bands were detected using chemiluminescence (Pierce) and x-ray film (CL-Xposure, ThermoFisher Scientific).

**PMK-3 Antisera Generation.** α-PMK-3 antisera was generated under contract by Life Technologies Corporation (ThermoFisher Scientific). Briefly, the following peptide: DDSQARTRFEWRDAVWKE, corresponding to αα 438-452 of PMK-3(L), was coupled to Keyhole Limpet Hemocyanin (KLH) and injected into two rabbits (#74 and #75). A pre-bleed was taken, test bleeds were taken at days 28, 56 and both animals were exsanguinated at day 70/72. Sera was aliquoted and stored at -80$^{o}$C without further processing.

**Sensitivity-testing of PMK-3 Antisera.** The sensitivity of newly-generated PMK-3 antisera (Rabbit #75, 70-day bleed) was tested as follows: As described above, GJ3080 worms contain an unstable extrachromosomal array designed to express its constituent genes at, or near, endogenous levels [2]. The array of GJ3080 worms includes the entire *islo-1, pmk-3, pmk-2* and *pmk-1* operon. A GFP tag is inserted C-terminally on PMK-3. Fluorescence analysis of live worms reveals PMK-3::GFP is ubiquitous and nuclear localized in GJ3080 worms, but present at very low abundance since long exposures were required to image these animals (**Fig. S2B)**. This finding is in agreement with an earlier report [1]. Western analysis of array-containing GJ3080 worms using our PMK-3 antisera also reported low-level PMK-3::GFP expression, with blots requiring long exposures, and signal present just above background. Notably, despite being present in low abundance, the (L) form of the recombinant GFP fusion protein was unambiguously detectable (**Fig. S2C,** compare worms with and without the array). Presence of the recombinant (S) form was less certain. Endogenous PMK-3 isoforms could not be reliably detected under these conditions. We confirmed that the PMK-3::GFP made by the array in GJ3080 worms is functional because *pmk-3(tm745)*; P*tbb-6*::GFP mutants containing the array (strain SLR0166) recovered their ability to initiate retrograde response signaling when exposed to *atp-3* feeding RNAi, as indicated by strong induction of the gut-specific P*tbb-6*::GFP reporter (**Fig. S2D**). Notably, GJ3080 worms exposed to *atp-3* RNAi showed no change in the subcellular distribution or intensity of PMK-3::GFP in any tissue, consistent with PMK-3 activity likely being regulated post-translationally.

**Specificity-testing of PMK-3 Antisera.** To test whether our newly-generated PMK-3 antisera cross-reacted with PMK-1 or PMK-2 we undertook the following experiments using sera from Rabbit #75 (70-day bleed). PMK-1 is expressed ubiquitously, while PMK-2 is restricted to neurons [7]. Both proteins are closely-related to PMK-3 at the primary sequence level (**Fig. S1**). We confirmed PMK-3 antisera specificity as follows: Under basal conditions, wild type (N2) worms incubated with PMK-3 antisera showed one weakly cross-reactive band upon western analysis that did not match the expected size of any PMK-1 or PMK-2 isoform (**Fig. S3A**). A band of the same size was also present in *pmk-3(tm745)* knock-out mutants. This protein was found to be of bacterial origin and reflected *E. coli* caught either in the gut or attached to the cuticle during sample extract preparation (**Fig. S3B**). ZD721 worms contains a gain-of function mutation in PMK-2 that results in its constitutive activation [7]. These worms also showed no cross-reactivity with PMK-3 antisera by western analysis (**Fig. S3A**). AU78 worms contain a GFP reporter gene that is strongly induced following infection by the pathogenic PA14 strain of *Pseudomonas aeruginosa,* in a PMK-1 dependent manner [8]. Unexpectedly, three PMK-3 cross-reactive bands were apparent in AU78 worms following acute exposure to PA14 (**Fig. S3A**). Further studies revealed all three proteins were of PA14 origin, and not related to PMK-3 (**Fig. S3B**). Consistent with this conclusion, we observed that a GFP reporter gene coupled to the proximal promoter of *pmk-3* (**Fig. 1A**) was dampened, not elevated, by growth on PA14 (**Fig S2E**).

**Homology Modeling.** An homology model of PMK-3(L) was built using the algorithm of Nielsen and colleagues [9] via the CPHmodels-3.3(beta) server (http://www.cbs.dtu.dk/services/CPHmodels/). Human p38α (C162S) [Protein Data Bank (PDB) structure 1R3C [10]] served as the model-building template (PDB Blast E-value= 2e-75). Amino acids 110-459 of PMK-3(L) were included in the final model. PMK-1 and PMK-2 were modeled similarly, using PDB structures 4LOQ (*Mus musculus* p38α) and 5ETC (inactive human p38α with an ordered activation loop), as structural templates.

**Lifespan Analyses**. Lifespan studies were performed as described previously [11], and without the use of FudR. All plates were maintained at 20ºC. The first day of adulthood was designated as day one. A full description of all lifespan experiments is provided in **Table S2.** Survival data was analyzed for significance using the Log-rank test. Unadjusted significance values ($p<0.05$) are provided in tables, but they are color coded to highlight comparisons that remained significant after correction for multiple testing (Bonferroni method). Refer specifically to the footnote in relevant tables for precise Bonferroni adjustments that were applied.

***In silico*** **Data Mining.** The *C. elegans* v4 cisRED database [12] contains conserved DNA motifs in the upstream promoter regions of 3847 *C. elegans* transcripts (WS170 genome release). Only transcripts with orthologs in three or more of the following nematode species were included in the database - *C. briggsae, C. brenneri, C. remanei, C. japonica, Pristionchus pacificus, Brugia malayi* or *Trichinella spiralis*. Promoters were defined as extending 1500 kb upstream of the transcription start site, or to the nearest upstream gene, which ever was smallest. Motif 2, identified previously in the promoter of PMK-3 responsive genes (see Fig. 1C in [3]) was used to query the cisRED database. This sequence was approximated using RTTKYGAAAY, where R = G or A, K = G or T, Y = T or C and both nucleotide choices were weighted equally. This analysis identified 206 elements spread over 188 genes. We used the DAVID Bioinformatics toolbox (v6.8) [13] to identify cellular pathways in which these genes functioned. We found a statistically-significant over-representation of genes involved in endocytosis (9/188 genes, *p-value*: 0.000072), which remained significant after correction for multiple comparisons (Benjamini-Hochberg FDR method, *q-value*: 0.0025). Proteins encoded by these genes were plotted onto the Endocytosis map of KEGG [14, 15].

**Microarray Analysis of Endocytosis Gene Set.** We used the GEO profiles database [16, 17] to analyze microarray data (GEO#: GSE38196) [18] for changes in mRNA expression of nine endocytosis genes of interest (*alx-1, C01A2.4, istr-1, rab-11.1, rfip-1, rme-1, usp-50, vps-2, vps-32.1*). mRNA for these studies was isolated from wild type worms and *atfs-1(tm4525)* loss-of-function mutants that had been synchronized by bleach treatment, raised in liquid culture in the presence of *spg-7* or vector (pL4440)-only feeding RNAi (HT115 bacteria), then harvested at the L4 stage. Three independent samples were collected for each of the four conditions. mRNA was quantified using the GeneChip *C. elegans* genome array (Affymetrix). Two probe sets were available for *alx-1, istr-1, rfip-1, rme-1, usp-50* and *vps-2.* Three probe sets were available for *rab-11.1*. We normalized mRNA expression level across each probe set using wild type worms cultured on vector as the normalization control. We then tested for differences in mRNA expression across conditions using Student's t-test ($p<0.05$), again setting wild type worms cultured on vector as the normalization control. Error propagation methods were used to calculate a variance for each test condition. We controlled for multiple comparisons using the Benjamini-Hochberg FDR method ($q<0.05$). We also pooled probe sets for each respective gene and repeated the analysis. Our conclusions did not change from those reported in **Figure 6D**, except *usp-50* required the FDR threshold to be raised to $q<0.08$ to reach significance.

**Quantitative PCR Analysis of Endocytosis Gene Set.** We used reverse transcription quantitative PCR (RT-qPCR) to measure changes in mRNA expression of *alx-1, C01A2.4, istr-1, rab-11.1, rfip-1, rme-1, usp-50, vps-2* and *vps-32.1* in worms exposed to *isp-1* mediated ETC disruption, with and without functional PMK-3 present. Three to seven fully independent biological replicates were collected and analyzed per test condition. Briefly, SLR115 (wild type) and SLR0150 [*pmk-3(tm745)*] worms were cultured on lawns containing pL4440 vector only (control), or feeding RNAi lawns targeting *isp-1* at full strength or diluted to one-tenth strength (balanced with pL4440 control bacteria). Animals were seeded as synchronously arrested L1, obtained from freshly starved NGM/OP50 plates (< 24 hours starvation), at a density of 1000 worms per 10 cm diameter plate. When animals became one day old adults, 2000 worms per condition were collected using S-Basal [19], pooled, washed free of bacteria (3 x 15 ml S-Basal), then frozen at -80°C. RNA was isolated from each sample using TRIzol Reagent (Thermo Fisher Scientific) in conjunction with column purification (RNeasy Mini Kit, Qiagen, #74104). Contaminating genomic DNA was removed on column using RNase-free DNase (Qiagen, #79254). RNA was eluted into water. cDNA was prepared using MultiScribe reverse transcriptase (Applied Biosystems, #4308228), oligo $(dT)_{18}$ (Thermo Fisher Scientific), RiboLock RNase Inhibitor (Thermo Fisher Scientific) and 0.5 -1 µg template RNA. For qPCR, we used the PowerUp SYBR Green Master Mix (Applied Biosystems, #A25742) in conjunction with a StepOnePlus Real Time PCR System (Thermo Fisher Scientific, # 4376600). All primers were used at a final concentration of 300 nM (refer to the following table for specific primer pairs)

| **Target gene** | **Primer (5'-3')** |
|---|---|
| *alx-1,* forward | TGGCTACATTCGGCTTTCTC |
| *alx-1,* reverse | GCGACATCTGAGCGGTTATTA |
| *CO1A2.4,* forward | CGAACAAGACGCTATCGTGAA |
| *CO1A2.4,* reverse | CTGGAAACCTTTGTGGGAAGA |
| *istr-1,* forward | CCACCACAGTCACTTCGTAAA |
| *Istr-1,* reverse | CTGGACCTCCTGGCTTATTG |
| *rab-11.1,* forward | GTCGAATCTCCTGTCTCGTTTC |
| *rab-11.1,* reverse | CCTTCACTGTCTTGCCTTCTAC |
| *rfip-1,* forward | AAAGCCAAAGTAGCAGAACTAATG |
| *rfip-1,* reverse | TCAGCAAGTGACGGTGTATC |
| *rme-1,* forward | TTGGCTGGAAACGAGGATAAA |
| *rme-1,* reverse | AGCTCCGTAGACTCTCATCAA |
| *ups-50,* forward | ACACTTCCAGAACGACCAATAG |
| *ups-50,* reverse | ATTCGGCTTAGTTGCTCTATCC |
| *vps-2,* forward | GCAGCTGAATCTTCCACAAATC |
| *vps-2,* reverse | CCATCACCTCCTCCTTCATATC |
| *vps-32.1,* forward | TGGCTCTCCAGTGTTTGAATAG |
| *vps-32.1,* reverse | TTCGAGGGCTTCTCTCTGATA |
| *pmp-3,* forward | GATGAGCAGGAAGAGGAAGAAG |
| *pmp-3,* reverse | GGTGAAAGTGTAGCCGAATCTA |
| *pmk-3,* forward | CGGGCTTCGTCGTTATGTT |
| *pmk-3,* reverse | AGACTACGACCCGCAAATTC |
| *tbb-6*, forward | TCTTCAAACTGACGGGACATAC |
| *tbb-6*, reverse | AATCGCACGAGGAACATACTT |

Relative quantification of gene expression was performed using the $\Delta\Delta C_t$ method, with *pmp-3* serving as the housekeeping control gene [20], and mRNA from wild type worms cultured on vector serving as the normalization control. Primer pair amplification efficiencies ranged from 1.76 - 2.0 and were accounted for in our final analysis. Statistical significance was analyzed using a one-tailed Student's t-test.

**Extracellular Vesicle (EV) Isolation.** We followed the procedure of Russell and colleagues ([21] to isolate EVs from *C. elegans*. Specifically, 350,000 synchronous N2 (wild type) and 350,000 SLR117 [*isp-1(qm150)*] worms were cultured to young adulthood on 2% peptone-enriched NGM plates seeded with OP50 bacteria (15,000 worms per 10 cm plate, 20°C). These worms were seeded as arrested L1 larvae, and sourced from parents also cultured on 2% peptone-enriched NGM plates. Upon reaching young adulthood, animals were collected in S-Basal buffer, allowed to settle by gravity for 5 minutes, then washed five more times with 50 ml S-Basal. Worms were resuspended in 15 ml ice-cold 100 mM NaCl, after which an equal volume of 60% ice-cold sucrose was added and the tube mixed. Worms were then floated by centrifugation (1000 X G, 8 min., Beckman GS-6R with GH_3.8 swinging bucket rotor). Buoyant worms were collected and transferred into a fresh 50 ml conical tube, washed thrice with 50 ml S-Basal, then resuspended in S-Basal supplemented with 2.5 ug/ml cholesterol to a final density of 1 animal per µl (keeping the maximum volume in each 50 ml tube at 45 ml). Worm samples were then incubated at 20°C for 24 hours, with end-over-end rotation. At the end of the incubation period animals were removed by centrifugation (2500 X G, 10 min., Beckman

GS-6R with GH_3.8 swinging bucket rotor) and discarded. The EV-enriched supernatant was passed through a 0.2 µm filter and the flow through collected. Large lipid particles were pelleted at 18,000 X G for 30 min. at 4°C. The supernatant was decanted and then EVs concentrated via an Amicon Ultra-10K device (Ultra-15, Millipore Sigma cat # UFC901024). EVs were centrifuged at 2500 X G until the volume was reduced to 1.5 ml. Halt ™ Protease Inhibitor Cocktail (supplemented with EDTA) (Thermo Fisher Scientific, # 78430) was added to the concentrate (1X final) then the sample was passed over a 10 ml Sepharose CL-2B size exclusion column using S-Basal as the mobile phase. 15 x 1 ml elution fractions were collected and then pooled as follows: Fractions 2-6 (EV-enriched), 7-11 (few EVs, enriched in non-vesicle nanoparticles), and 12-16 (only non-vesicle nanoparticles). These three pooled samples were each further concentrated using an Amicon Ultra-10K device (as described above) to a final volume of 100 µl. All samples were utilized in downstream analyses within 72 hours of harvesting.

To determine whether the EV-enriched fractions from *isp-1(qm150)* mutants could alter the expression of stress reporter genes relative to EV-containing fractions from wild type worms, we exposed CL2166, SJ4100 and SLR115 reporter strains to EV fractions for the duration of their development. Specifically, the three Sepharose CL-2B pooled fractions described above were each diluted to 600 µl using S-Basal. Six mature *E. coli* (OP50) lawns were evenly overlaid with 100 µl of the relevant EV-containing fraction from N2 or SLR117 worms and allowed to air dry. These *E. coli* lawns were previously prepared by spotting a 200 µl aliquot of OP50 in fresh LB media ($OD_{600}$ 0.9) onto a 6 cm NGM agar plate and then allowing it to grow to maturity for three days. When the lawns had re-dried following addition of the EV-containing fractions, 60 synchronous eggs from the relevant reporter strain were transferred onto the lawn and allowed to develop into first day adults. Reporter gene expression was then quantified as described in the next section.

**Fluorescence Imaging and Quantification**. Worm images were captured using an Olympus DP71 CCD camera connected to an Olympus SZX12 or SZX16 fluorescence dissecting microscope. Third allele versions of ALX-1, CO1A2.4 and ISTR-1 encoding C-terminal GFP fusion proteins were cultured on undiluted *isp-1* feeding RNAi and GFP fluorescence recorded at the first day of adulthood. All worm images were quantified using ImageJ (NIH). Briefly, worm outlines were manually traced and mean pixel density computed (n = 20 worms per condition, unless stated otherwise). This measure served as the comparative metric between conditions. Samples were normalized to the relevant control condition and statistical significance was analyzed using Student's *t-test* (Excel, Microsoft), without further correction to p-values.

### SUPPLEMENTAL FIGURE LEGENDS

**Figure S1. Selection of a PMK-3 epitope for antibody generation.** CLUSTALX-generated sequence alignment of all predicted p38 MAPK splice variants of *C. elegans. Light yellow box:* Peptide sequence used for PMK-3 antibody production. *Grey boxes* on PMK-3 sequence indicate regions of exceptional sequence variation relative to other PMK isoforms. The sequence insertion shown at position 457 of PMK-3 (*red lower-case sequence*) represents the in-frame intron found in the RT-PCR-derived cDNA clone obtained by Berman and colleagues [22]. *Pink box:* Predicted TQY/TGY double phosphorylation motif of activation loop. *Orange* and *green boxes:* Residues in PMK-3 removed by the *ok169* and *tm745* deletions, respectively. *Red arrows:* PMK-3 exon/exon splice junctions. *Grey bars* under alignment highlight proportionally regions of homology. Table in *lower right* indicates estimated molecular weights of each protein isoform.

**Figure S2. Sensitivity of PMK-3 antisera.**

(**A**) *Upper panel*: Schematic of *pmk-3* genetic locus copied from **Fig. 1A** (main text). Position of the GFP fusion junction in the PMK-3 translational reporters of strains DA1750 and GJ3080 are marked (*long arrows). Lower panel:* Two promoters control expression of the *pmk-3* locus (*green* bars): a 1.5 kb sequence immediately upstream of the start codon of PMK-3(L) that extends 500 bp into the 3' end of the upstream *islo-1* gene [22]; and the promoter of *islo-1* itself, which controls expression of an operon comprised of *islo-1, pmk-3, pmk-2* and *pmk-1* (*lower panel). pmk-1* and *pmk-2* encode the only other p38 paralogs present in the *C. elegans* genome.

Neither PMK-1 nor PMK-2 can compensate for the mitochondrial retrograde response controlled by PMK-3 [3].

(**B**) Strain GJ3080 contains an extrachromosomal array encoding C-terminally tagged long- and short forms of PMK-3 (A). The array is co-marked with mCherry. Weak, nuclear GFP staining is detectable in all cells following integrated signal capture. Scale bar: 200 µm.

(**C**) Antisera targeting an 18 αα sequence unique to PMK-3 (**Fig. S1**) recognizes a weakly-expressed GFP-tagged form of PMK-3(L) within strain GJ3080. Arrows mark recombinant PMK-3. Asterisks mark cross-reactive bands present in control strain extracts [*pmk-3(tm745)* and GJ3080 worms that have lost the array].

(**D**) Movement of the extrachromosomal array in GJ3080 worms into the *pmk-3(tm745)* mutant background (strain SLR0166) shows GFP-tagged PMK-3 is functional since it rescues the knock-out allele, as indicated by intestinal induction of the P*tbb-6*::GFP reporter following exposure to *atp-3* feeding RNAi (1/10$^{th}$ strength). Worms containing the array are marked by constitutive mCherry expression. Scale bar: 200 µm.

(**E**) Strain DA1750 contains an extrachromosomal array encoding GFP fused in-frame to a 3 kb fragment of the *pmk-3* genomic locus that includes 1.5 kb of sequence immediately upstream of the start codon of PMK-3(L) (A). Reporter gene expression is dampened following exposure to pathogenic *Pseudomonas aeruginosa* (strain PA14) (+24 hours). Scale bar: 200 µm.

**Figure S3. Specificity of PMK-3 antisera.**

(**A**) No cross-reactivity of PMK-3 antisera toward PMK-1 and PMK-2. Worm of the listed genotype *(top, see also* ***Table S1****)* were cultured on *Escherichia coli* (strain OP50) or pathogenic *Pseudomonas aeruginosa* (strain PA14), then treated with either 1% Triton-X-100 (Tx-100) and then fractionated into soluble (*top left panel*) and insoluble components (*bottom left panel*), or simply treated with 1% SDS detergent (*top and bottom panels are identical)*. Extracts were analyzed by western blotting. Expected protein sizes were: PMK-1, 43.9 kDa; PMK-2a-d, 46.0, 45.5, 23.1 and 22.6 kDa respectively. Endogenous PMK-3(L) and (S) were not detected under these conditions (expected protein sizes: 54.9 and 35 kDa, respectively). Worms become colonized by PA14 and the strongly cross-reactive proteins in these lanes are all of bacterial origin (*refer to panel B*). The weak band at 37kDa in panel A is from OP50 carryover (*compare panel B, last lane*).

(**B**) Worms grown on PA14 pathogenic bacteria do not induce expression of PMK-3 in compensation for loss of *pmk-1*. Briefly, worms lacking *pmk-3* or *pmk-1* were cultured to adulthood on *E. coli* (OP50) then half exposed to pathogenic *P. aeruginosa* (PA14) for 24 hours. Whole-worm protein extracts (1% SDS) were prepared for western analysis using PMK-3 antisera. Control protein extracts were also prepared from each bacterial strain. Unlike *E. coli,* PA14 colonizes the gut, resulting in significant carry over of PA14 cells during worm extract preparation. PA14 bacterial cells contain a strongly reactive epitope(s) that is recognized by PMK-3 antisera. *E. coli* also contains a protein(s) this is recognized by the PMK-3 antisera, but it only becomes evident when bacterial cells are present in high quantities (last lane). Weak carry over of some of these *E. coli* proteins during worm extract preparation is evident in **Figs. S2C** as well as **Fig. S4**, but they do not interfere with our general conclusions.

**Figure S4. Raw data for western blot images shown in Fig. 1B.**

(**A**) Original anti-PMK-3 western blot corresponding to **Fig. 1B** of main text. All samples are whole-worm lysates (1% SDS) derived from the listed strain (refer to **Table S1** for full genotype information). Each strain contains a newly-constructed PMK-3 or DLK-1 recombinant isoform (*top*). All lanes contain equal amounts of protein. Double arrow highlights shift in migration of the long form of PMK-3 caused by EQE double point mutation.

(**B**) Fully independent replicate of analysis in (A). *Top panel:* 10 second α-PMK-3 immunoblot exposure; *middle panel:* 5-minute α-PMK-3 immunoblot exposure; *bottom panel;* stain of original SDS-PAGE gel after nitrocellulose transfer showing equality of protein loading. Bands corresponding to wild type and EQE mutated PMK-3(L) [and potentially PMK-3(S)] are marked with double arrows. *Asterisks* mark non-specific bands recognized by PMK-3 antisera. Most of these are likely bacterial in origin (compare **Fig. S3**, OP50 lane).

(**C, D**) Effect of *atp-3* RNAi (C) or *isp-1* RNAi (D) on PMK-3(L) abundance in transgenic P*vha-6*::PMK-3 worms. SLR153.1 and SLR153.2 are different strains containing independently-assembled P*vha-6*::PMK-3 arrays. *Arrows* mark PMK-3(L). *Asterisks* in (C) mark undigested proteins of *E. coli* origin (refer to **Fig. S3** for details).

**Figure S5. Structure-based sequence alignment of PMK-3(L) with other p38 MAPK isoforms.**

Homology models of PMK-3(L) (αα 110-459), PMK-2a (αα 33-380) and PMK-1 (αα 16-360) were generated using PDB structures 1R3C, 4LOQ and 5ETC, respectively, as described in **S1 Text**. The Magic Fit function of the program Swiss-Prot was then used to superpose all structures onto the PMK-3(L) model. This function utilizes a PAM 200 matrix to maximize structural (Cα) overlap between the most related sequences in each structure relative to PMK-3(L). The resulting structure-based sequence alignment is shown. *Orange box*: residues absent in

PMK-3(S); *green residues:* ATP-interacting residues; *magenta:* TQY/TGY activation loop phosphorylation site; *blue box:* epitope used for PMK-3 antisera generation.

**Fig. S6. Additional fluorescent images of *pmk-3(tm745)* knockout mutants expressing *pmk-3* transgenes under the control of the *rgef-1* neuronal promoter.**

(**A**) Corresponds to **Fig. 2D** of main text. Additional images of strain SLR0251 displaying strong neuronal mCherry expression but also leaky gut expression. Use of a lower *isp-1* feeding RNAi dose (1/10$^{th}$ strength) illustrates a positive correlation between P*rgef-1* leakage and P*tbb-6*::GFP expression in gut tissue (*lower row*). *Asterisk* in top row indicates worm that has lost the array. All *scale bars*: 250 μm.

(**B**) Construction of neuronal-expressed *pmk-3* transgenes. Constructs are comprised of a two-gene operon comprised of: (*left* to *right*) a neuron-specific promoter (*Prgef-1* or P*rab-3)*, *pmk-3* cDNA, splice leader sequence (SL2), mCherry reporter, *unc-54* 3'UTR. Strain names of worms carrying the relevant transgene are indicated. Boxes in pink indicate insertion of a decoy open reading frame (ORF) designed to reduce gut mis-expression (see **S1 text** for details).

**Figure S7. Additional survival analyses.** Survival statistics for all panels is provided in **Table S2**.

(**A**) Corresponds to **Fig. 3B** *(left panel)* of main text. A second, independently generated SLR0153 line (here named Line #1) restored full life extension to *pmk-3(tm745)* mutants relative to SLR0155 worms (compare with **Fig. 3C** *(left panel).*

In (B-D), worms were cultured on a pathogenic dose of ½ strength *atp-3* bacterial feeding RNAi from the time of hatching.

(**B**) Wild type worms (strain SLR115) are sick, small and show no significant increase in life span when cultured on ½ strength *atp-3* feeding RNAi, as previously reported [4]. *pmk-3(tm745)* knock-out mutants arrest as L3 larvae and die prematurely.

(**C**) Ubiquitous re-expression of PMK-3(L) overcomes the L3 arrest phenotype induced by ½ strength *atp-3* feeding RNAi in *pmk-3(tm745)* null mutants. Rescue is partial since not all animals exit L3 arrest and accordingly survival does not return to wild type levels. Partial rescue may reflect incomplete penetrance of the array across cells of individual worms (mosaicism).

(**D**) Intestinal re-expression of PMK-3(L) in *pmk-3(tm745)* mutants enhances survival in response to ½ strength *atp-3* feeding RNAi to the same extent as ubiquitously re-expressed PMK-3 (compare solid green lines in panels B and C). In both panels B and C, worms that have lost the array (-) are included to control for potential strain background effects.

**Figure S8**. ***Myc*-tagged DLK-1b(EE) expressed in *dlk-1(tm4024)* knockout mutants becomes activated only in the presence of ETC stress.**

(**A**) Schematic of *dlk-1* genetic loci. mRNA splice variants are shown. Size of encoded protein products listed on *right* (αα - amino acids). Genomic DNA deleted in *dlk-1(tm4024)* mutants is shown with a *red* bar. Amino acids of DLK-1 altered by point mutation in this study are colored *red*. Exons encoding relevant protein domains are boxed. Position of the myc epitope insertion used in strain SLR0182 is marked (*long arrows)*.

(**B**) Recombinant *dlk-1b(EE)* transgene generation, with and without a *myc* epitope tag. Both constructs are comprised of a two-gene operon assembled as follows: (*left* to *right*) promoter (P*vha-6)*, *dlk-1b* cDNA containing EE point mutation in the SDGLSD sequence (A), splice leader sequence (SL2), mCherry reporter, *unc-54* 3'UTR. Strain names of worms carrying the relevant transgene are indicated, along with a summary of the effect of each transgene on *Ptbb-6::GFP* reporter expression under the listed conditions.

(**C**) Intestinal expression of P*tbb-6*::GFP, following mitochondrial ETC disruption by *isp-1* feeding RNAi, can be restored in *dlk-1(tm4024)* mutants following gut-specific (*vha*-6 promoter) re-expression of DLK-1b(EE)*myc*. Control strains SLR115 (wild type) and SLR0176 [*dlk-1(tm4024)* knockout] highlight the degree of rescue. All *scale bars*: 500 µm (note magnification has been increased for strain SLR0182).

Why untagged DLK-1b(EE) expressed in wild type worms results in constitutive P*tbb-6::GFP* induction [see (B) and **Fig. 4B,** main text], but *myc*-tagged DLK-1b(EE) expressed in *dlk-1(tm4024)* knockout mutants becomes activated only in the presence of ETC stress, is unclear. The *myc* epitope may disrupt the activity or stability of mutant DLK-1b(EE), or other DLK-1 isoforms (A) might be required to heterodimerize with DLK-1b(EE) and induce its constitutive activity in the absence of ETC stress.

**Figure S9. No alteration in PMK-3::GFP fluorescence following mitochondrial ETC disruption in DA1750 worms.**

Strain DA1750 contains an extrachromosomal array encoding GFP fused in-frame to a 3 kb fragment of the PMK-3 genomic locus that includes 1.5 kb of sequence immediately upstream of the start codon of PMK-3(L) (see **Fig. S2A**). Following growth on bacterial feeding RNAi targeting *isp-1* (1/10$^{th}$ strength), reporter gene expression in DA1750 worms remains unaltered relative to vector-only treated animals (V*ec.*). Control SLR115 worms, containing the P*tbb-6*::GFP reporter, show strong reporter induction following growth on *isp-1* RNAi. Scale bar: 200 µm.

**Figure S10. Knockdown of *istr-1* by RNAi does not inhibit the efficacy of the RNAi machinery.** Wild type worms (n=20 per condition) were fed from the arrested L1 stage onward, defined mixes of bacteria containing RNAi targeting *skn-1* and either *istr-1* or no other target (these bacteria contained empty vector, *Vec*). Shown is the percentage of F1 progeny that hatch at each *skn-1* concentration for each test RNAi. Curves were analyzed using a log-rank test. *istr-1* RNAi showed no capacity beyond the diluting effect of vector control to reduce the potency of *skn-1* RNAi (*p-value* 0.8304).

**Figure S11. Extracorporeal EVs isolated from starved *isp-1(qm150)* mutants do not activate ETC stress reporter genes.**

Extracellular vesicles isolated from wild type (N2) or mutant *isp-1(qm150)* worms do not induce the expression of mitochondrial retrograde-response reporter genes when supplemented to the growth media of unstressed reporter gene-containing animals. *Ptbb-6::GFP, Pgst-4::GFP* and *Phsp-6::GFP* are sensitive to the PMK-3, SKN-1 and ATFS-1 retrograde responses, respectively. GFP fluorescence was measured for ~60 animals following full development on EV populations. GFP fluorescence was normalized to animals treated with buffer alone. Each dot represents the GFP signal from one test sample, averaged across 20 worms (n = 3-4 independent replicates). No differential effect of EVs on reporter signaling was found (ANOVA, $p>0.5$).

### SUPPLEMENTAL TABLE LEGENDS

**Table S1. Worm strains.** List of strains employed in current study, and their associated genotype information.

**Table S2. Lifespan Analyses -** Description of all lifespan experiments and their statistical significance.

### SUPPLEMENTAL REFERENCES

1. van der Vaart A, Rademakers S, Jansen G. DLK-1/p38 MAP Kinase Signaling Controls Cilium Length by Regulating RAB-5 Mediated Endocytosis in Caenorhabditis elegans. PLoS Genet. 2015;11(12):e1005733. doi: 10.1371/journal.pgen.1005733. PubMed PMID: 26657059; PubMed Central PMCID: PMCPMC4686109.
2. Sarov M, Schneider S, Pozniakovski A, Roguev A, Ernst S, Zhang Y, et al. A recombineering pipeline for functional genomics applied to Caenorhabditis elegans. Nature methods. 2006;3(10):839-44. Epub 2006/09/23. doi: 10.1038/nmeth933. PubMed PMID: 16990816.
3. Munkacsy E, Khan MH, Lane RK, Borror MB, Park JH, Bokov AF, et al. DLK-1, SEK-3 and PMK-3 Are Required for the Life Extension Induced by Mitochondrial Bioenergetic Disruption in C. elegans. PLoS Genet. 2016;12(7):e1006133. doi: 10.1371/journal.pgen.1006133. PubMed PMID: 27420916; PubMed Central PMCID: PMCPMC4946786.
4. Rea SL, Ventura N, Johnson TE. Relationship Between Mitochondrial Electron Transport Chain Dysfunction, Development, and Life Extension in Caenorhabditis elegans. PLoS Biol. 2007;5(10):e259. Epub 2007/10/05. doi: 06-PLBI-RA-2325 [pii]

10.1371/journal.pbio.0050259 [doi]. PubMed PMID: 17914900; PubMed Central PMCID: PMCPMC1994989.
5. Fraser AG, Kamath RS, Zipperlen P, Martinez-Campos M, Sohrmann M, Ahringer J. Functional genomic analysis of C. elegans chromosome I by systematic RNA interference. Nature. 2000;408(6810):325-30. PubMed PMID: 11099033.
6. Bhaskaran S, Butler JA, Becerra S, Fassio V, Girotti M, Rea SL. Breaking Caenorhabditis elegans the easy way using the Balch homogenizer: an old tool for a new application. Analytical Biochemistry. 2011;413(2):123-32. Epub 2011/03/01. doi: 10.1016/j.ab.2011.02.029. PubMed PMID: 21354098.
7. Pagano DJ, Kingston ER, Kim DH. Tissue expression pattern of PMK-2 p38 MAPK is established by the miR-58 family in C. elegans. PLoS Genet. 2015;11(2):e1004997. doi: 10.1371/journal.pgen.1004997. PubMed PMID: 25671546; PubMed Central PMCID: PMCPMC4335502.
8. Kim DH, Feinbaum R, Alloing G, Emerson FE, Garsin DA, Inoue H, et al. A Conserved p38 MAP Kinase Pathway in Caenorhabditis elegans Innate Immunity

10.1126/science.1073759. Science. 2002;297(5581):623-6.
9. Nielsen M, Lundegaard C, Lund O, Petersen TN. CPHmodels-3.0--remote homology modeling using structure-guided sequence profiles. Nucleic acids research. 2010;38(Web Server issue):W576-81. Epub 2010/06/15. doi: 10.1093/nar/gkq535. PubMed PMID: 20542909; PubMed Central PMCID: PMC2896139.
10. Patel SB, Cameron PM, Frantz-Wattley B, O'Neill E, Becker JW, Scapin G. Lattice stabilization and enhanced diffraction in human p38 alpha crystals by protein engineering. Biochim Biophys Acta. 2004;1696(1):67-73. PubMed PMID: 14726206.
11. Khan MH, Ligon M, Hussey LR, Hufnal B, Farber R, 2nd, Munkacsy E, et al. TAF-4 is required for the life extension of isp-1, clk-1 and tpk-1 Mit mutants. Aging (Albany NY). 2013;5(10):741-58. Epub 2013/10/11. doi: 10.18632/aging.100604. PubMed PMID: 24107417; PubMed Central PMCID: PMCPMC3838777.
12. Sleumer MC, Bilenky M, He A, Robertson G, Thiessen N, Jones SJ. Caenorhabditis elegans cisRED: a catalogue of conserved genomic elements. Nucleic Acids Res. 2009;37(4):1323-34. doi: 10.1093/nar/gkn1041. PubMed PMID: 19151087; PubMed Central PMCID: PMCPMC2651782.

13. Huang da W, Sherman BT, Lempicki RA. Systematic and integrative analysis of large gene lists using DAVID bioinformatics resources. Nat Protoc. 2009;4(1):44-57. doi: 10.1038/nprot.2008.211. PubMed PMID: 19131956.
14. Fabris F, Freitas AA. New KEGG pathway-based interpretable features for classifying ageing-related mouse proteins. Bioinformatics. 2016. doi: 10.1093/bioinformatics/btw363. PubMed PMID: 27318209.
15. Kanehisa M, Goto S, Kawashima S, Okuno Y, Hattori M. The KEGG resource for deciphering the genome. Nucl Acids Res. 2004;32(suppl_1):D277-80. doi: 10.1093/nar/gkh063.
16. Edgar R, Domrachev M, Lash AE. Gene Expression Omnibus: NCBI gene expression and hybridization array data repository. Nucleic Acids Res. 2002;30(1):207-10. PubMed PMID: 11752295; PubMed Central PMCID: PMCPMC99122.
17. Barrett T, Wilhite SE, Ledoux P, Evangelista C, Kim IF, Tomashevsky M, et al. NCBI GEO: archive for functional genomics data sets--update. Nucleic Acids Res. 2013;41(Database issue):D991-5. doi: 10.1093/nar/gks1193. PubMed PMID: 23193258; PubMed Central PMCID: PMCPMC3531084.
18. Nargund AM, Pellegrino MW, Fiorese CJ, Baker BM, Haynes CM. Mitochondrial import efficiency of ATFS-1 regulates mitochondrial UPR activation. Science. 2012;337(6094):587-90. Epub 2012/06/16. doi: 10.1126/science.1223560. PubMed PMID: 22700657.
19. Wood WB, editor. The Nematode Caenorhabditis elegans. New York: Cold Spring Harbor Laboratory; 1988.
20. Hoogewijs D, Houthoofd K, Matthijssens F, Vandesompele J, Vanfleteren JR. Selection and validation of a set of reliable reference genes for quantitative sod gene expression analysis in C. elegans. BMC Mol Biol. 2008;9:9. PubMed PMID: 18211699.
21. Russell JC, Merrihew GE, Robbins JE, Postupna N, Kim T-K, Golubeva A, et al. Isolation and characterization of extracellular vesicles from Caenorhabditis elegans for multi-omic analysis. 2018;bioRxiv doi: bioRxiv 476226; doi: https://doi.org/10.1101/476226
22. Berman K, McKay J, Avery L, Cobb M. Isolation and characterization of pmk-(1-3): three p38 homologs in Caenorhabditis elegans. Mol Cell Biol Res Commun. 2001;4(6):337-44. PubMed PMID: 11703092.
